# Mapping the planet’s critical natural assets

**DOI:** 10.1101/2020.11.08.361014

**Authors:** Rebecca Chaplin-Kramer, Rachel A Neugarten, Richard P Sharp, Pamela M Collins, Stephen Polasky, David Hole, Richard Schuster, Matthew Strimas-Mackey, Mark Mulligan, Carter Brandon, Sandra Diaz, Etienne Fluet-Chouinard, LJ Gorenflo, Justin A Johnson, Christina M Kennedy, Patrick W Keys, Kate Longley-Wood, Peter B McIntyre, Monica Noon, Unai Pascual, Catherine Reidy Liermann, Patrick R Roehrdanz, Guido Schmidt-Traub, M Rebecca Shaw, Mark Spalding, Will R Turner, Arnout van Soesbergen, Reg A Watson

**Author notes:** Correspondence to: Rebecca Chaplin-Kramer, **Email:**.

## Abstract

Sustaining the organisms, ecosystems, and processes that underpin human well-being is necessary to achieve sustainable development. Here we identify critical natural assets, natural and semi-natural ecosystems that provide 90% of the total current magnitude of 14 types of nature’s contributions to people (NCP). Critical natural assets for maintaining local-scale NCP (12 of the 14 NCP mapped) comprise 30% of total global land area and 24% of national territorial waters, while 44% of land area is required for maintaining all NCP (including those that accrue at the global scale, carbon storage and moisture recycling). At least 87% of the world’s population lives in the areas benefiting from critical natural assets for local-scale NCP, while only 16% lives on the lands containing these assets. Critical natural assets also overlap substantially with areas important for biodiversity (covering area requirements for 73% of birds and 66% of mammals) and cultural diversity (representing 96% of global Indigenous and non-migrant languages). Many of the NCP mapped here are left out of international agreements focused on conserving species or mitigating climate change, yet this analysis shows that explicitly prioritizing critical natural assets for NCP could simultaneously advance development, climate, and conservation goals. Crafting policy and investment strategies that protect critical natural assets is essential for sustaining human well-being and securing Earth’s life support systems.

---

Human actions are rapidly transforming the planet, driving losses of nature at an unprecedented rate that negatively impacts societies and economies – from accelerating climate change to increasing zoonotic pandemic risk^1,2^. Recognizing the accelerating severity of the environmental crisis, the global community committed to Sustainable Development Goals (SDGs) and the Paris Agreement on climate change in 2015. In 2022, the UN Convention on Biological Diversity (CBD) will adopt new targets for conserving, restoring and sustainably managing multiple dimensions of biodiversity, including nature’s contributions to people (NCP)^3^. Collectively, these three policy frameworks will shape the sustainable development agenda for the next decade. All three depend heavily on safeguarding natural assets^1,4^, the living components of our lands and waters. For instance, restoring and ending conversion and degradation of forests, wetlands and peatlands could sequester 9 Gt CO_2_ per year by 2050^5^. While ambitious new targets to protect species and ecosystems have been proposed, including “half earth” (conserving half the earth’s area for nature)^6^ and “30 by 30” (30% protected by 2030)^7^, these targets have been criticized for insufficiently accounting for the needs of people, including Indigenous and local communities^8^. Protecting critical natural assets, however, shows directly how conservation contributes to human well-being.

Despite the urgency of safeguarding natural assets, we still have limited understanding of the spatial extent of ecosystems providing essential benefits to humanity^9^. Leveraging recent advances in scientific understanding, data availability (Extended Data Tables 1-2) and computational power, we undertake an global analysis of 14 NCP (Extended Data Fig. 1), the most comprehensive set of NCP mapped globally to date^10,11^. Twelve of these NCP deliver primarily local benefits (though some subsequently enter global supply chains), including contributions to the provision of food, energy, and raw materials; the regulation of water quality and disaster risk; and recreational activities (Fig. 1a). We prioritize these 12 “local” NCP at the country level to identify areas needed to maintain consistent provision within each country. In contrast, we prioritize at the global scale for two NCP related to climate regulation (terrestrial ecosystem carbon storage and vegetation-regulated atmospheric moisture recycling), whose benefits accrue at continental scales or globally. Through multi-objective spatial optimization, we map the location of the planet’s *critical natural assets*, defined as the natural and semi-natural ecosystems providing 90% of current levels of each NCP. Beyond this target there are diminishing returns, with disproportionately more natural area required to reach incrementally higher levels of NCP (Fig. 1b). We focus on natural ecosystems (Extended Data Table 3), excluding developed lands (croplands and urban areas), to provide insights for conservation priorities relevant to the CBD; global priorities for restoration^12^ or for management of developed areas^13^ are beyond the scope of this first effort to map critical natural assets. Our analysis reveals three key findings about critical natural assets: 1) their extent and location; 2) the number of people benefitting from, and living within, these areas; and 3) the degree of overlap between critical natural assets delivering local benefits and those delivering global benefits, as well as other global priorities for the CBD (biodiversity and cultural diversity).

**Figure 1.**
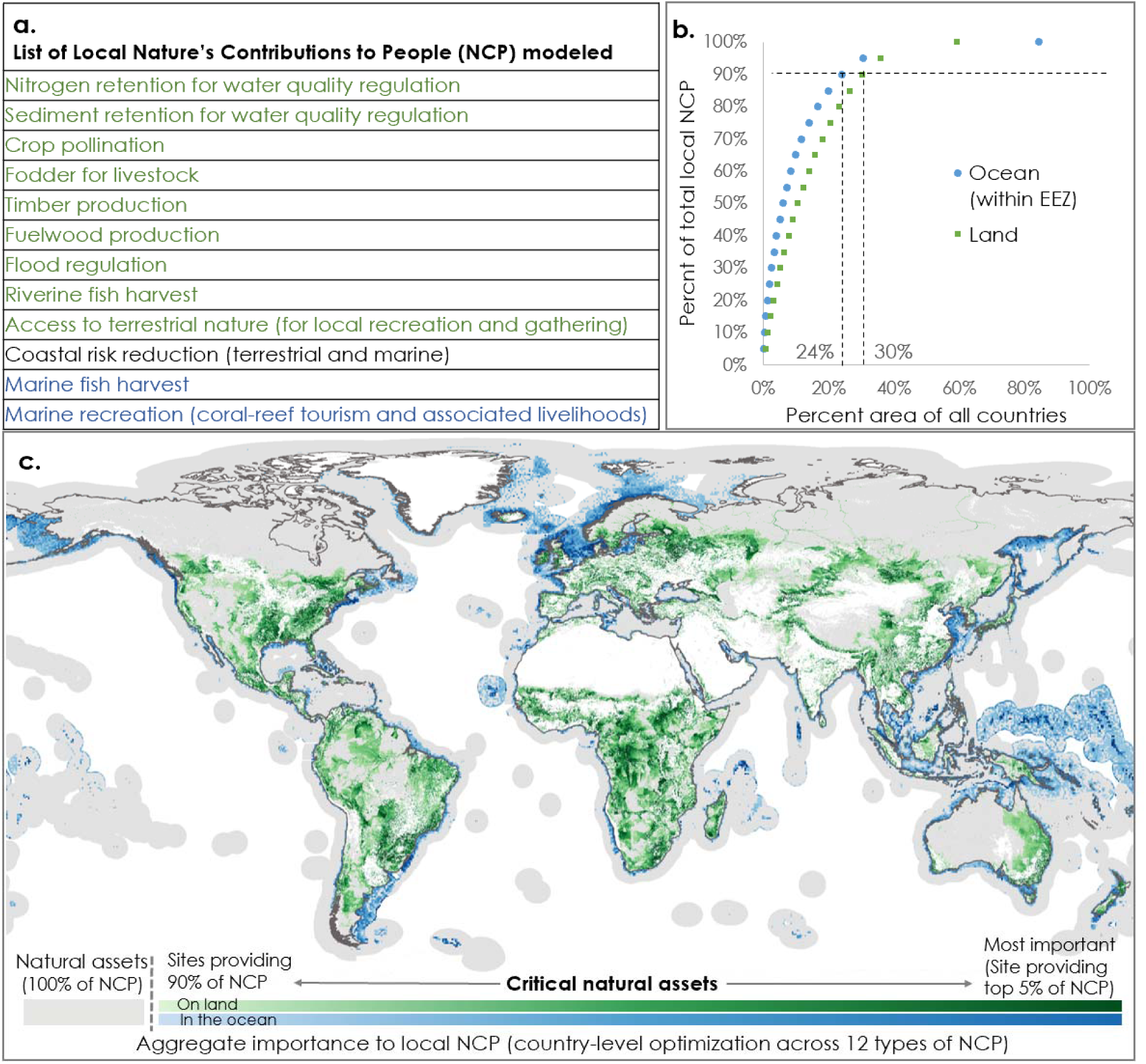
Critical natural assets, defined as the natural and semi-natural terrestrial and aquatic ecosystems required to maintain 12 of nature’s “local” contributions to people (local NCP) on land (green) and in the ocean (blue). (a) The 12 local NCP analyzed (i.e., not including global NCP, shown in Fig. S4). (b) The NCP accumulation curve, reflecting the total area required to maintain target levels of all NCPs in every country, with dotted lines denoting the area of critical natural assets (90% of NCP in 30% of land area and 24% of EEZ area). Areas selected by optimization within each country are aggregated across all countries to create a single global accumulation curve; see Table S1 for area requirements in individual countries. (c) Map of critical natural assets, with darker shades connoting critical natural assets that are associated with higher levels of aggregated NCP. Grey areas show the extent of remaining natural assets not designated “critical” but included in this analysis; white areas (cropland, urban and bare areas, ice and snow, and ocean areas outside the EEZ) were excluded from the optimization.

Critical natural assets that provide 90% of current levels of each of the 12 local NCP (Fig. 1a) occupy only 30% (41 million km^2^) of total land area (excluding Antarctica) and 24% (34 million km^2^) of marine Exclusive Economic Zones, reflecting the steep slope of the aggregate NCP accumulation curve (Fig. 1b). Despite this modest proportion of global land area, the shares of countries’ land areas that are designated as critical are highly variable. The 20 largest countries require only 24% of their land area, on average, to maintain 90% of current levels of NCP, while smaller countries (10,000 - 1.5 million km^2^) require on average 40% of their land area (SI Table 1). This high variability in the NCP-area relationship is primarily driven by the proportion of countries’ land areas made up by natural assets (i.e., excluding barren, ice and snow, and developed lands) but even when this is accounted for, there are outliers (Extended Data Fig. 2). Outliers may be due to spatial patterns in human population density (for example, countries with dense population centers and vast expanses with few people, such as Canada and Russia, require far less area to achieve NCP targets) or large ecosystem heterogeneity (if greater ecosystem diversity yields higher levels of diverse NCP in a smaller proportion of area, which may explain patterns in Chile and Australia).

Critical natural asset “hotspots” (highest-contributing areas, denoted by the darkest blue or green shades in Fig. 1c) often coincide with diverse, relatively intact natural areas near or upstream from large numbers of people. Many hotspots coincide with areas of greatest spatial congruence between multiple NCP (Extended Data Fig. 3). Spatially correlated pairs of “local” NCP (Extended Data Table 4) include those related to water (flood risk reduction with nitrogen retention, nitrogen with sediment retention); forest products (timber and fuelwood); and those occurring closer to human-modified habitats (pollination with nature access and with nitrogen retention). Coastal risk reduction, forage production for grazing, and riverine fish harvest are the most spatially distinct from other NCP. In the marine realm, there is substantial overlap between fisheries and the other two NCP, but not between coastal risk reduction and reef tourism (which each have much smaller critical areas than exist for fisheries).

We estimate that ∼87% of the global population, 6.4 billion people, benefit directly from at least one of the 12 local NCP provided by critical natural assets, while only 16% live on the lands providing these benefits (and they may also benefit; Fig 2a). To quantify the number of beneficiaries of critical natural assets, we spatially delineate their benefiting areas (which varies based on NCP: e.g., areas downstream, within the floodplain, in low-lying areas near the coast, or accessible by a short travel). Our optimization selects for the provision of 90% of the current value of each NCP, but it is not guaranteed that 90% of the people in the world would benefit (since it does not include considerations for redundancy in adjacent pixels and therefore many of the areas selected benefit the same populations), so it is interesting that nearly that many do. While this estimate of “local” beneficiaries likely underestimates the total number of people benefiting because it includes only NCP for which beneficiaries can be spatially delineated to avoid double-counting, it is striking that the vast majority, 6.1 billion people, live within one hour’s travel (by road, rail, boat or foot, taking the fastest path^14^) of critical natural assets, and more than half of the world’s population lives downstream of these areas (Fig. 2b). Material NCP are often delivered locally but many also enter global supply chains, making it difficult to delineate beneficiaries spatially for these NCP. However, past studies have calculated that globally more than 54 million people benefit directly from the timber industry^15^,157 million from riverine fisheries^16^, 565 million from marine fisheries^17^,1.3 billion from livestock grazing^18^, and across the tropics alone 2.7 billion are estimated to be dependent on nature for one or more basic needs^19^.

**Figure 2.**
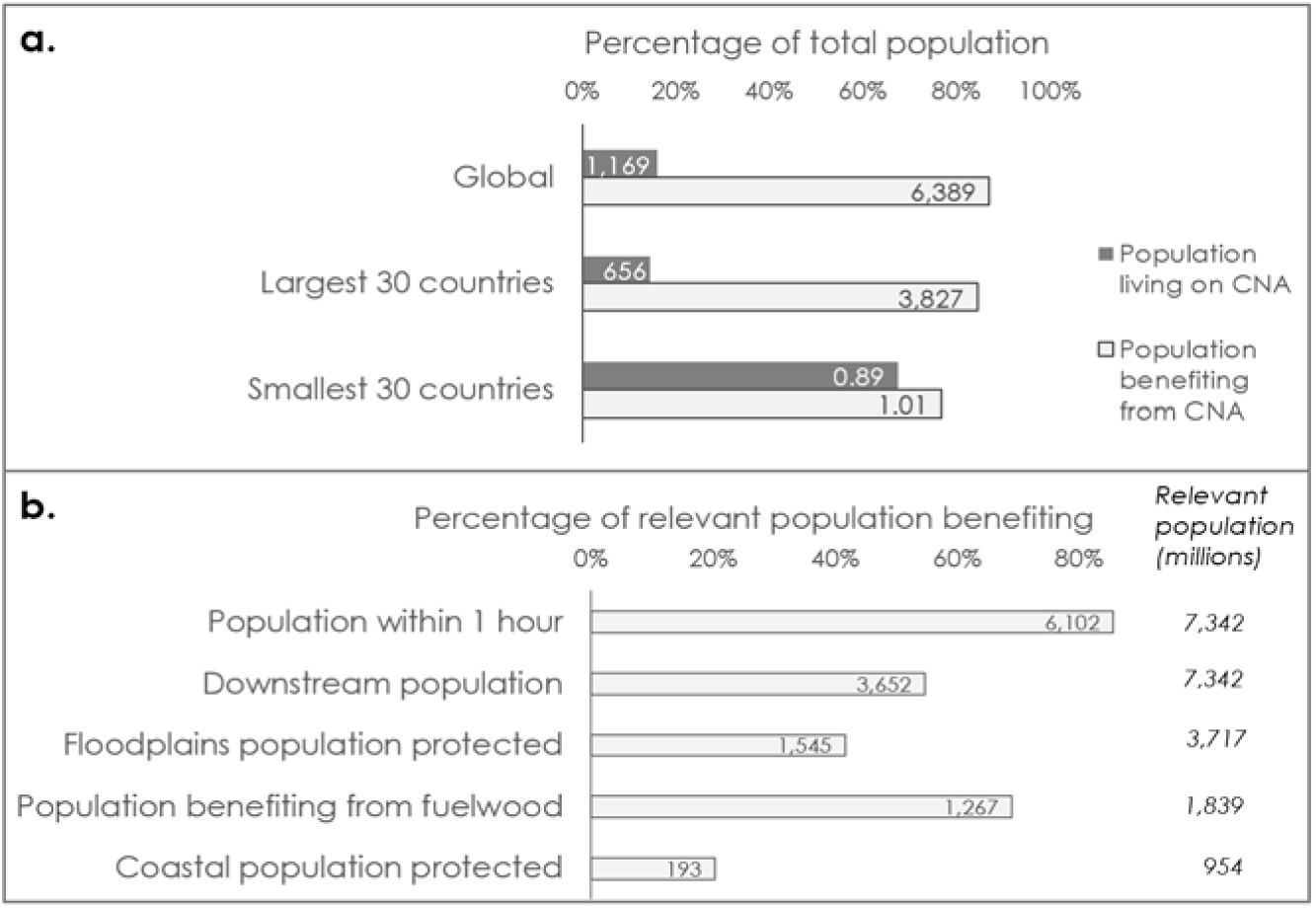
People benefiting from and living on critical natural assets (CNA). “Local” beneficiaries were calculated through the intersection of areas benefiting from different NCP, to avoid double-counting people in areas of overlap; only those NCP for which beneficiaries could be spatially delineated were included (i.e., not material NCP that enter global supply chains: fisheries, timber, livestock or crop pollination). Bars show percentages of total population globally and for large and small countries (a) or the percent of relevant population globally (b). Numbers inset in bars show millions of people comprising that percentage. Numbers to the right of bars in (b) show total relevant population (in millions of people, equivalent to total global population from Landscan 2017 for population within 1 hour’s travel or downstream, but limited to the total population living within 10km of floodplains or along coastlines <10m above mean sea level for floodplain and coastal populations protected, respectively, and to rural poor populations for fuelwood).

Nearly all countries have a large percentage (>80%) of their populations benefitting from critical natural assets, but small countries have much larger proportions of their populations living within the footprint of critical natural assets than do large countries (Fig. 2a; SI Table 2). When people live in these areas, and especially when current levels of use of natural assets are unsustainable, incentives or compensation may be needed to maintain the benefits these assets provide. While protected areas are an important conservation strategy, critical natural assets for NCP should not be protected using designations that restrict human access and use, or they could cease to provide many of the values that make them so critical. Other area-based conservation measures such as Indigenous and local governance, Payments for Ecosystem Services programs, and sustainable use of land- and seascapes can all contribute to maintaining critical flows of NCP in natural and semi-natural ecosystems.

Unlike the 12 local NCP prioritized here at national scales, certain benefits of ecosystems accrue continentally or globally. We therefore optimize two additional NCP at a global scale: vulnerable terrestrial ecosystem carbon storage (i.e., the proportion of total ecosystem carbon lost in a typical disturbance event^20^, hereafter “ecosystem carbon”) and vegetation-regulated atmospheric moisture recycling (the supply of atmospheric moisture and precipitation sustained by plant life^21^, hereafter “moisture recycling”). Over 80% of the natural asset locations identified as critical for the 12 local NCP are also critical for the two global NCP (Fig. 3). The spatial overlap between critical natural assets for local and global NCP comprises 24% of land area, with an additional 14% of land area critical for global NCP that is not considered critical for local NCP (Extended Data Fig. 4). Together, critical natural assets for securing both local and global NCP require 44% of total global land area. When each NCP is optimized individually (carbon and moisture NCP at the global scale; the other 12 at the country scale), the overlap between carbon or moisture NCP and the other NCP exceeds 50% for all terrestrial (and freshwater) NCP except coastal risk reduction (which overlaps only 36% with ecosystem carbon, 5% with moisture recycling).

**Figure 3.**
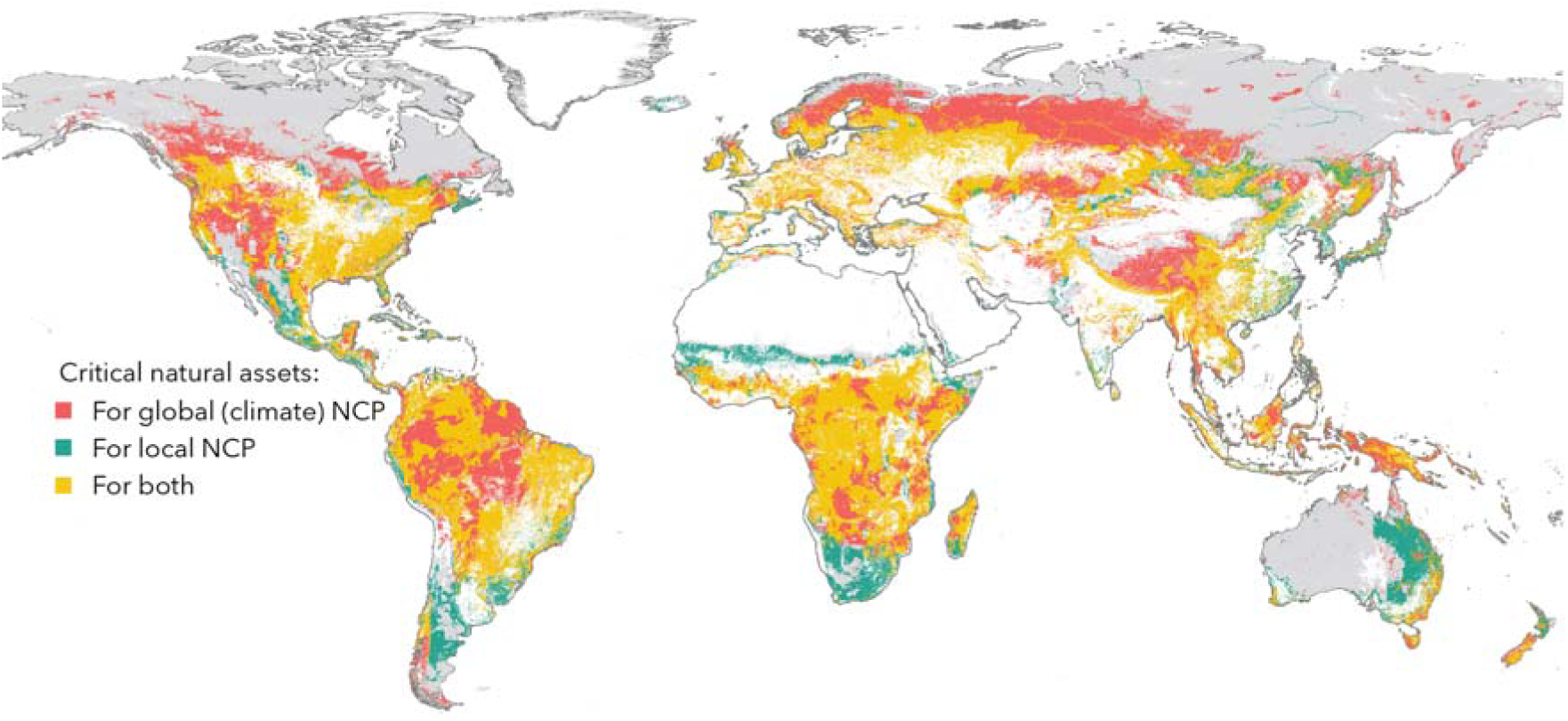
Spatial overlaps between critical natural assets for local and global NCP. Red and teal denote where critical natural assets for global climate NCP (providing 90% of ecosystem carbon and moisture recycling globally) or for local NCP (providing 90% of the 12 NCP listed in Fig 1), respectively, but not both, occur; gold shows areas where the two overlap (24% of the total area). Together, local and global critical natural assets comprise 44% of total global land area (excluding Antarctica). Grey areas show natural assets not defined as “critical” by this analysis, though still likely providing some values to certain populations. White areas were excluded from the optimization.

Synergies can also be found between NCP and biodiversity and cultural diversity. Critical natural assets for local NCP at national levels overlap with part or all of the Area of Habitat (AOH, mapped based on species range maps, habitat preferences and elevation^22^) for 60% of 28,177 terrestrial vertebrates (SI Table 3). Birds (73%) and mammals (66%) are better represented than reptiles and amphibians (44%). Only 34% of endemic vertebrate species overlap with critical natural assets for local NCP. Cultural diversity (proxied by linguistic diversity) has far higher overlaps with critical natural assets than does biodiversity; these areas intersect 96% of global Indigenous and non-migrant languages^23^ (SI Table 4). Despite the larger land area required for maintaining the global NCP compared to local NCP, global NCP priority areas overlap with only 2% more species (60% of species AOH) and with slightly fewer languages (92%). The degree to which languages are represented in association with critical natural assets is consistent across most countries, even at the high end of language diversity (countries containing >100 Indigenous and non-migrant languages, such as Indonesia, Nigeria, and India). This high correspondence provides further support for the link between and protection of critical natural assets in which Indigenous cultures benefit from and help maintain critical natural assets.

Although these 14 NCP are not comprehensive of the myriad ways nature benefits people^1^, they capture, spatially and thematically, many elements explicitly mentioned in the First Draft of the CBD’s post-2020 Global Biodiversity Framework (GBF)^24^: food security, water security, protection from hazards and extreme events, livelihoods, and access to green and blue spaces. Our emphasis here is to highlight the contributions of natural and semi-natural ecosystems to human well-being, specifically contributions that are often overlooked in mainstream policy. For example, considerations for global food security often include only crop production rather than nature’s contributions to it via pollination or vegetation-mediated precipitation, or livestock production without partitioning out the contribution of grasslands from more intensified feed production. Our synthesis of these 14 NCP at the global scale complements recent similar efforts at regional scales^25^ and represents a substantial advance beyond other global prioritizations that include NCP limited to ecosystem carbon stocks, fresh water, and marine fisheries^26–28^, though still falls short of including all important contributions of nature such as relational values of nature^29^. This same multi-NCP optimization approach could accommodate additional NCP as spatial data become available at sufficient resolution and appropriate scales.

There is uncertainty in the identification of critical natural assets related to: 1) the omission of many NCP that were not able to be mapped, and 2) model error in the individual NCP that we were able to include. Regarding this first source of uncertainty, further analyses indicate that results are fairly robust to inclusion of additional NCP. Dropping one of the 12 local NCP at a time results in <1-3% change in the total global land area required to maintain 90% of current levels of these NCP (Extended Data Table 5) and a high degree of spatial agreement. In fact, 62% of the total area on land is shared by all optimization solutions, and 97% of the area is included in 11 of the 12 solutions; similar values are found across most countries (SI Table 1). Regarding the second source of uncertainty, we acknowledge that all of the NCP models have errors and consistent global-scale modeling will miss details important for certain specific locations. Validation of NCP is particularly difficult given there are no direct measurements for many NCP with assessment reliant on remotely sensed proxies. We utilize the best available global modeling approaches and data, most of which have been validated in at least some locations^16,20,30–37^. As availability of global models for many of these NCP increases, future work should move toward ensemble modeling approaches, which have been shown to increase accuracy and reduce uncertainty compared to individual model outputs.^38,39^

Data and modeling gaps prevented a broader exploration of issues relevant to the ecological supply of NCP. Although results presented here suggest that nationally prioritized areas for local NCP can deliver on global priorities in many regions, they also highlight a need for integrated modeling to represent interactions between different NCP. For example, atmospheric moisture evapotranspired by Amazonian forests falls as rain in other parts of South America, supporting ecosystems that provide food, fuel, and other benefits^21^. Further work is needed to move beyond the spatial overlaps explored here toward understanding functional interdependencies between NCP. We also acknowledge that urban and cropland systems were omitted from this analysis due to data and modeling limitations that would fail to adequately capture the NCP supported by different land use types and land management practices within those systems. As data and modeling gaps are filled, future assessment of critical natural assets should consider possible gains from restoring and sustainably managing human-dominated systems^12,13^ and how these different conservation strategies can complement one another.

There are also, perhaps even more pronounced, data and modeling gaps to fill on the social side of NCP. In the NCP modeled here, we represent realized benefits of natural assets—weighted by beneficiary population when feasible—but this understates the range of ways in which natural assets directly and indirectly contribute to people’s wellbeing. Limited socio-economic data and lack of reliable models linking NCP to wellbeing indicators preclude more precise valuation of most NCP at the global scale. Additional insight could be gained from mapping critical natural assets that support the most vulnerable or dependent^19^ people, including Indigenous Peoples whose livelihoods and cultural identities directly depend on nature (and indeed overlap substantially with critical natural assets, based on our estimates of Indigenous language diversity on these lands; SI Table 4), and the poor who may lack access to anthropogenic substitutes for NCP (see also Philosophical Considerations in SI Methods). Recent progress in linking ecological modeling with integrated assessment modeling and general equilibrium economic modeling^40^ shows great promise for assessing the benefits of critical natural assets to society and the global economy. Such efforts could also reveal telecoupling of critical natural assets arising from transboundary flows between countries such as via international trade^41^.

Identifying critical natural assets could enable national and global leaders to prioritize the conservation of a wide range of NCP. We find it encouraging that securing 90% of the NCP mapped here is feasible with an area comparable to other proposed conservation targets^6,7,28,42^. Global analyses such as this can set a broader context for local decisions including understanding of distant connections that extend outside a country’s borders, provide rapid information for global actors such as CBD signatories and NGOs working on conservation priorities across many countries, and supplement gaps in local information while it is still being generated.^43^ We emphasize the value of the approach developed here more than the maps or data; this approach can be adapted and refined at national or local scales, the scales at which policy implementation occurs, with the best available data and complemented by input from local experts and diverse stakeholders, to improve accuracy and public legitimacy^44,45^ and to ensure human rights and diverse human relationships with nature are safeguarded. Moreover, creation and use of spatially-explicit information allows for a focus on ecosystem quality over quantity, helping to avoid potentially perverse outcomes of area targets for conservation.^46^ This approach for identifying critical natural assets is a vital step forward in empowering actors at all levels to make decisions that benefit both nature and people.

## Methods

### Modeling Nature’s Contributions to People (NCP)

The 14 NCP in this analysis (Extended Data Fig. 1) were chosen to span development and climate goals, and to be mappable with spatially explicit data representing the period 2000-2020. We use European Space Agency (ESA) 2015, for land cover, Landscan 2017 for population^47^ (these were the most current data available at the time we began our analysis). We focus on “nature’s contributions” to key benefits of interest (e.g., security in food, water, hazards, materials, culture), meaning we partition out the role of natural and semi-natural ecosystems in producing those benefits. For food security, we include the contributions of pollination to crop production, vegetation-mediated atmospheric moisture recycling to crop and livestock production (included as a global NCP), grassland fodder production to livestock production, and wild riverine and marine fisheries. For water security, we include the contributions of water quality regulation, via sediment retention and nutrient retention, but not water yield since the role of ecosystems in determining the quantity of water is minimal (other than by evapotranspiration which is already captured in the vegetation-mediated moisture recycling, and regulation of timing of flows which is captured in flood risk reduction). For security of protection from natural hazards, we include flood risk reduction and coastal risk reduction. For materials, we include timber production, fuelwood production, and access to nature (which could be used for gathering, and also links to culture). For cultural benefits we include coral reef tourism (as the only globally mapped form of marine-based tourism) and access to nature again (which in addition to gathering also captures recreation or other uses of nearby greenspace). Finally, for climate security we include total ecosystem carbon storage (as a global NCP). Below we briefly summarize the models used to map these local NCP (Extended Data Table 1) and global NCP (Extended Data Table 2), full information on each model is available in the SI Methods.

#### Local NCP

1. *Nitrogen retention* to regulate water quality for downstream populations is modeled using the InVEST^48^ Nutrient Delivery Ratio model, which is based on fertilizer application, precipitation, topography, and the retention capacity of vegetation, and has been previously used in global applications^49^. The people benefitting from nitrogen retention are those who would otherwise be exposed to nitrogen contamination in their drinking water. In this analysis, the number of people downstream were calculated for every pixel of habitat, to provide a sense of which habitat potentially benefits the most people. Ideally, to map realized nitrogen retention, we would be able to convert biophysical service production into a measure of change in well-being, whether monetary, in health terms, or otherwise. However, the state of the science and data available globally precludes this for most services, so our proxy was the number of people downstream who could potentially benefit from that retention. NCP for nitrogen retention is expressed as nitrogen retention on natural and semi-natural pixels multiplied by the number of people downstream of those pixels. (See SI Methods Section 1 for more detail.)
2. *Sediment retention* to regulate water quality for downstream populations is modeled by adapting the InVEST Sediment Delivery Ratio (SDR) model, which maps overland sediment generation and delivery to the stream using the Revised Universal Soil Loss Equation (RUSLE) and a conductivity index based on the upslope and downslope areas of each pixel. Ideally sediment retention would be delineated for reservoirs, irrigation canals, or other water delivery infrastructure that is most impacted by sedimentation, but lacking a comprehensive global dataset identifying all such infrastructure, we again use the proxy of number of people downstream (as described for nitrogen retention, above). NCP for sediment retention is expressed as sediment retention on natural and semi-natural pixels multiplied by the number of people downstream of those pixels. (See SI Methods Section 2.)
3. *Crop pollination* is modeled with a simplified version of InVEST, mapping the potential contribution of wild pollinators to nutrition production based on the sufficiency of habitat surrounding farmland and the pollination dependency of crops^49^. NCP for crop pollination is expressed in terms of the average equivalent number of people fed by pollination-dependent crops, attributed to nearby ecosystems based on the area of pollinator habitat within pollinator flight distance of crops. (See SI Methods Section 3.)
4. *Fodder production for livestock* is modeled using Version 3 of Co$ting Nature^50^. Supply of fodder is calculated as the livestock-accessible tonnes of dry matter productivity for the non-cropland cover fraction and demand is estimated by the head count of livestock in a grid cell multiplied by the average biomass intake requirements per animal. The NCP as fodder production for livestock is expressed in terms of an index (0-1), rescaled from the realized service which is reported as the smaller of the supply or demand (if consumption exceeds productivity, the gap is assumed to be met with feed). The best available global inputs for dry matter productivity, livestock headcount, cropland and land cover are used as input. (See SI Methods Section 4.)
5. *Timber production* includes commercial (e.g., for trade/export) and domestic (e.g., for local use) timber, modeled using Version 3 of Co$ting Nature as two spatially mutually exclusive layers, because they represent two different sets of beneficiaries. NCP for timber production is expressed as an index (0-1) based on forest productivity and accessibility for harvest. Total potential sustainable supply of timber is estimated from the best available global above-ground carbon stock map multiplied by fractional tree cover for rural areas only. The sustainable harvest is calculated as the reciprocal of the number of years taken to develop the stock at the annual sequestration rate, according to dry matter productivity data. Demand is calculated differently for commercial vs. domestic timber based on different assumptions of accessibility. Commercial accessibility is defined as within six hours’ travel time of a population center of >50K people and on slope gradients <70%. Domestic accessibility is defined as areas inaccessible for commercial harvest and harvest rates are based on per capita consumption multiplied by population within 10 km. (See SI Methods Section 5.)
6. *Fuelwood production* is calculated as a byproduct of the timber model from Version 3 of Co$ting Nature. NCP for fuelwood production, like timber, is represented as an index (0-1) based on forest productivity and accessibility for harvest, but in this case specifically by rural people. Fuelwood can overlap spatially with domestic and commercial timber use, given that domestic and commercial timber harvest will not consume all sustainably available woody biomass in all places, due to the slope gradient limit and/or in places where demand is less than supply, and fuelwood is often a by-product of timber harvest. (See SI Methods Section 6.)
7. *Flood regulation* is modeled using Version 2 of WaterWorld^31^. To map nature’s influence on flood risk reduction, we identify the upstream places where canopies, wetlands, and soils (green storage) retain and slowly release rainfall, to the benefit of downstream communities on floodplains. NCP for flood regulation is expressed as an index (0-1) based on “green” water storage multiplied by the number of people downstream on floodplains. (See SI Methods Section 7.)
8. *Access to nature* is used as a proxy for numerous direct and indirect benefits to people, such as recreation, hunting and gathering, aesthetics, mental and physical health, and other cultural values that depend on the ability of people to access nature. This proxy-NCP is expressed as the number of urban and rural^51^ people within 1 hour (or 6 hours, for sensitivity analysis) travel of natural and semi-natural habitat, taking the least-cost path (by foot, road, rail or boat) across a friction layer^14^ (See SI Methods Section 8.)
9. *Riverine fish catch* is based on spatial disaggregation of nationally reported catch for 2007-2014^16^ and updated to include catch estimated by household consumption surveys in 32 countries with severe underreporting^52^. Catches from large lakes were excluded. To spatially disaggregate the global catch of 13.3 ×10^6^ tonnes within country borders, a multiple linear regression model of total fish catch in river basins compiled from the literature was fitted with three predictor variables: population density, river discharge, and percent wetland cover (n=40, R^2^_adj_ =0.69). NCP for riverine fish harvest is represented as metric tonnes of fish caught per km^2^ of land area per year, spatially allocated to the locations of the harvest. (See SI Methods Section 9).
10. *Marine fish catch* is based on the Sea Around Us data to map fish catch for 2010-2014 within 30 min grid cells across the ocean^53,54^. NCP for marine fish harvest is represented as metric tonnes of fish caught per km^2^ per year, spatially disaggregated to the locations of the catch. (See SI Methods Section 10.)
11. *Coral reef tourism* is taken from the Mapping Ocean Wealth dataset^30^, which reports the NCP for coral reef-associated tourism as dollars of tourism expenditure (in deciles 1-10). National expenditure data are spatially distributed based on three independent sources: hotel rooms from the commercial Global Accommodation Reference Database (GARD), dive shops and dive sites from Diveboard, and user-generated photos from the image-sharing website, Flickr. (See SI Methods Section 11).
12. *Coastal risk reduction* is modeled with InVEST for terrestrial and coastal/off-shore habitats^55–58^, updating previous global modeling^49^ through the inclusion of new data and projecting the value back to the habitat. Coastal risk reduction depends on the physical exposure to coastal hazards (based on wind, waves, sea level rise, geomorphology, bathymetry). with and without natural habitat to attenuate storm surge, and the people exposed. NCP for coastal risk reduction is expressed as a unitless index of the coastal risk reduced by habitat multiplied by the number of people within the protective distance of the habitat. (See SI Methods Section 12.)

#### Global NCP

1. *Vulnerable terrestrial ecosystem carbon storage* is mapped as the above-ground and below-ground ecosystem carbon lost in a ‘typical’ disturbance event, rather than the total stock^20^. This includes terrestrial and coastal (mangrove, salt marsh, seagrass) ecosystem carbon pools (aboveground, belowground, and soils), based on what carbon is likely to be released if the ecosystem were converted. (See SI Methods Section 13.)
2. *Atmospheric moisture recycling* is the process of water arising from the surface of the earth as evaporation, flowing through the atmosphere as water vapor, and returning to the surface of the earth as precipitation. Sources of evaporation include canopy interception, soil interception, soil evaporation, vegetation transpiration, and open water evaporation. We employed an Eulerian moisture tracking model, WAM-2 layers^21^, to quantify the source of moisture, where it travels through the atmosphere, and where it falls out downwind. The NCP of moisture recycling, which is to say the moisture associated with intact vegetation, is expressed as the volume of water evapo-transpired that falls on all rainfed productive land (cropland, rangelands, and working forests^59^). (See SI Methods Section 14).

### Attribution of value to natural assets

The first step in identifying critical natural assets is to attribute the magnitude of benefits and, where possible, the number of beneficiaries, to the ecosystems providing those benefits (e.g., attributing the value of pollination occurring on croplands to the nearby habitat supplying the pollinators, or the coastal risk reduced and number of people protected along the coastline to offshore as well as onshore habitats). We define natural assets as natural and semi-natural terrestrial ecosystems (including semi-natural lands like rangelands and production forests, but excluding cropland, urban areas, bare areas, and permanent snow and ice; Extended Data Table 3) and inland and marine waters. Model outputs for pollination and coastal risk reduction are mapped back to habitat based on the pollinator flight distance (SI Methods Section 3) and protective distance of coastal habitat (SI Methods Section 12), respectively. For sediment and nitrogen retention, the count of people downstream of each habitat pixel was summed according to a hydrologic flow accumulation (SI Methods Section 1) and for nature access, the count of people was calculated for each pixels by summing the population pixels within 1 hour travel time according to a friction surface (SI Methods Section 8). All other model outputs are coarser than ESA resolution and are masked to the LULC types defined as natural assets relevant to that NCP (e.g., only forests for timber but excluding forests for grazing; Extended Data Table 3).

### Optimization of NCP

Using integer linear programming (prioritizr, SI Methods Section 22), we identify minimum areas required 1) within each country’s land borders and marine Exclusive Economic Zones (EEZs) for the local NCP and 2) within all global land area (excluding Antarctica) and all countries’ combined EEZ area for the global NCP, to reach target levels (ranging from 5% to 100%) of every NCP. This optimization selects for the highest values across all NCP, providing the most benefit and/or to the greatest number of people, but not accounting for complementarity or redundancies of adjacent pixels (i.e., not dynamically optimizing after each pixel’s selection). We define the land and marine area for the 90% target as “critical natural assets” because the remaining 10% of aggregate NCP value requires disproportionately more area to achieve. Land and marine borders are based on Flanders Marine Institute (2020; Extended Data Table 2), and overlapping claims are excluded from the national analyses. The 12 local NCP are optimized for each country, then aggregated globally, while the two global NCP are optimized globally. In addition to these two main optimizations, we assess the sensitivity to scale by optimizing the 12 local NCP globally (instead of by each country), both with and without the 2 global NCP, and by substituting different scales of beneficiaries mapping for people downstream and for access to nature (Extended Data Table 5). We also assess the sensitivity of the area and location of critical natural assets (the optimization solution for the 90% target) to different NCP combinations. These variations include optimizing for each NCP individually, and optimizing for all NCP but dropping each local NCP from the set of 12 to evaluate its effect on the overall optimization (Extended Data Table 5). We also examine the correspondence between NCP and the robustness of these different solutions, by calculating the percentage of area shared by different pairs of services (Extended Data Table 4) or the percentage of area shared by all solutions (SI Table 1). We summarize the land and ocean areas required by country in SI Table 1.

### Number of people benefiting from critical natural assets

We map the areas benefiting from critical natural assets in order to calculate the number of direct beneficiaries of these assets, and to compare the number of beneficiaries to the number of people living on the lands comprising these critical natural assets. For this analysis we are only able to include NCP for which the flow of the benefit can be spatially delineated: downstream water quality regulation (sediment retention, nitrogen retention), flood mitigation, nature access, fuelwood provision, and coastal risk reduction. The benefiting areas of some of the material NCP that are traded (fish, timber, livestock, crops that are pollinated) or the location of people who buy those traded goods are not easily mapped, so people benefiting from these NCP are not included in this analysis of beneficiaries. However, people within an hour of critical natural assets may provide a surrogate for many of the material NCP that are locally consumed. For water quality regulation, we take the population within the areas downstream (SI Methods Section 1) of critical natural assets. For nature access, we take the population within an hour’s travel (by foot, car, boat or rail; SI Methods Section 8) of critical natural assets. Likewise, for the other NCP we take the relevant population downstream, within the protective distance, or a gathering distance of critical natural assets. The relevant population for each NCP is considered to be the total global population for nature access and water quality regulation, but is limited to the total population living within 10 km of floodplains for flood mitigation, population along coastlines (in exposed areas: <10m above mean sea level) for coastal risk reduction, and rural poor populations for fuelwood. Total “local” beneficiaries are calculated through the intersection of areas benefiting from different NCP, to avoid double-counting people in areas of overlap. We calculate the number of people and percent of relevant population benefiting globally for each NCP (Fig. 2b) and the total “local” beneficiaries globally (Fig. 2a) and by country (SI Table 2).

### Overlap analysis

We evaluate how well local and global critical natural assets align spatially with each other, and with biodiversity (terrestrial vertebrate species Area of Habitat (AOH)^22^; SI Methods Section 15) and cultural diversity (proxied by the number of Indigenous and non-migrant languages^23^; SI Methods Section 16), to identify synergies between these different potential priorities. To examine the level of overlap between areas identified as critical for the 12 local NCP vs. the 2 global NCP, we calculate the area (globally and by country; SI Table 1) where local NCP are selected by the optimization (at the 90% target level) and global NCP are not, where global NCP are selected by the optimization and local NCP are not, and where both are selected by their respective optimizations (the overlap). To calculate the species and languages represented by critical natural assets, we count the number of species whose AOH area targets overlaps these areas (SI Table 3) and the number of languages partially intersecting these areas (SI Table 4) globally and within each country. (See SI Methods Section 23 for more detail.)

## Acknowledgments

We thank all the participants of two working groups hosted by Conservation International and the Natural Capital Project for their insights and intellectual contributions. For further advice or assistance, we thank Alison Adams, Katrina Brandon, Kate Brauman, Abigail Cramer, Gretchen Daily, Jon Fisher, Rachelle Gould, Lisa Mandle, Jamie Montgomery, Amanda Rodewald, David Rossiter, Elizabeth Selig, Adrian Vogl, and T. Max Wright. The two working groups that provided the foundation for this analysis were funded by the Marcus and Marianne Wallenberg Foundation and a grant provided by Betty and Gordon Moore.

## Author Contributions

RCK, RPS, MM, EFC, LG, PWK, KLW, PBM, MN, CRL, PRR, MS, JAJ, AvS, and RAW provided data, RS and MSM performed the optimizations, RCK, RPS, LG, and PRR performed additional analyses, CB, SD, UP, GST, MRS, and WRT provided framing for policy relevance, RCK, RAN, PMC, SP, and DH directed the project, RCK and RAN wrote the first draft of the paper, and all authors contributed to revisions.

## Data/Code Availability Statement

All final outputs and code are available on Open Science Framework: https://osf.io/r5xz7/.

## Additional Information

Supplementary Information is available for this paper. Correspondence and requests for materials should be addressed to.

## References

1. IPBES. Summary for policymakers of the global assessment report on biodiversity and ecosystem services of the Intergovernmental Science-Policy Platform on Biodiversity and Ecosystem Services. (IPBES secretariat, Bonn, Germany, 2019).

2. Dobson, A. P. et al. Ecology and economics for pandemic prevention. Science 369, 379–381 (2020).

3. Diaz, S. et al. Assessing nature’s contributions to people. Science 359, 270–272 (2018).

4. Hole, DG., Collins, P., Tesfaw, A., Barrera, L., Mascia, MB and Turner, WR. Make nature’s role visible to achieve the SDGs. Global Sustainability (2021).

5. Roe, S. et al. Contribution of the land sector to a 1.5°C world. Nat. Clim. Chang. 9, (2019).

6. Wilson, E. O. Half-earth: Our Planet’s Fight for Life. 256 (WW Norton & Company, 2016).

7. Baillie, J. & Zhang, Y.-P. Space for nature. Science 361, 1051 (2018).

8. Büscher, B. et al. Half-Earth or Whole Earth? Radical ideas for conservation, and their implications. Oryx 51, 407–410 (2017).

9. Schmidt-Traub, G. et al. Integrating climate, biodiversity, and sustainable land-use strategies: innovations from China. National Science Review (2020) doi:10.1093/nsr/nwaa139.

10. Chaplin-Kramer, R. et al. Mapping the planet’s critical natural assets for people. bioRxiv 2020.11.08.361014 (2021) doi:10.1101/2020.11.08.361014.

11. Myers, S. S. et al. Human health impacts of ecosystem alteration. Proc. Natl. Acad. Sci. U. S. A. 110, 201218656 (2013).

12. Strassburg, B. B. N. et al. Global priority areas for ecosystem restoration. Nature 586, 724–729 (2020).

13. Kennedy, C. M., Oakleaf, J. R., Theobald, D. M., Baruch-Mordo, S. & Kiesecker, J. Managing the middle: A shift in conservation priorities based on the global human modification gradient. Glob. Chang. Biol. 25, 811–826 (2019).

14. Weiss, D. J. et al. A global map of travel time to cities to assess inequalities in accessibility in 2015. Nature 553, 333–336 (2018).

15. Lebedys, A. & Yanshu, L. Contribution of the forestry sector to national economies, 1990-2011. Forest Finance Working Paper (FAO) eng no. 09 (2014).

16. McIntyre, P. B., Liermann, C. A. R. & Revenga, C. Linking freshwater fishery management to global food security and biodiversity conservation. Proceedings of the National Academy of Sciences 113, 12880–12885 (2016).

17. Selig, E. R. et al. Mapping global human dependence on marine ecosystems. Conservation Letters e12617 (2018).

18. FAO. World Livestock: Transforming the livestock sector through the sustainable development goals. http://bearecon.com/portfolio-data/fao-ldg/ca1201en.pdf (2018).

19. Fedele, G., Donatti, C. I., Bornacelly, I. & Hole, D. G. Nature-dependent people: Mapping human direct use of nature for basic needs across the tropics. Glob. Environ. Change 102368 (2021).

20. Noon, M. L. et al. Mapping the irrecoverable carbon in Earth’s ecosystems. Nature Sustainability 5, 37–46 (2021).

21. Keys, P. W., Wang-Erlandsson, L. & Gordon, L. J. Revealing Invisible Water: Moisture Recycling as an Ecosystem Service. PLoS One 11, e0151993 (2016).

22. Brooks, T. M. et al. Measuring terrestrial area of habitat (AOH) and its utility for the IUCN Red List. Trends Ecol. Evol. 34, 977–986 (2019).

23. Gorenflo, L. J., Romaine, S., Mittermeier, R. A. & Walker-Painemilla, K. Co-occurrence of linguistic and biological diversity in biodiversity hotspots and high biodiversity wilderness areas. Proceedings of the National Academy of Sciences 109, 8032–8037 (2012).

24. Convention on Biodiversity. First draft of the post-2020 global biodiversity framework. CBD/WG2020/3/5. (2021).

25. O’Connor, L. M. J. et al. Balancing conservation priorities for nature and for people in Europe. Science 372, 856–860 (2021).

26. Jung, M. et al. Areas of global importance for conserving terrestrial biodiversity, carbon and water. Nat. Ecol. Evol. (2021) doi:10.1038/s41559-021-01528-7.

27. Sala, E. et al. Protecting the global ocean for biodiversity, food and climate. Nature 592, 397–402 (2021).

28. Dinerstein, E. et al. A “Global Safety Net” to reverse biodiversity loss and stabilize Earth’s climate. Sci Adv 6, (2020).

29. Chan, K. M. A., Gould, R. K. & Pascual, U. Editorial overview: Relational values: what are they, and what’s the fuss about? Curr. Opin. Environ. Sustain. 35, A1–A7 (2018).

30. Spalding, M. et al. Mapping the global value and distribution of coral reef tourism. Mar. Policy 82, 104–113 (2017).

31. Gunnell, K., Mulligan, M., Francis, R. A. & Hole, D. G. Evaluating natural infrastructure for flood management within the watersheds of selected global cities. Sci. Total Environ. 670, 411–424 (2019).

32. Bagstad, K. J., Semmens, D. J., Waage, S. & Winthrop, R. A comparative assessment of decision-support tools for ecosystem services quantification and valuation. Ecosystem Services 5, 27–39 (2013).

33. van der Ent, R. J., Tuinenburg, O. A., Knoche, H.-R., Kunstmann, H. & Savenije, H. H. G. Should we use a simple or complex model for moisture recycling and atmospheric moisture tracking? Hydrol. Earth Syst. Sci. 17, 4869–4884 (2013).

34. Redhead, J. W. et al. National scale evaluation of the InVEST nutrient retention model in the United Kingdom. Sci. Total Environ. 610-611, 666–677 (2018).

35. Arkema, K. K. et al. Coastal habitats shield people and property from sea-level rise and storms. Nat. Clim. Chang. 3, 913–918 (2013).

36. Cabral, P. et al. Assessing Mozambique’s exposure to coastal climate hazards and erosion. International Journal of Disaster Risk Reduction 23, 45–52 (2017).

37. Benez-Secanho, F. J. & Dwivedi, P. Does Quantification of Ecosystem Services Depend Upon Scale (Resolution and Extent)? A Case Study Using the InVEST Nutrient Delivery Ratio Model in Georgia, United States. Environments 6, 52 (2019).

38. Hooftman, D. A. P. et al. Reducing uncertainty in ecosystem service modelling through weighted ensembles. Ecosystem Services 53, 101398 (2022).

39. Willcock, S. et al. Ensembles of ecosystem service models can improve accuracy and indicate uncertainty. Sci. Total Environ. 747, 141006 (2020).

40. Johnson, J. A. et al. Global Futures: Modelling the Global Economic Impacts of Environmental Change to Support Policy Making. (2020).

41. Kapsar, K. E. et al. Telecoupling research: The first five years. Sustain. Sci. Pract. Policy 11, 1033 (2019).

42. Hannah, L. et al. 30% land conservation and climate action reduces tropical extinction risk by more than 50%. Ecography 43, 943–953 (2020).

43. Chaplin-Kramer, R. et al. Conservation needs to integrate knowledge across scales. Nature ecology & evolution vol. 6 118–119 (2022).

44. Neugarten, R. A. et al. Trends in protected area representation of biodiversity and ecosystem services in five tropical countries. Ecosystem Services 42, 101078 (2020).

45. Evans, K., Guariguata, M. R. & Brancalion, P. H. S. Participatory monitoring to connect local and global priorities for forest restoration. Conserv. Biol. 32, 525–534 (2018).

46. Barnes, M. D., Glew, L., Wyborn, C. & Craigie, I. D. Prevent perverse outcomes from global protected area policy. Nat Ecol Evol 2, 759–762 (2018).

47. Rose, A. N., McKee, J. J., Urban, M. L. & Bright, E. A. LandScan 2017. (2018).

48. Sharp, R. et al. InVEST 3.8.0 User’s Guide. http://releases.naturalcapitalproject.org/invest-userguide/latest/ (2020).

49. Chaplin-Kramer, R. et al. Global modeling of nature’s contributions to people. Science 366, 255–258 (2019).

50. Mulligan, M. et al. Mapping nature’s contribution to SDG 6 and implications for other SDGs at policy relevant scales. Remote Sens. Environ. 239, 111671 (2020).

51. Cattaneo, A., Nelson, A. & McMenomy, T. Global mapping of urban-rural catchment areas reveals unequal access to services. Proc. Natl. Acad. Sci. U. S. A. 118, (2021).

52. Fluet-Chouinard, E., Funge-Smith, S. & McIntyre, P. B. Global hidden harvest of freshwater fish revealed by household surveys. Proc. Natl. Acad. Sci. U. S. A. 115, 7623–7628 (2018).

53. Watson, R. A. & Tidd, A. Mapping nearly a century and a half of global marine fishing: 1869–2015. Mar. Policy 93, 171–177 (2018).

54. Pauly, D., Zeller, D. & Palomares, M. D. Sea Around Us Concepts, Design and Data (2020). Available at: http://seaaroundus.org.

55. Burke, L., Reytar, K., Spalding, M. & Perry, A. Reefs at Risk Revisited. http://www.wri.org/publication/reefs-risk-revisited (2011).

56. UNEP-WCMC & Short, F. T. Global distribution of seagrasses (version 6.0). http://data.unepwcmc.org/datasets/7 (2017).

57. Mcowen, C. et al. A global map of saltmarshes. Biodiversity Data Journal 5, e11764 (2017).

58. Bunting, P. et al. The Global Mangrove Watch—A New 2010 Global Baseline of Mangrove Extent. Remote Sensing 10, 1669 (2018).

59. Ellis, E. C. & Ramankutty, N. Putting people in the map: anthropogenic biomes of the world. Front. Ecol. Environ. 6, 439–447 (2008).

