## Extended data for "Mapping the planet’s critical natural assets"

**Extended Data Table 1 | List of local nature's contributions to people (NCP) included in this analysis**

| NCP | Source | Units | Original resolution | Realm |
|---|---|---|---|---|
| Nitrogen retention for water quality regulation | Chaplin-Kramer et al. 2019 (Ref. 10), InVEST (updated) | Kg nitrogen retained * number of people downstream | 10 arc-sec (~300 m) | Land |
| Sediment retention for water quality regulation | Chaplin Kramer et al., InVEST (new for this analysis) | T sediment retained * number of people downstream | 10 arc-sec (~300 m) | Land |
| Crop pollination contribution to nutrition production | Chaplin-Kramer et al. 2019 (Ref. 10), InVEST (updated) | "People fed equivalents"; average of pollination-derived energy (KJ), folate, and vitamin A production divided by annual dietary requirements per capita. | 10 arc-sec (~300 m) | Land |
| Fodder production for livestock | Mulligan et al. 2020 (Ref. 51), Co$ting Nature v3 (updated) | Index (0-1) of dry matter productivity utilized by livestock | 5 arc-min (~10 km) | Land |
| Timber production (commercial and domestic) | Mulligan et al. 2020 (Ref. 51), Co$ting Nature v3 (updated) | Index (0-1) of accessible timber harvest for commercial & domestic use (optimized separately) | 5 arc-min (~10 km) | Land |
| Fuel wood production | Mulligan et al. 2020 (Ref. 51), Co$ting Nature v3 (updated) | Index (0-1) of fuel wood accessible to local rural communities | 5 arc-min (~10 km) | Land |
| Flood regulation | Gunnell et al. 2019 (Ref. 32), WaterWorld v2 (updated) | Index (0-1) of green water storage * number of people downstream. | 5 arc-min (~10 km) | Land |
| Access to nature (local recreation and gathering) | Chaplin Kramer et al. (new for this analysis) | Count of people within 10 km of natural and semi-natural habitat | 10 arc-sec (~300 m) | Land |
| Riverine fish catch | McIntyre et al. 2016 (Ref. 16) (updated) | Metric tonnes of fish caught per sq km (of land area) per year | 5 arc-min (~10 km) | Land, freshwater |
| Marine fish catch | Watson and Tidd 2018 (Ref. 54); (updated) | Metric tonnes of fish caught per sq km (of ocean) per year | 30 arc-min (~55 km) | Ocean |
| Coral reef tourism (nature-based recreation and associated livelihoods) | Spalding et al. 2017 (Ref. 31) | Dollar expenditures (expressed in deciles 1-10) | 30 arc-sec (~1 km) | Ocean |
| Coastal risk reduction | Chaplin-Kramer et al. 2019 (Ref. 10), InVEST (updated) | Unitless risk reduction index * number of people within protective distance | 10 arc-sec (~300 m) | Land and ocean |

All NCP (mapped in Extended Data Fig. 1) are "realized," either as an end use or benefit (e.g., timber or fish harvest per unit area of land or water), or, where possible given current data, weighted by number of beneficiaries. All NCP are attributed to the "natural" ecosystems providing the benefit (see Table 4), resampled to 2 km for prioritization. "Source" indicates the data source, but where indicated, datasets were updated (or newly generated) for this analysis by the authors, as detailed in the SI Methods.

**Extended Data Table 2 | Global climate NCP and other supporting datasets**

| Data set | Source | Units | Original resolution |
|---|---|---|---|
| ***Climate regulation NCP (not included in national-level optimization; optimized globally)*** | | | |
| Ecosystem carbon storage* | Noon et al2021 (Ref. 30) | Tonnes of carbon/ha (for terrestrial ecosystems and mangroves) | 1 arc-sec (~30 m) |
| Atmospheric moisture recycling* | Keys et al. 2016 (Ref. 21) (updated) | Fraction of evapotranspiration from vegetation that is providing precipitation to rainfed productive lands | 1.5 degree |
| ***Biological diversity*** | | | |
| Terrestrial vertebrates Area of Habitat (AOH)* | Roerhdanz, Conservation International, (new for this analysis) Based on IUCN Red List data and methods from Brooks et al. 2019 (Ref. 22) | N/A (species Area of Habitat (AOH) polygons for 29,000 mammals, birds, amphibians, and reptiles) | N/A (vector) |
| ***Cultural diversity*** | | | |
| Languages* | Gorenflo et al. 2012 (Ref. 23) (updated) | N/A (indigenous and non-migrant language range polygons) | N/A (vector) |
| ***Additional input datasets*** | | | |
| Land cover | ESA Climate Change Initiative 2017 | N/A (Masks for NCP layers included all land cover classes from ESA 2015 except for cropland, mosaic cropland, urban areas, bare areas, water bodies, permanent snow & ice) | 10 arc-sec (~300 m) |
| Coastal habitat | ESA Climate Change Initiative 2017 (terrestrial coastal habitat), Burke et al. 2011 (Ref. 56) (coral reefs), Bunting et al. 2018 (Ref. 59) (mangroves), UNEP-WCMC & Short 2018 (Ref. 57) (seagrass), Mcowen et al. 2017 (Ref. 58) (salt marsh) | N/A (coastal habitat types) | ESA: 10 arc-sec<br>Burke et al. 2011: N/A<br>Bunting et al:<br>UNEP-WCMC and Short 2018:<br>Mcowen et al: N/A (vectors) |
| Dry matter productivity* | Copernicus Service Information 2019 | kg/ha/day (converted to a 10-year average 2009-2018, so kg/ha) | 30 arc-sec (~1 km) |
| Human population* | Landscan 2017, Rose et al. 2018 (Ref. 48) | Ambient or average day / night population count (people per square km) | 30 arc-sec (~1 km) |
| Urban and rural catchment areas* | Cattaneo et al. 2021 (Ref. 52) | Urban to rural gradient classification (1 being most urban 30 being most rural) | 30 arc-sec (~1 km) |
| Friction surface* | Weiss et al. 2018 (Ref. 14) | Minutes required to move one meter | 30 arc-sec (~1 km) |
| Land and ocean boundaries | Flanders Marine Institute 2020<br>Union of ESRI country shapefile and Exclusive Economic Zones v3 | N/A (polygons) | NA |

*These datasets are mapped only for land.

**Extended Data Table 3 | ESA Land Use/Land Cover (LULC) classes to which different terrestrial NCP were masked**

| ID | Description | Grazing: herbaceous only | Timber: forests only | Fuelwood: woody only | All other terrestrial NCP: all natural/semi-natural | Inland fisheries: + water |
|---|---|---|---|---|---|---|
| 10 | Cropland, rainfed | | | | | |
| 11 | Cropland, rainfed, herbaceous cover | | | | | |
| 12 | Cropland, rainfed, tree or shrub cover | | | | | |
| 20 | Cropland, irrigated or post-flooding | | | | | |
| 30 | Mosaic cropland (>50%) / natural vegetation (tree, shrub, herbaceous cover)(<50%) | X | X | X | X | X |
| 40 | Mosaic natural vegetation (tree, shrub, herbaceous cover) (>50%) / cropland(<50%) | X | X | X | X | X |
| 50 | Tree cover, broadleaved, evergreen, closed to open (>15%) | | X | X | X | X |
| 60-62 | Tree cover, broadleaved, deciduous, closed to open (>15%) | | X | X | X | X |
| 70-72 | Tree cover, needleleaved, evergreen, closed to open (>15%) | | X | X | X | X |
| 80-82 | Tree cover, needleleaved, deciduous, closed to open (>15%) | | X | X | X | X |
| 90 | Tree cover, mixed leaf type (broadleaved and needleleaved) | | X | X | X | X |
| 100 | Mosaic tree and shrub (>50%) / herbaceous cover (<50%) | X | X | X | X | X |
| 110 | Mosaic herbaceous cover (>50%) / tree and shrub (<50%) | X | X | X | X | X |
| 120-122 | Shrubland | X | | X | X | X |
| 130 | Grassland | X | | | X | X |
| 140 | Lichens and mosses | X | | | X | X |
| 150 | Sparse vegetation (tree, shrub, herbaceous cover) (<15%) | X | X | X | X | X |
| 151 | Sparse tree (<15%) | X | X | X | X | X |
| 152 | Sparse shrub (<15%) | X | | X | X | X |
| 153 | Sparse herbaceous cover (<15%) | X | | | X | X |
| 160 | Tree cover, flooded, fresh or brackish water | | X | X | X | X |
| 170 | Tree cover, flooded, saline water | | X | X | X | X |
| 180 | Shrub/ herbaceous cover, flooded, fresh/saline/brackish water | X | | X | X | X |
| 190 | Urban areas | | | | | |
| 200-202 | Bare areas | | | | | |
| 210 | Water bodies | | | | | X |
| 220 | Permanent snow and ice | | | | | |

**Extended Table 4 | Correspondence between different pairs of nature's contributions to people (NCP), based on individual optimizations for each NCP.** Vulnerable carbon and moisture regulation were optimized globally; all other NCP were optimized at the country level. Columns show the percent of the area required to maintain 90% of that NCP alone, and then the % of overlap with the area required by other NCP. As different NCP require vastly different areas, overlaps between NCP pairs may matter more for one of the NCP than the other (e.g., sediment retention requires less than a third of the area of flood mitigation or nitrogen retention, so while the overlap accounts for a large percentage of the area required by sediment, it accounts for a much smaller percentage of the areas required by the other two). The highest correspondence between NCP is shown in green (darker shades highlighting the top 10% of overlap values, lighter shades highlighting the top 25%).

| | | carbon | moisture | coastal | flood | fuelwood | riverine fish | grazing | nature access | nitrogen | pollination | sediment | timber |
|---|---|---|---|---|---|---|---|---|---|---|---|---|---|
| *Total area required (million sq km)* | | *46.6* | *46.0* | *0.2* | *28.1* | *14.6* | *3.2* | *19.9* | *15.1* | *28.6* | *4.6* | *9.3* | *19.7* |
| *% overlap with area required by:* | carbon | | 71% | 36% | 76% | 76% | 72% | 51% | 63% | 72% | 60% | 66% | 83% |
| | moisture | 70% | | 5% | 79% | 74% | 62% | 68% | 70% | 79% | 66% | 78% | 73% |
| | coastal | 0.14% | 0.02% | | 0.02% | 0.23% | 0.44% | 0.05% | 0.41% | 0.02% | 0.32% | 0.00% | 0.15% |
| | flood | 46% | 48% | 2% | | 64% | 52% | 44% | 57% | 70% | 59% | 66% | 61% |
| | fuelwood | 24% | 23% | 19% | 33% | | 35% | - | 55% | 33% | 53% | 38% | 59% |
| | riverine fish | 5% | 4% | 8% | 6% | 8% | | 4% | 9% | 5% | 12% | 6% | 6% |
| | grazing | 22% | 29% | 5% | 31% | - | 27% | | 47% | 36% | 55% | 44% | - |
| | nature access | 21% | 23% | 34% | 31% | 57% | 43% | 36% | | 32% | 63% | 38% | 37% |
| | nitrogen | 44% | 49% | 3% | 71% | 64% | 46% | 51% | 61% | | 65% | 83% | 59% |
| | pollination | 6% | 7% | 8% | 10% | 17% | 18% | 13% | 19% | 10% | | 15% | 10% |
| | sediment | 13% | 16% | 0% | 22% | 24% | 16% | 21% | 24% | 27% | 31% | | 20% |
| | timber | 35% | 31% | 16% | 43% | 80% | 39% | - | 49% | 41% | 43% | 43% | |

| | | coastal | marine fish | reef tourism |
|---|---|---|---|---|
| | *Total area required (million sq km)* | *0.10* | *33.797* | *0.15* |
| *% overlap with area required by:* | coastal | | 0.3% | 21% |
| | marine fish | 92% | | 82% |
| | reef tourism | 32% | 0.4% | |

**Extended Data Table 5 | Total area, land area and EEZ area required to provide 90% of current levels of NCP**

| Optimization description | % Total area | % Land | % EEZ |
|---|---|---|---|
| 1. Local Critical Natural Assets (LCNA):<br>Local NCP (12 NCP without global climate NCP) together, optimized by country | 27.3% | 30.4% | 24.2% |
| 2. Global Critical Natural Assets (GCNA):<br>Global climate NCP (carbon and moisture), optimized globally | 18.9% | 38.5% | 0.0% |
| 3. Total area of local NCP optimized by country + global NCP optimized globally (LCNA + GCNA) | 34.1% | 44.3% | 24.2% |
| ***Sensitivity to scale*** | | | |
| 4. Local NCP, optimized globally | 17.6% | 22.5% | 12.9% |
| 5. All NCP (the 12 + carbon + moisture), optimized globally | 26.0% | 39.5% | 12.9% |
| 6. Local NCP (as in Row 1) but with alternate nature access (longer travel time) | 27.6% | 31.0% | 24.2% |
| 7. Local NCP (as in Row 1) but with alternate hydro NCP (shorter attenuation) | 25.1% | 26.1% | 24.2% |
| ***Sensitivity to NCP: 11 local NCP optimized by country, as in Row 1, but dropping 1 NCP*** | | | |
| 8. Drop coastal risk reduction | 27.2% | 30.4% | 24.2% |
| 9. Drop timber production | 26.8% | 29.4% | 24.2% |
| 10. Drop flood mitigation | 26.7% | 29.2% | 24.2% |
| 11. Drop fuelwood | 27.3% | 30.4% | 24.2% |
| 12. Drop riverine fish | 27.2% | 30.3% | 24.2% |
| 13. Drop grazing | 25.7% | 27.1% | 24.2% |
| 14. Drop marine fish production | 15.0% | 30.3% | 0.2% |
| 15. Drop nature access (within 1 hour travel) | 27.2% | 30.2% | 24.2% |
| 16. Drop nitrogen retention | 26.8% | 29.5% | 24.2% |
| 17. Drop pollination | 27.2% | 30.4% | 24.2% |
| 18. Drop reef tourism | 27.3% | 30.4% | 24.2% |
| 19. Drop sediment retention | 27.2% | 30.4% | 24.2% |
| ***Individual NCP optimized by country (Rows 20-32) or globally (Rows 33-34):*** | | | |
| 20. Only coastal risk reduction | 0.1% | 0.1% | 0.1% |
| 21. Only timber production | 7.2% | 14.7% | 0.0% |
| 22. Only flood mitigation | 10.3% | 20.9% | 0.0% |
| 23. Only fuelwood | 5.3% | 10.8% | 0.0% |
| 24. Only riverine fish | 1.2% | 2.3% | 0.0% |
| 25. Only grazing | 7.2% | 14.7% | 0.0% |
| 26. Only marine fish production | 12.4% | 0.0% | 24.2% |
| 27. Only nature access within 1 hour travel (vs. within 6 hours) | 5.5% (9.3%) | 11.2% (18.8%) | 0.0% |
| 28. Only nitrogen retention with 500 km flow attenuation (vs. with 50 km) | 10.4% (7.3%) | 21.2% (14.9%) | 0.0% |
| 30. Only pollination | 1.7% | 3.4% | 0.0% |
| 31. Only reef tourism | 0.1% | 0.0% | 0.1% |
| 32. Only sediment retention with 500 km flow attenuation (vs. with 50 km) | 3.4% (2.4%) | 6.9% (4.9%) | 0.0% |
| 33. Only carbon | 17.0% | 34.5% | 0.1% |
| 34. Only moisture regulation | 16.8% | 34.1% | 0.0% |

**Extended Data Figure 1 | Individual maps for the 14 of Nature's Contributions to People (NCP) included in critical natural assets**

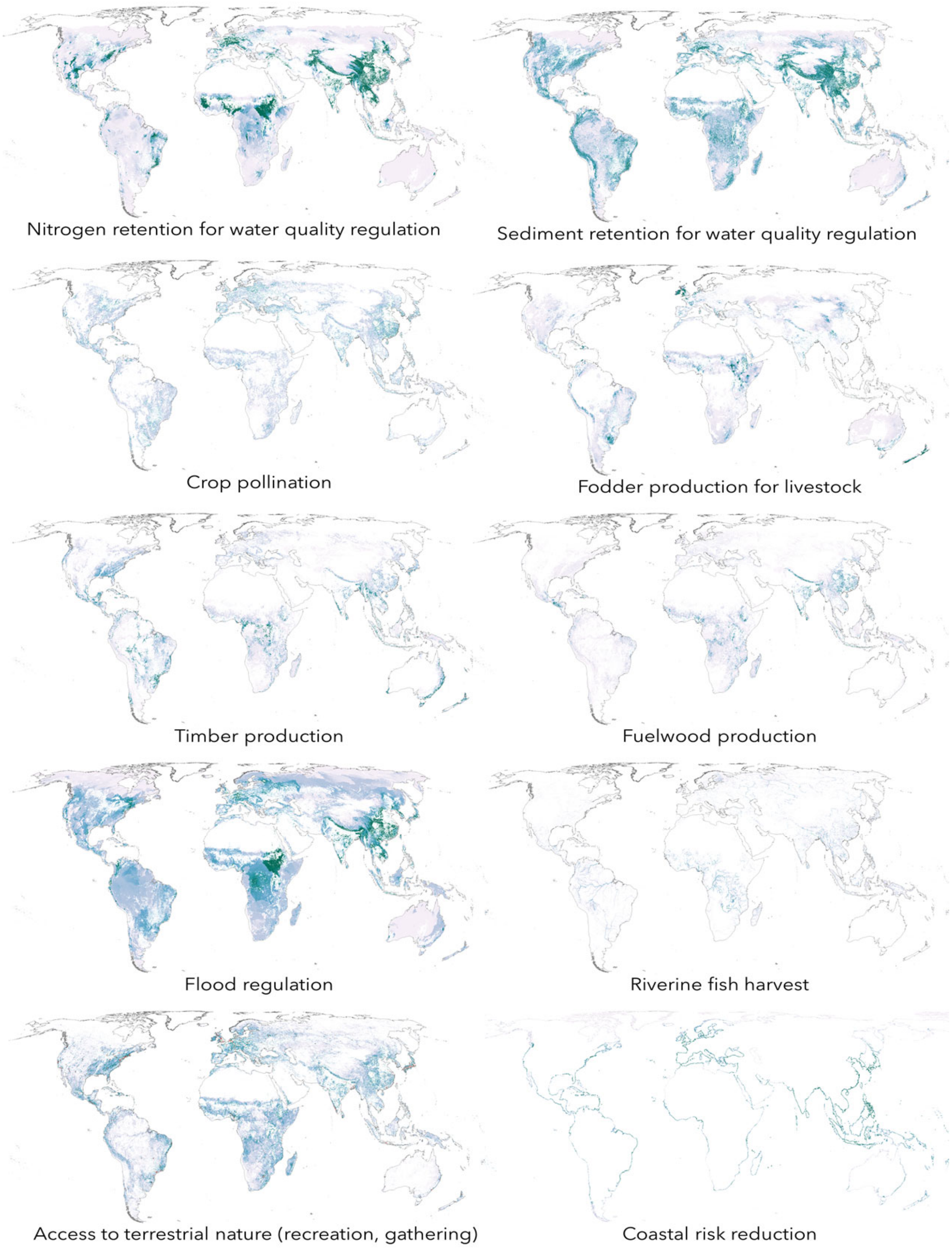

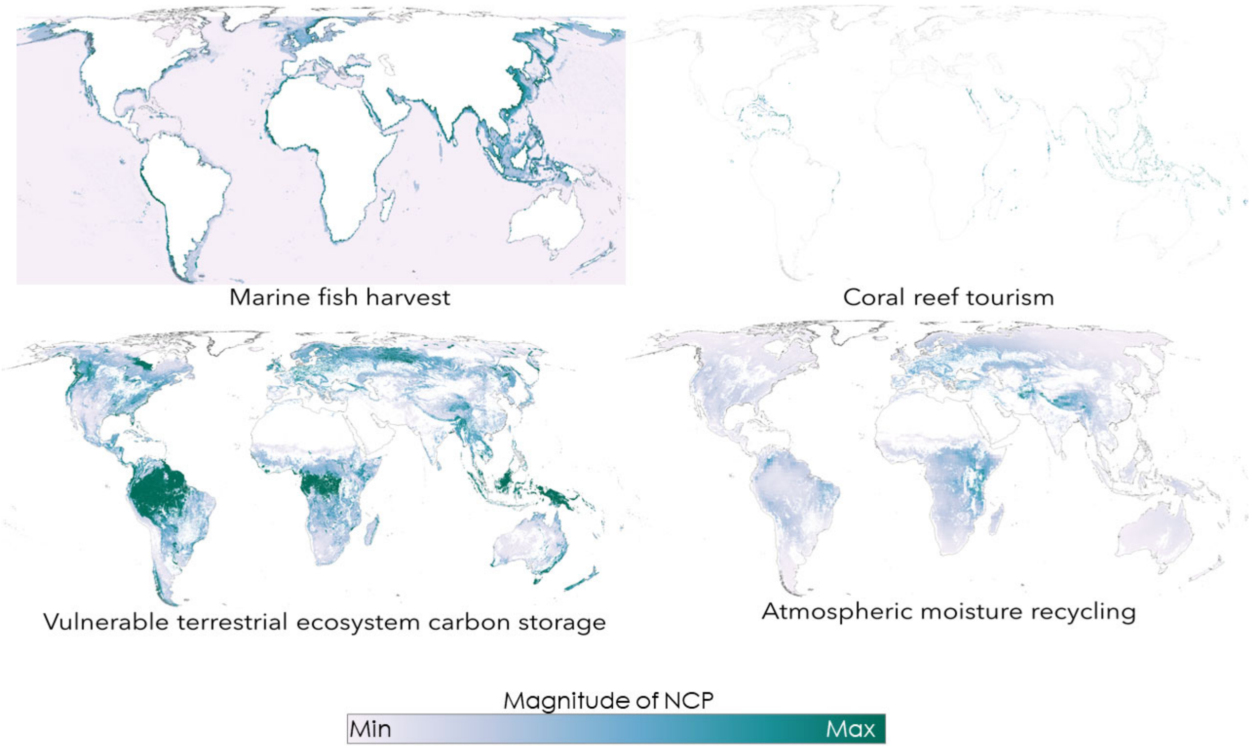
Marine fish harvest
Coral reef tourism
Vulnerable terrestrial ecosystem carbon storage
Atmospheric moisture recycling
Magnitude of NCP
Min
Max

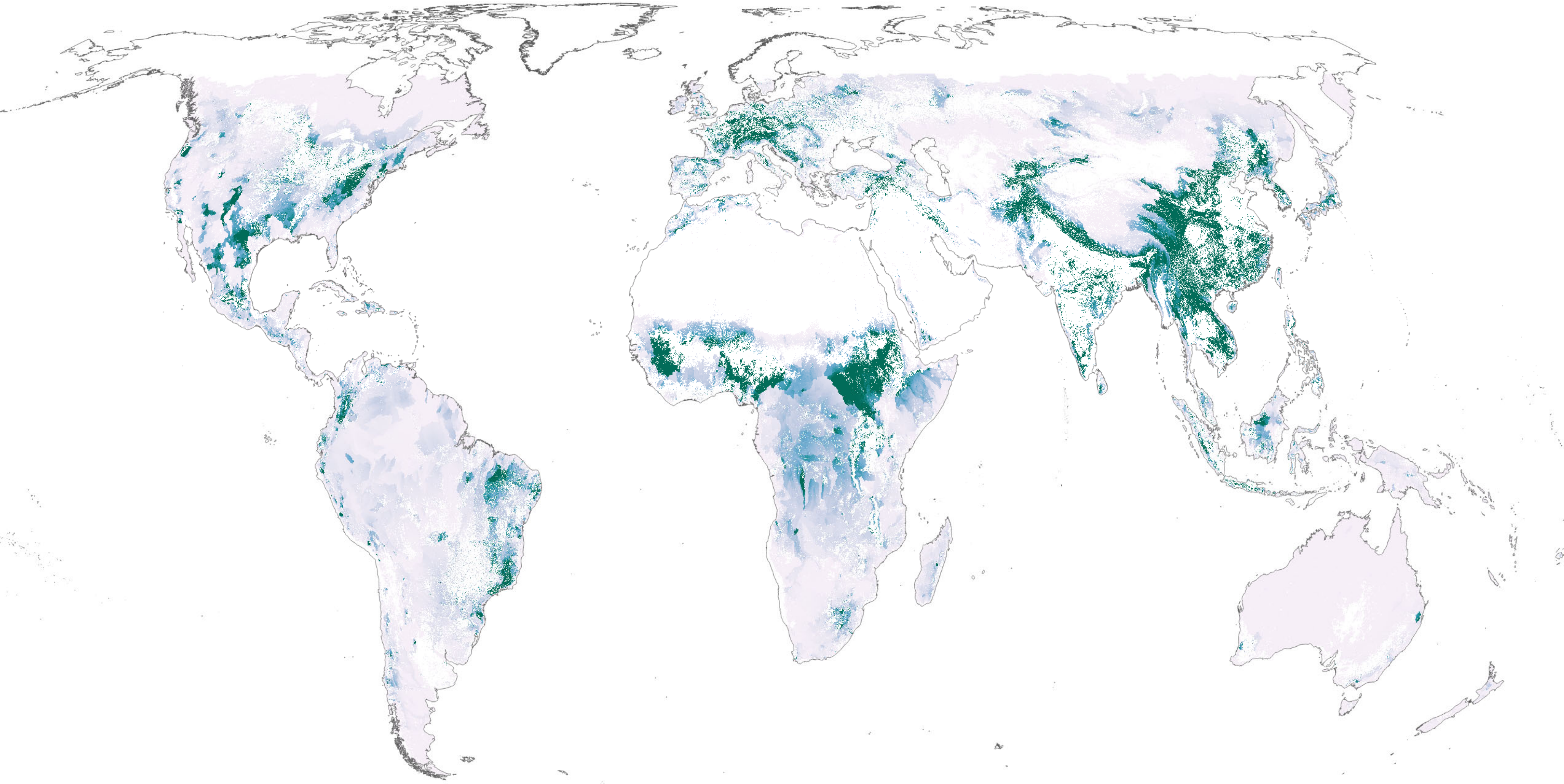

Nitrogen retention for water quality regulation

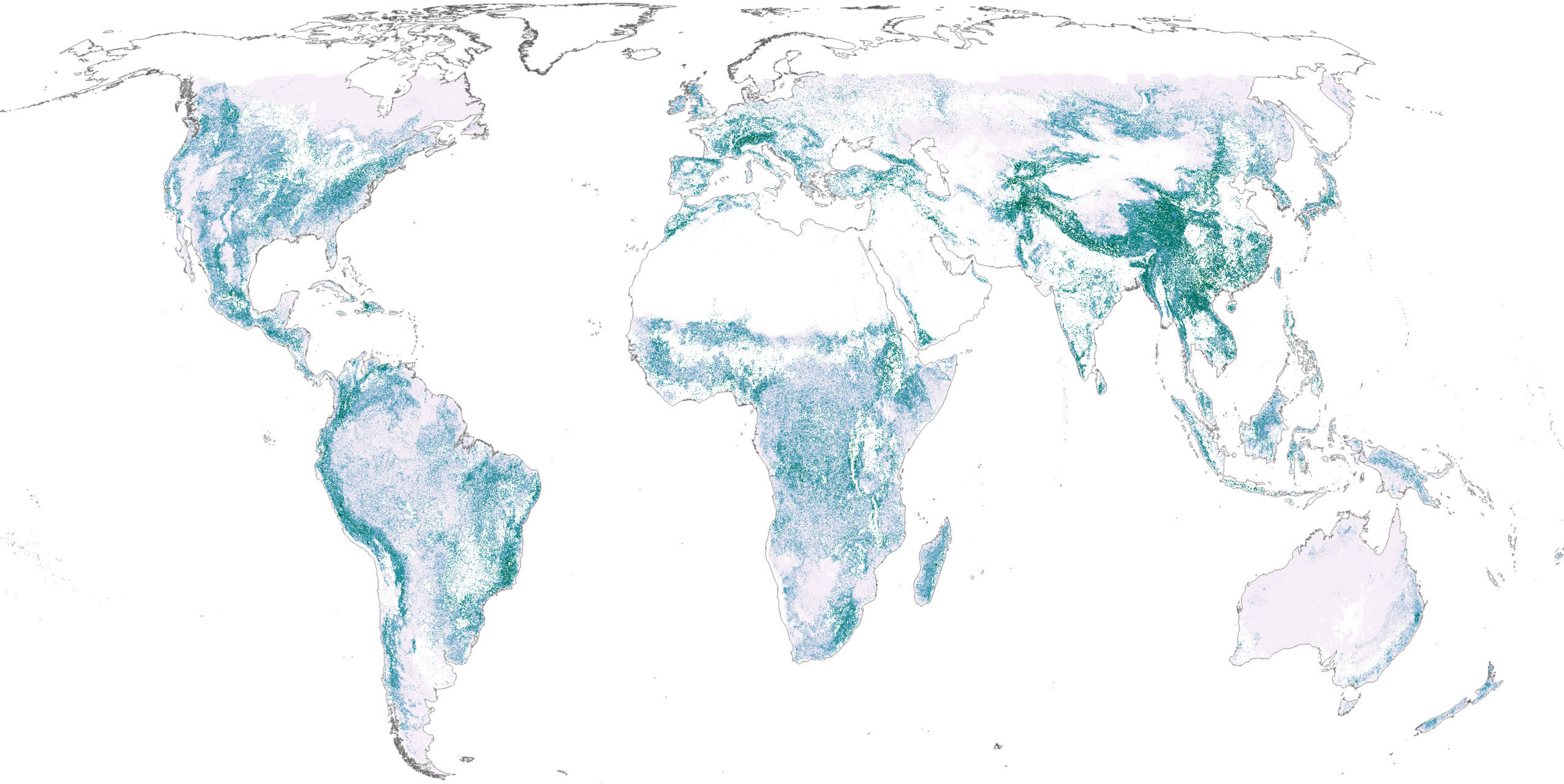

Sediment retention for water quality regulation

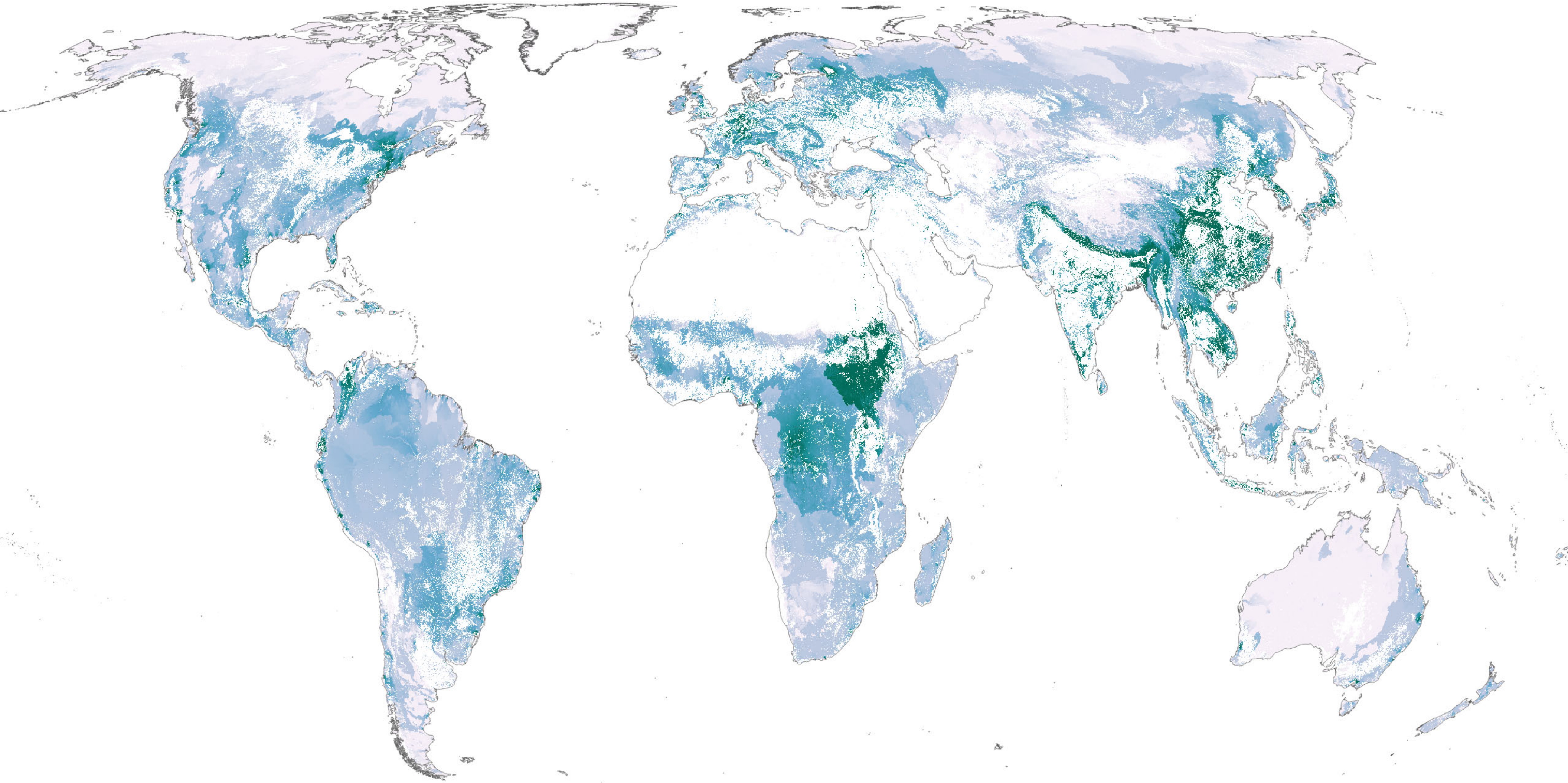

Flood regulation

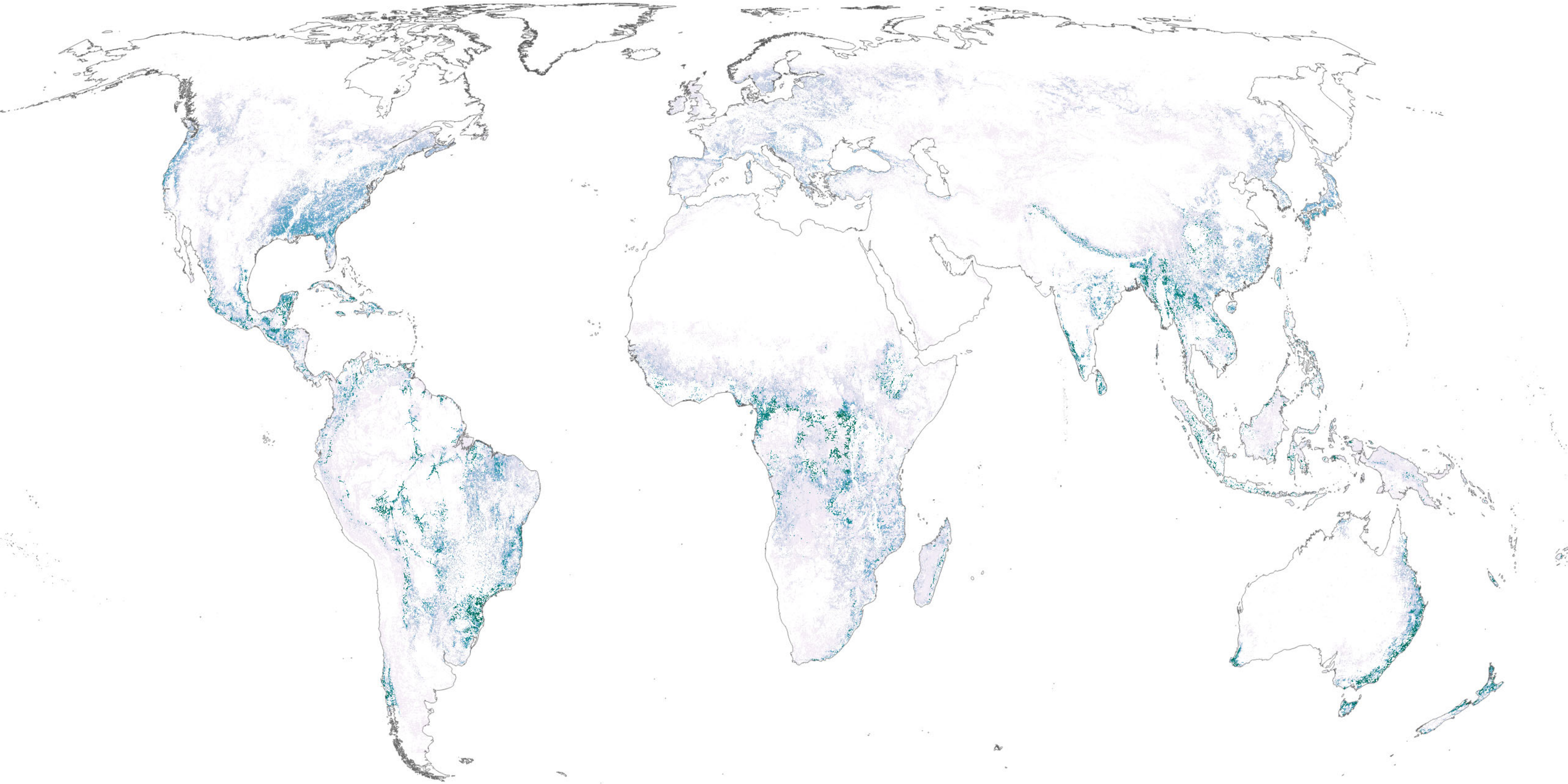

Timber production

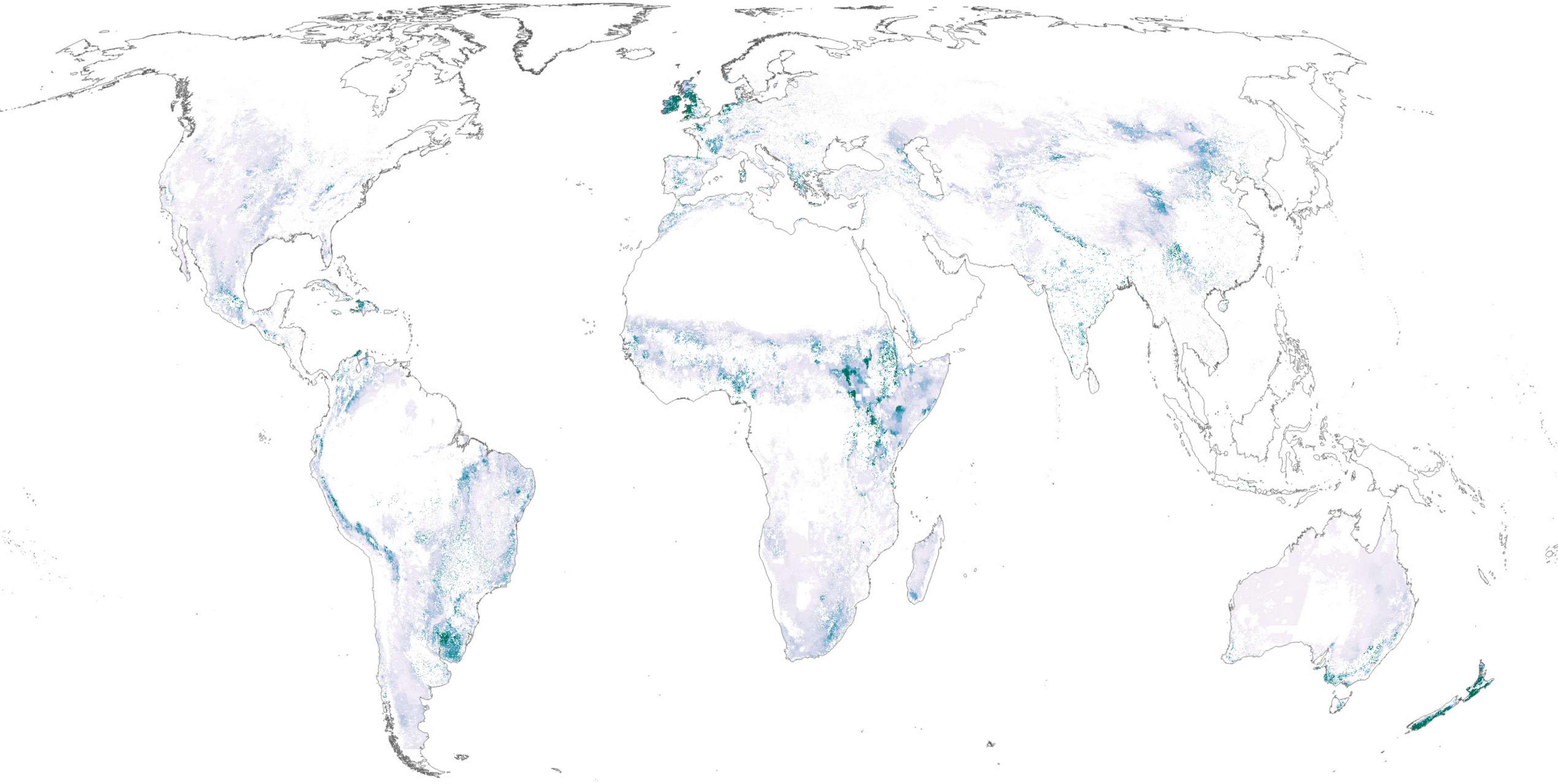

Fodder production for livestock

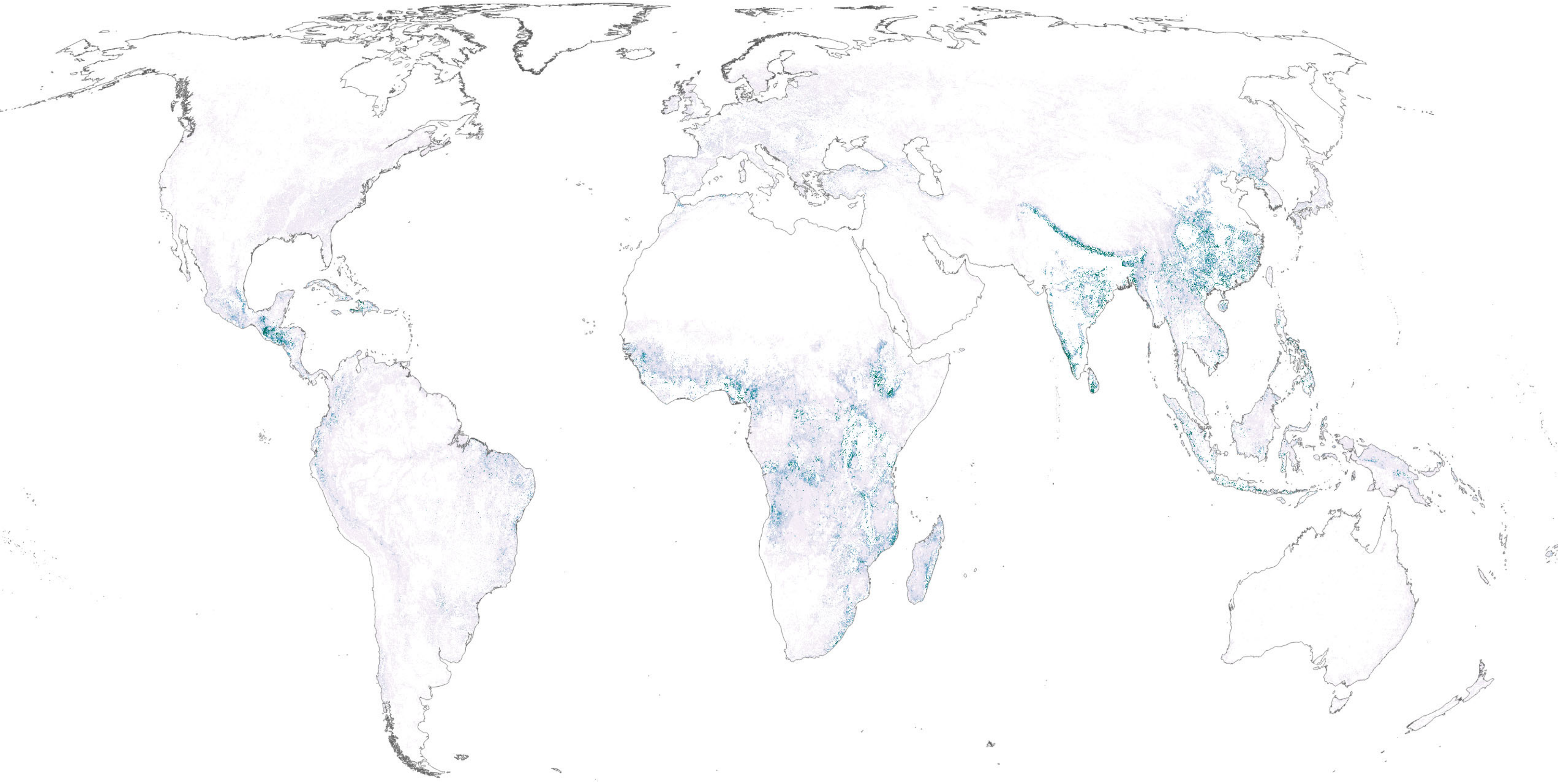

Fuelwood production

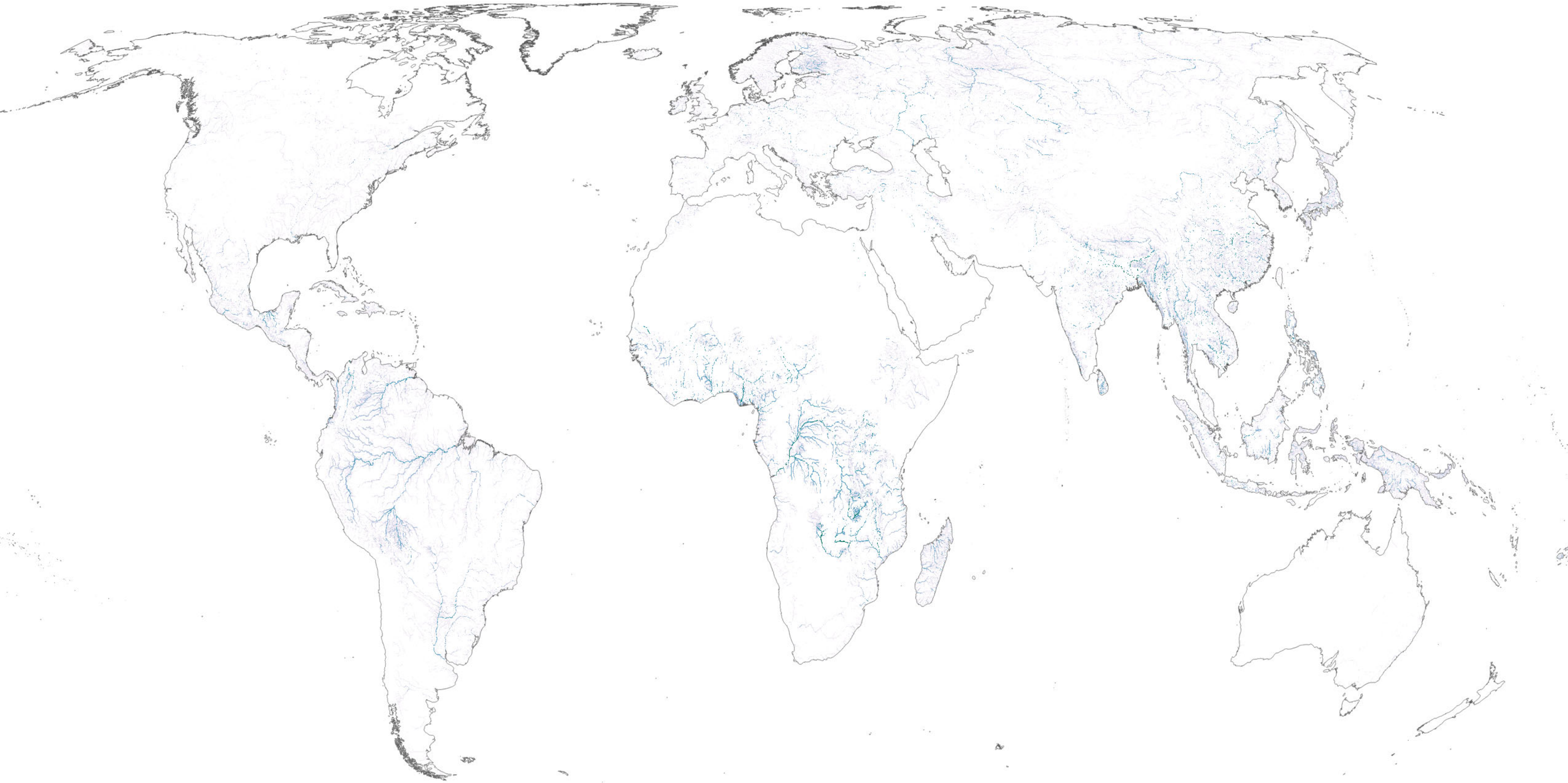

Riverine fish harvest

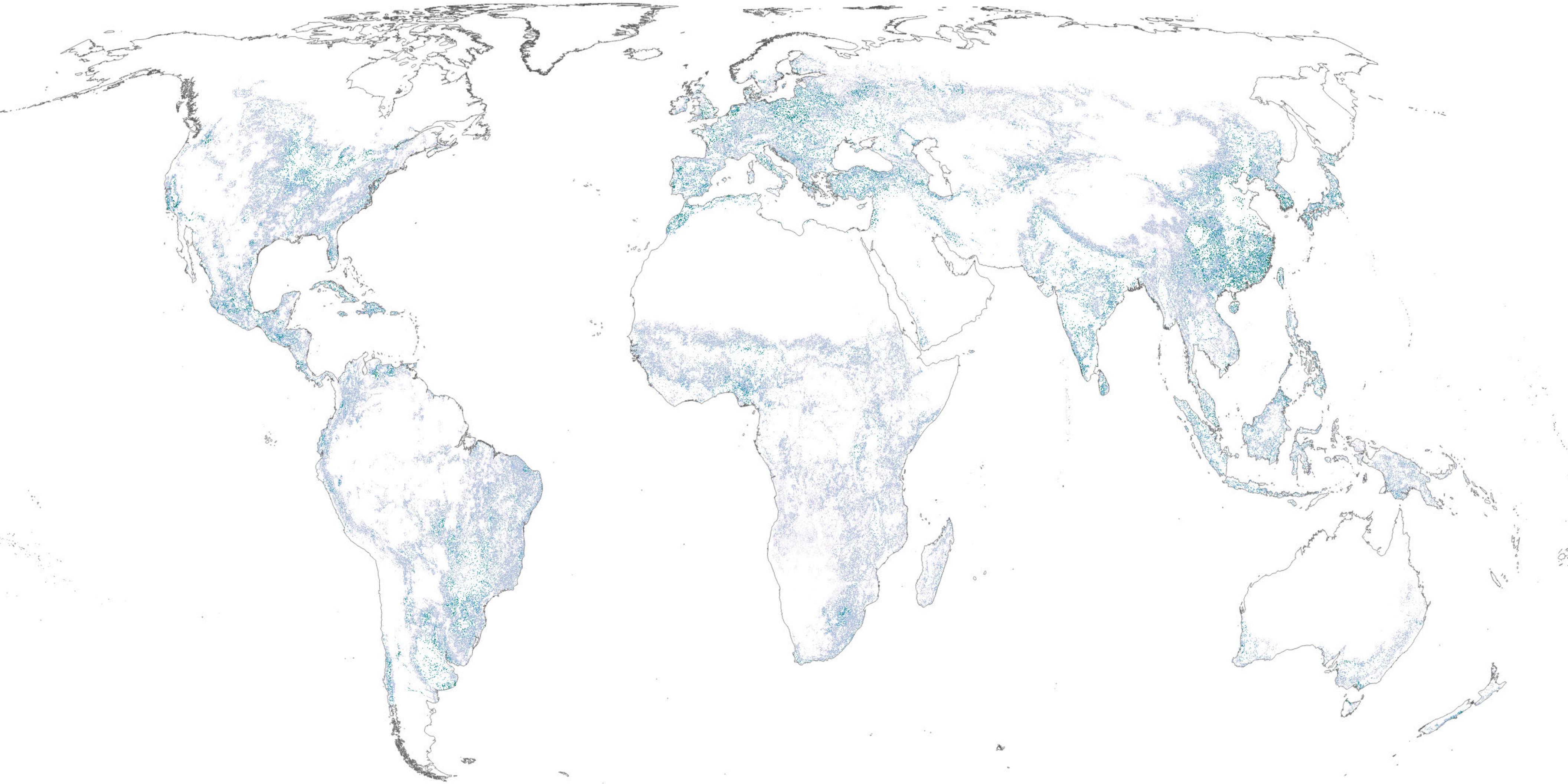

Crop pollination

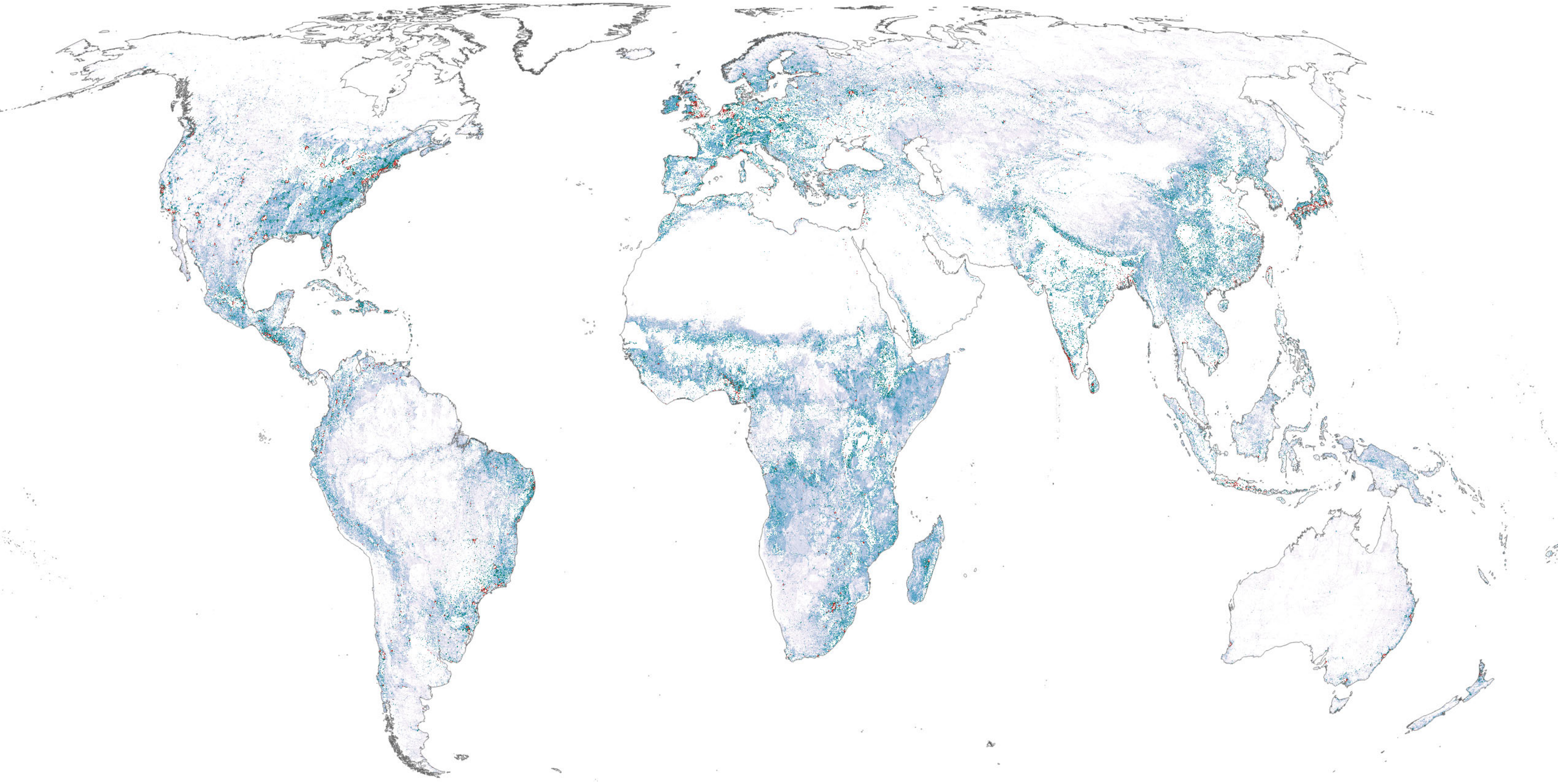

Access to terrestrial nature (recreation, gathering)

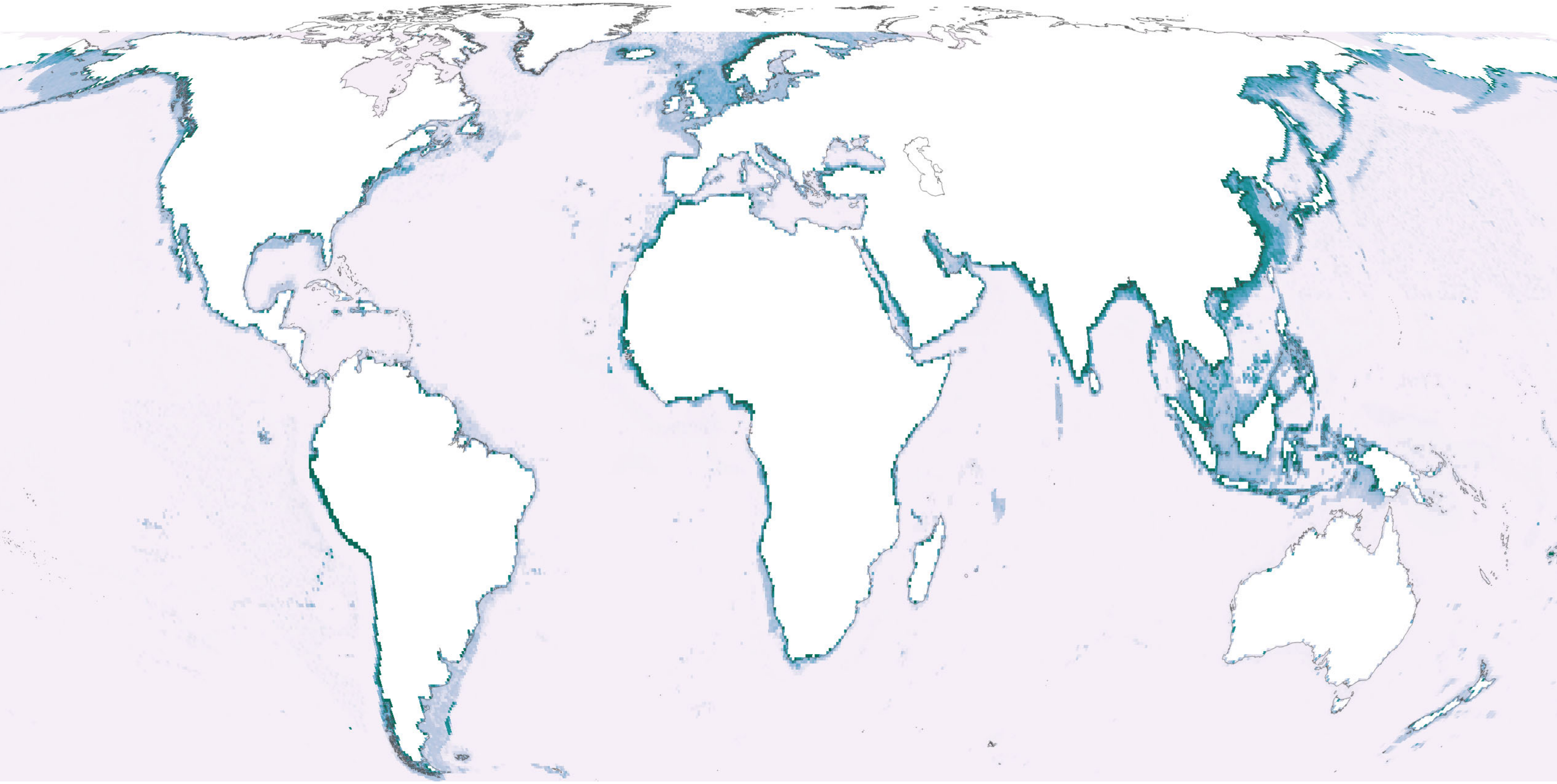

Marine fish harvest

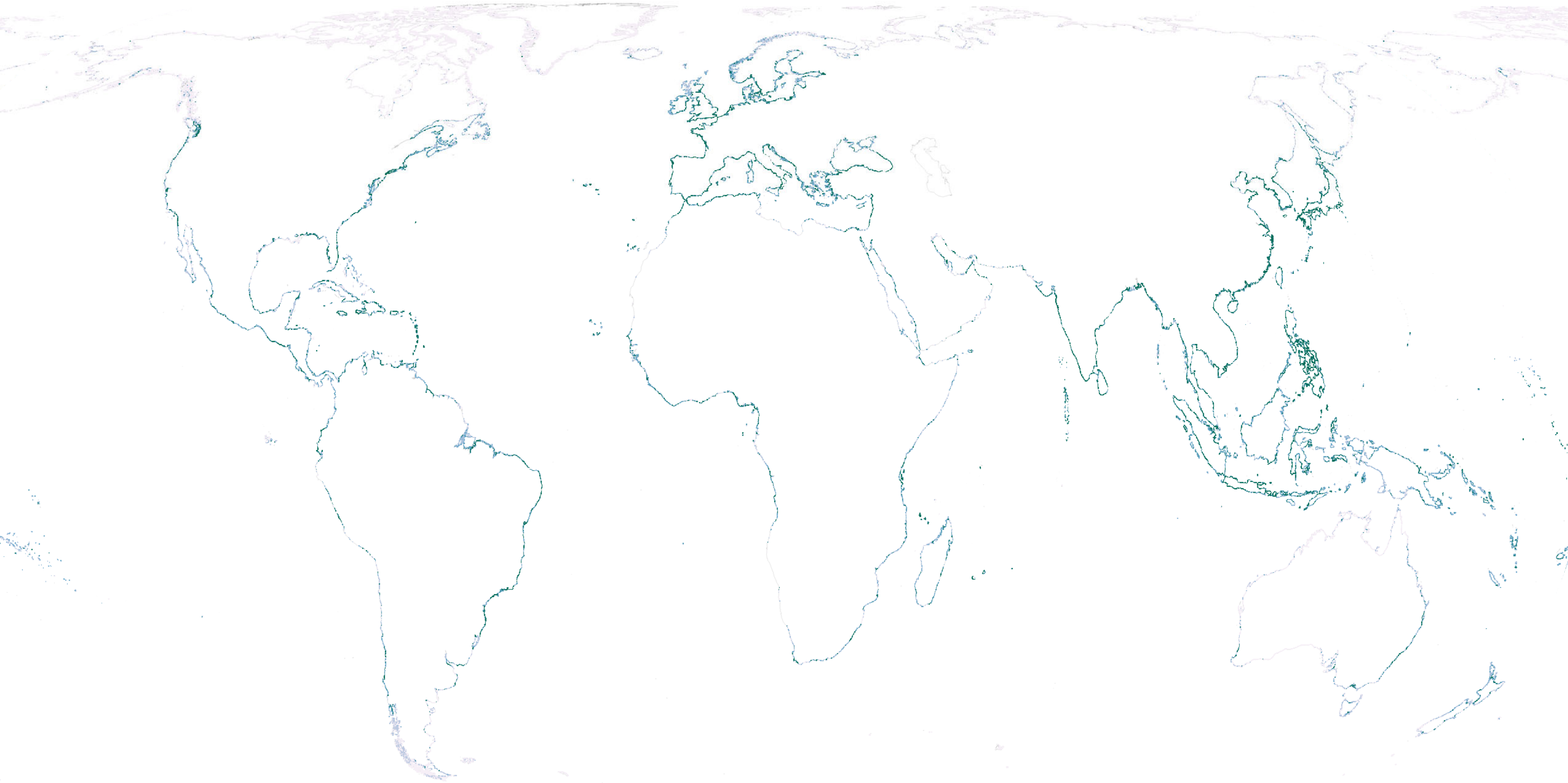

### Coastal risk reduction

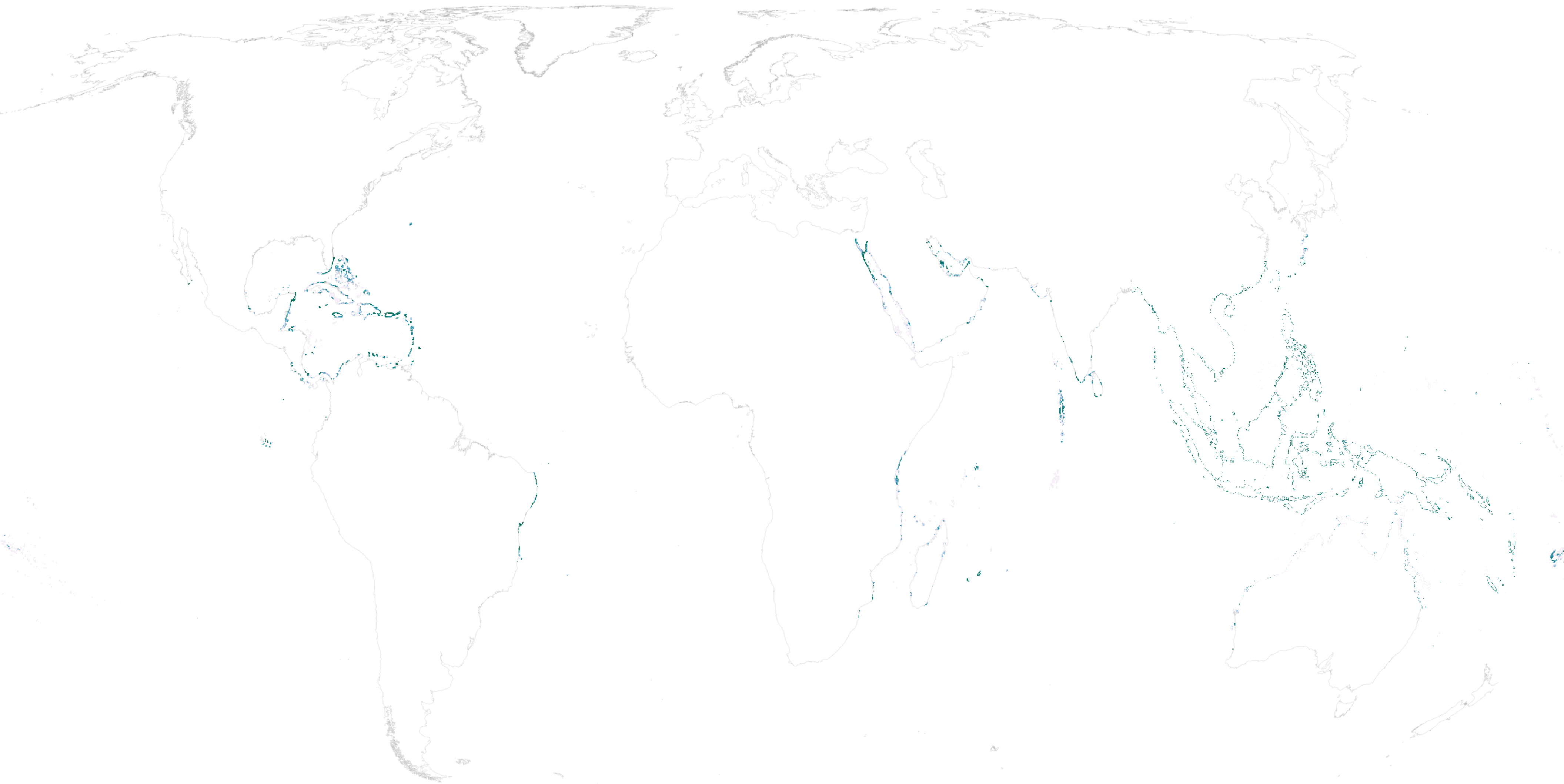

### Coral reef tourism

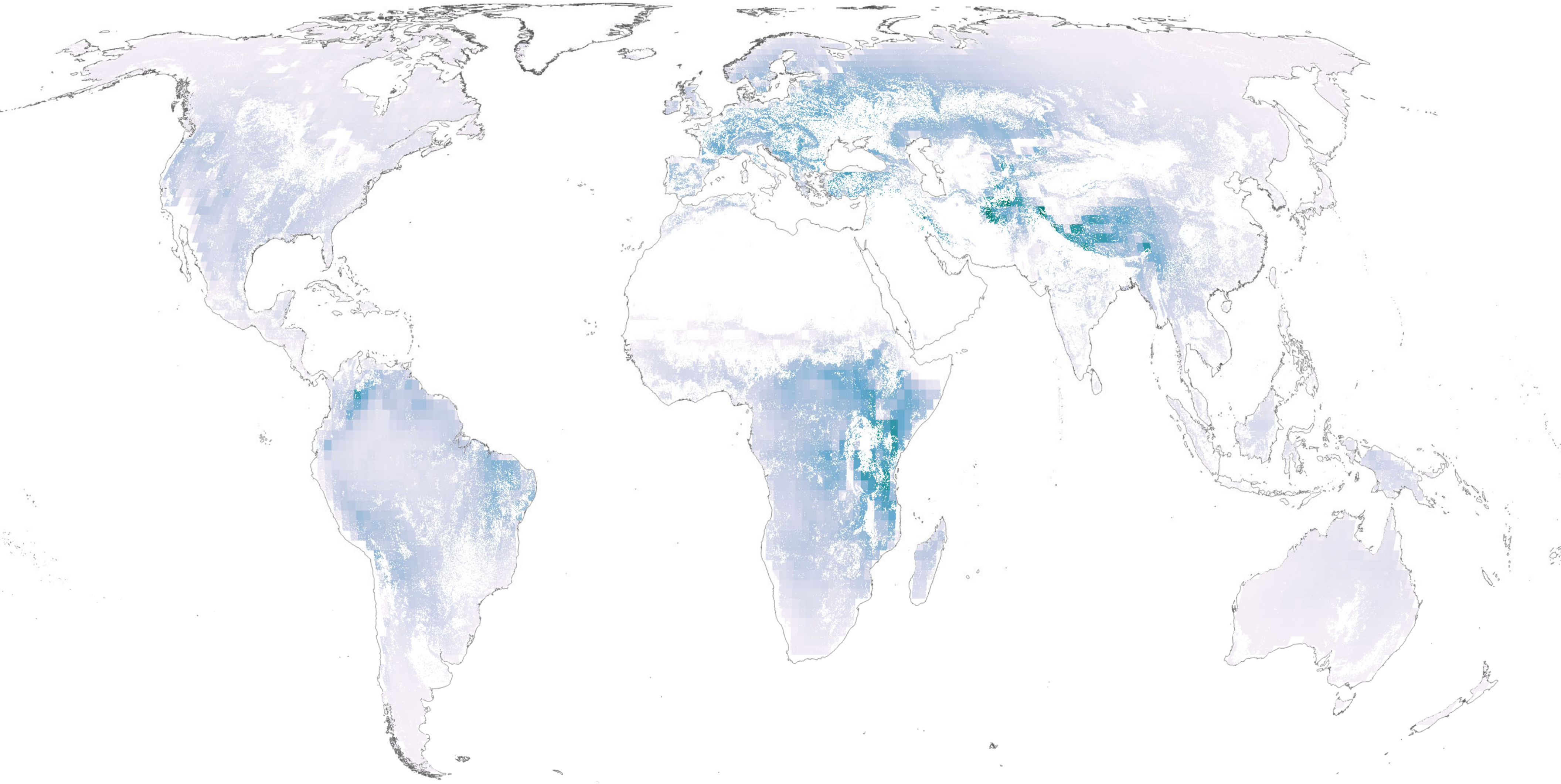

Atmospheric moisture recycling

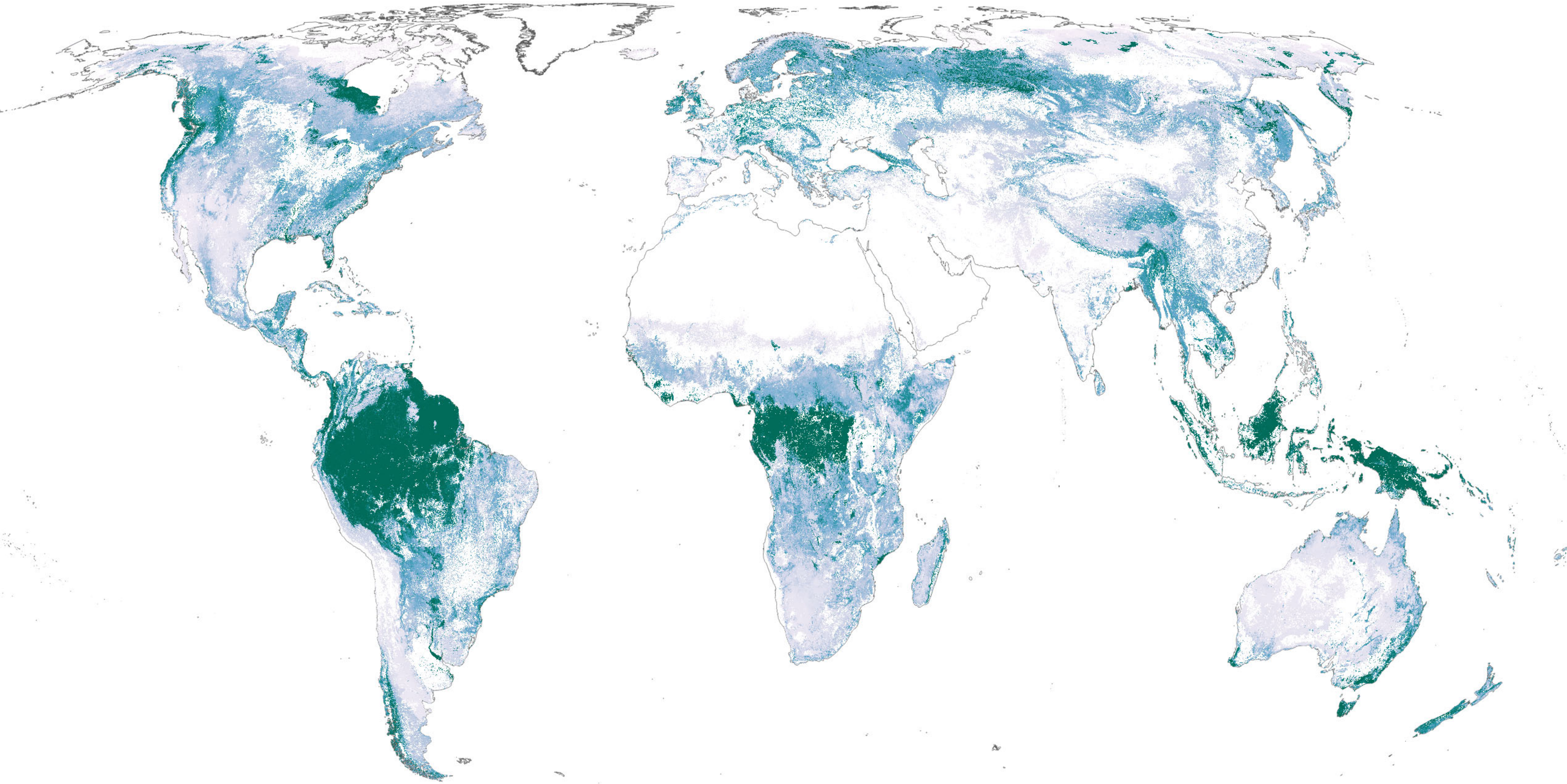

Vulnerable terrestrial ecosystem carbon storage

**Extended Data Figure 2 | Percent of land in critical natural assets (CNA) for local (a) and global (b) benefits, plotted against the percentage of total natural assets in a country**

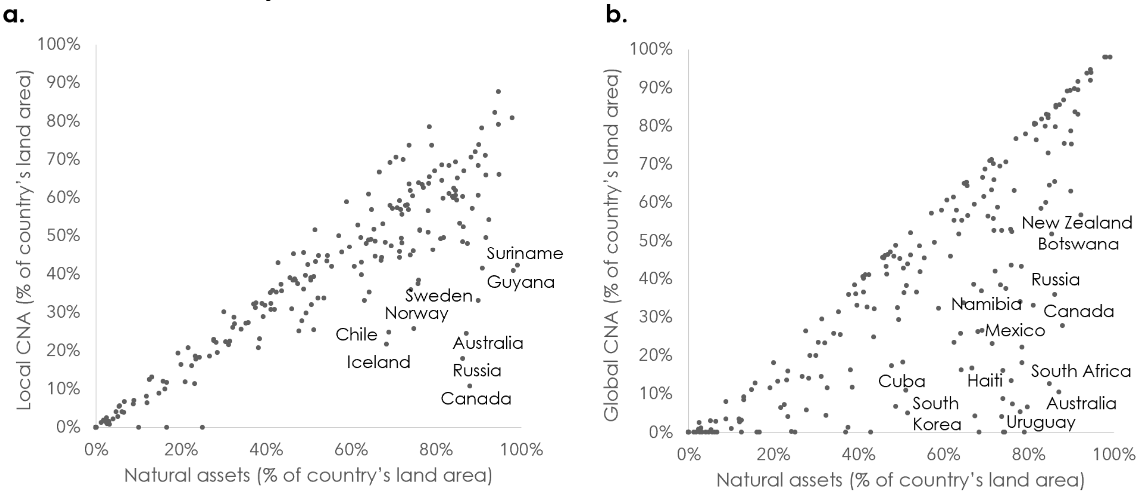

**Extended Data Figure 3 | Spatial congruence between NCP aligning with critical natural asset hotspots.**

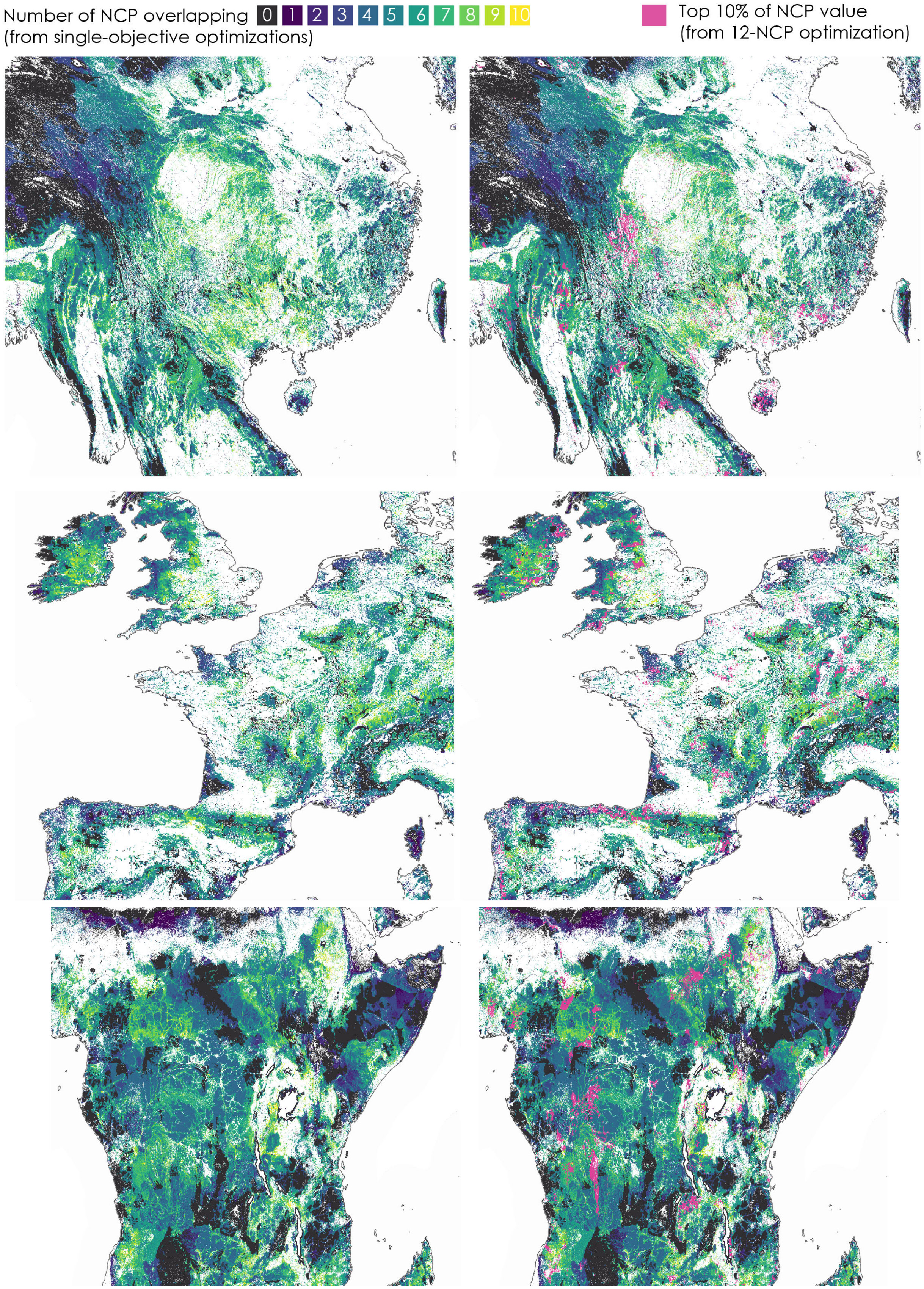

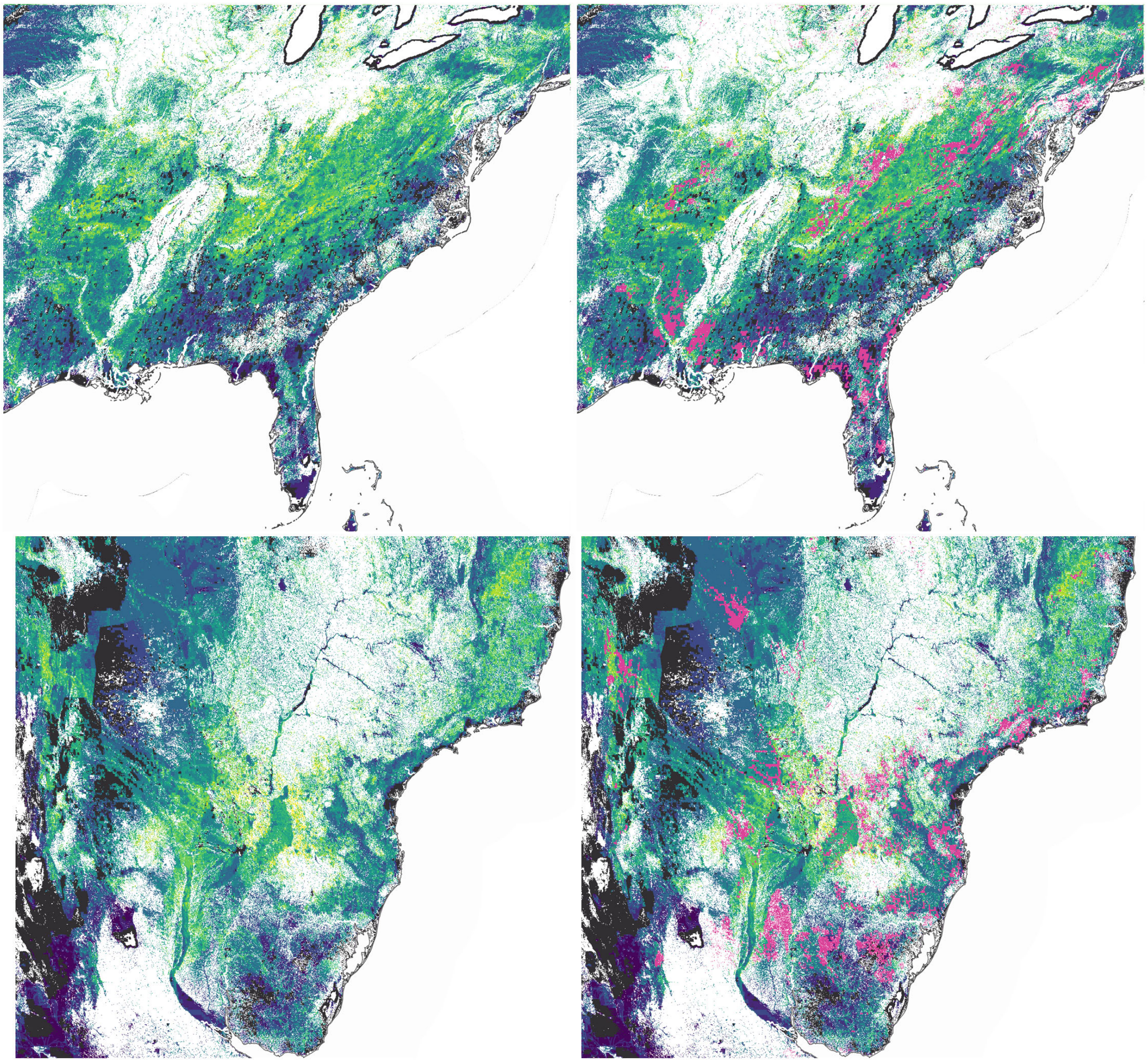

Number of NCP overlapping 0 1 2 3 4 5 6 7 8 9 10
(from single-objective optimizations)

Top 10% of NCP value
(from 12-NCP optimization)

**Extended Data Figure 4 | Critical natural assets identified through optimization at the global level of two climate-relevant NCP: vulnerable carbon and vegetation-mediated atmospheric moisture regulation.** As in Fig. 1, the NCP accumulation curve reflects the total area required to maintain target levels of both NCP (but in this case globally, not within each country), with dotted lines denoting the area of critical natural assets (90% of global climate NCP in 39% of land area). The map shows critical natural assets for global climate NCP, with darker shades connoting greater contribution to aggregate NCP.

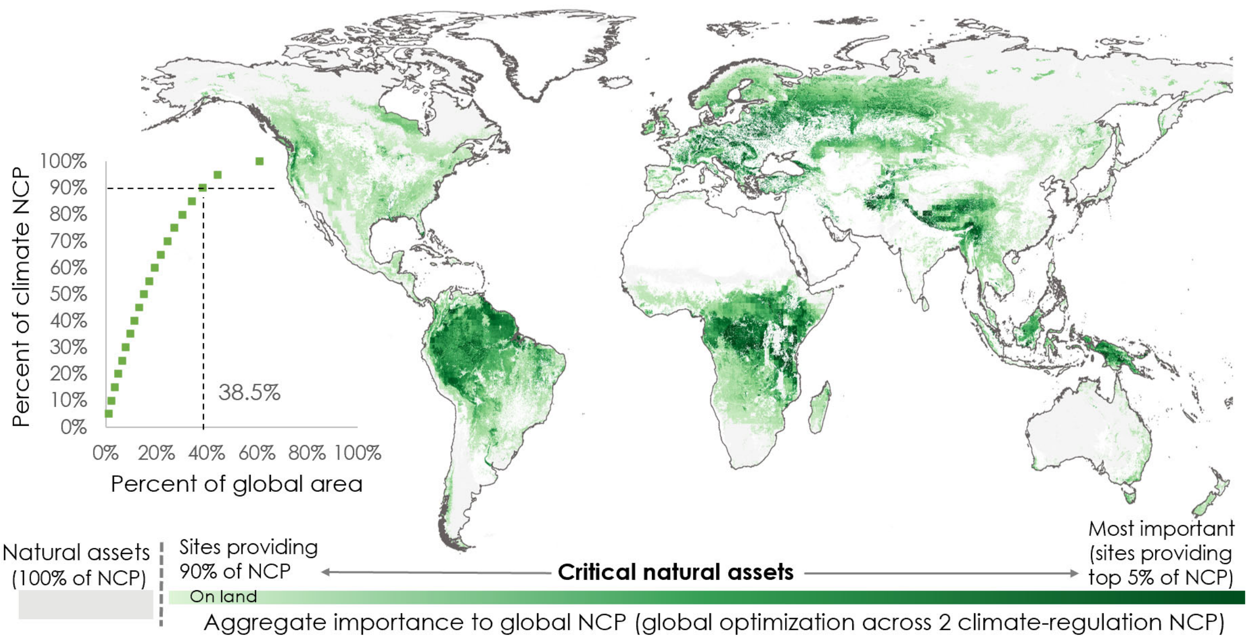


**Extended Data Figure 5 | Critical natural assets identified through optimization at the global level of 12 "local" NCP.** As in Fig. 1, the NCP accumulation curve reflects the total area required to maintain target levels of both NCP (but in this case globally, not within each country), with dotted lines denoting the area of critical natural assets (90% of the 12 NCP listed in Fig. 1a in 22% of land area and 13% of EEZ areas). The map shows critical natural assets for global climate NCP, with darker shades connoting greater contribution to aggregate NCP.

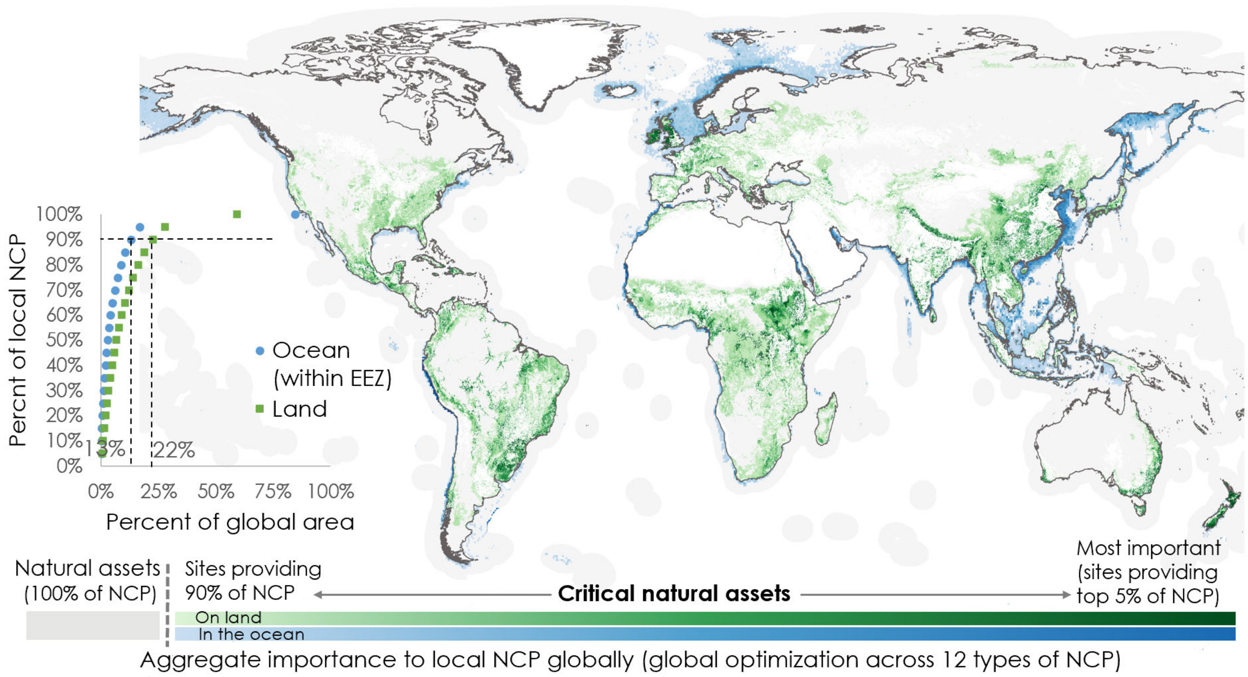
