## Supplementary material for "Mapping the planet’s critical natural assets": SI Methods

#### Mapping the planet's critical natural assets for people

### Materials and Methods

*Summary* 3
*Key terms* 3
"Local" NCP 4
1. 4
    Nitrogen retention 4
    Downstream beneficiaries 5
2. 7
    Sediment export 8
    Model changes necessary to calculate per-pixel sediment "retention" 8
    (sediment downslope deposition) 8
    Downstream beneficiaries 10
3. 10
    Pollination on-farm 10
    Equivalent people fed 10
    Mapping back to habitat 11
4. 11
5. 12
  (a) Commercial timber 13
  (b) Domestic timber 13
6. 14
7. 15
8. 15
      Computational Method 16
9. 17
10. 17
11. 18
12. 19
    Inclusion of new data 19
    Projecting coastal risk reduction value back onto habitat 20
Global climate NCP 20
  13. 21

14. 22
Diversity 23
15. 24
16. 24
Additional datasets 24
17. 25
18. 25
19. 26
20. 26
21. 27
22. 27
Code repository 26
Targets 27
Global and country-by-country optimization 27
Optimization objectives 27
23. 30
24. 31
Effect of masking and resolution on percent of natural habitat 30
No data values 30
Skewed and zero-inflated data 30
Outliers 31
Philosophical considerations 31

#### *Summary*

In this supplemental methods text, we describe the methods used to create the 12 local nature's contributions to people (NCP) included in our spatial prioritization (Extended Data Table 1), as well as the global (climate-related) NCP and additional global benefits (diversity) included in the subsequent overlap analysis (Extended Data Table 2), along with supporting datasets (Extended Data Table 2). For each of the NCP, we describe whether they were masked to focus on specific habitats (Extended Data Table 3) and whether they were multiplied by the relevant beneficiary population in order to attribute the realized value of the NCP to the natural assets (ecosystems) providing it. Then we discuss the optimization algorithm used in the spatial prioritizations and subsequent analyses, and highlight some of the limitations and caveats of this approach.

#### *Key terms*

We follow the definition given by the Intergovernmental Science-Policy Platform for Biodiversity and Ecosystem Services (IPBES) for **"Nature's Contributions to People" (NCP),** the contributions of living nature (i.e., diversity of organisms, ecosystems, and their associated ecological and evolutionary processes) to the quality of life for people. This differs from the framework presented in Chaplin-Kramer et al. (2019), where NCP was calculated as the proportional contribution of nature to meeting people's needs, which was important because both nature's contributions and people's needs were changing in the context of the scenarios explored there. However, we do try to maintain the combination of "nature's contributions" and "people's needs" in this approach, by mapping the benefits and/or beneficiaries to natural and semi-natural habitats. In many of the models for individual NCP presented below, they are referred to as ecosystem services, often split into "biophysical service" or "potential service" to represent **nature's contribution** and delineating a "beneficiary" for that particular contribution to represent the **people** in NCP. The realized service mapped to natural habitat as NCP here is either the potential service multiplied by the number of people benefitting (e.g., coastal risk reduction multiplied by number of people within the protective distance of the habitat, nitrogen retention multiplied by the number of people downstream of the habitat) or the final end benefit to an indeterminate or global beneficiary (e.g., equivalent number of people fed by wild pollination-derived crop production attributed to the habitat providing the pollinator).

### "Local" NCP

#### 1. Nitrogen retention for water quality regulation

Fertilizers like nitrogen are a major source of pollution to freshwater systems and drinking water. However, some may be retained by healthy ecosystems, regulating water quality in streams. This can be modeled to show how much nitrogen retention can be provided by nature (see Chaplin-Kramer et al. 2019 for more detail). The people benefitting from nitrogen retention are those who would otherwise be exposed to nitrogen contamination in their drinking water. In this analysis, the number of people downstream were calculated for every pixel of habitat, to provide a sense of which habitat potentially benefits the most people. Ideally, to map realized nitrogen retention, we would be able to convert biophysical service production into a measure of change in well-being, whether monetary, in health terms, or otherwise. However, the state of the science and data available globally precludes this for most services, so our proxy was the number of people downstream who could potentially benefit from that retention. NCP for nitrogen retention is, therefore, expressed as nitrogen retention on natural and semi-natural pixels multiplied by the number of people downstream of those pixels.

**Nitrogen retention**

Nitrogen retention is modeled using the InVEST Nutrient Delivery Ratio model (see (http://data.naturalcapitalproject.org/invest-releases/3.5.0/userguide/ndr.html for more detail), adapted for global runs using the methods described in Chaplin-Kramer et al. 2019 (section 3.1 of the Supplementary Methods), with the exception of the Digital Elevation Model (DEM) used. In this analysis, we use the 3s void-filled DEM from SRTM (Shuttle Radar Topography Mission) (doi:/10.5066/F7F76B1X), and watersheds were pre-delineated from the Hydrosheds (Lehner et al. 2008) dataset. Watersheds in the DEM must hydrologically drain in order to accurately model the accumulated nutrient deposition along a downstream path. However, errors in the DEM as well as small real-world depressions can dramatically alter the direction of flow. In an extreme case we found our hydrological model split the Nile into two rivers, one flowing "backwards" on the path that is a real world drain to the Toshka lakes. To handle this and similar cases we forced large watersheds (arbitrarily defined as 1 square degree) to have a single outlet defined at its lowest DEM value in the watershed. We found that smaller watersheds could reasonably have multiple outlets due to the difference in accuracy between SRTM and the HydroSHEDS polygon data so we did not force single outlets on smaller watersheds. Also note, SRTM only extends from 60 degrees N to 56 degrees S, so the model is limited to the sub-arctic region. Future efforts could fuse the STRM product with a new arctic DEM (at 2 m resolution) for full global coverage (Porter et al. 2018).

Because we are focused on the contribution of natural assets, we mask nitrogen retention to natural and semi-natural European Space Agency (ESA 2015) (including natural/crop mosaic) classes 30-180 (see Extended Data Table 3). While we include cropland and all other “non-natural” land-uses in the modeling of nitrogen export that could arrive on a natural or semi-natural pixel, we remove these classes from the NCP map to be consistent with other layers, and in acknowledgement of our uncertainty in land management in agricultural, urban and bare areas affecting nitrogen retention.

**Downstream beneficiaries**

We use population data from Landscan 2017 (Extended Data Table 2) to represent the number of people that may benefit from an upstream hydrological flow into each ESA land cover pixel (resolution: 10 arc-sec, ~300 m). The total number of downstream people who benefit from any given upstream pixel can be directly calculated by starting at that pixel and walking downstream. At each downstream step, the number of people in the current pixel is added to a running total. When the last step terminates by hitting a body of water or the edge of the raster, the final count is the number of people who are in the downstream flow path that can be written to the Downstream beneficiary raster at the originating step.

In practice, it is more efficient to start at the terminating edge of the flow and proceed upstream. At each upstream step the running total is added to the current pixel and written to the Downstream beneficiary raster at the same time. This allows the calculation of the entire Downstream beneficiary raster in one pass but requires extra bookkeeping to calculate originating drains and track upstream flow branching.

As a one-dimensional example, take a flow direction starting from the left (upstream) to the right (downstream). If the population along each cell is represented as the following:

*Beneficiary raster (input)*

| | | | | | | | | | | | |
|---|---|---|---|---|---|---|---|---|---|---|---|
| 10 | 20 | 15 | 9 | 7 | 9 | 83 | 1000 | 9 | 1 | 78 | 273 |

Then the computed beneficiary raster is the following (each cell represents the number of people downstream from that pixel):

*Downstream beneficiary raster (output)*

| | | | | | | | | | | | |
|---|---|---|---|---|---|---|---|---|---|---|---|
| 1514 | 1504 | 1484 | 1469 | 1460 | 1453 | 1444 | 1361 | 361 | 352 | 351 | 273 |

However, this direct measurement of the number of people along a flowpath can yield artificially high values in large watersheds simply because they have long flowpaths. An extreme example is the Nile River watershed, which accumulates millions of people on pixels thousands of kilometers upstream from Cairo.

Qualitatively, beneficiaries that are closer to the point of service provision will benefit more and therefore should “count” more than those further down on the flowpath. Analogously, beneficiaries that are a Very Long distance downstream should contribute minimally or not at all to the number of beneficiaries on the flowpath for a given pixel at all.

To “weight” the importance of nearby beneficiaries more heavily than distant beneficiaries, we sum all downstream beneficiaries as described above but scale the number of distant beneficiaries, using the distance half-life of the expected benefit as the scaling term. For example, if the benefits of a given service were expected to dissipate by half within 100 meters, beneficiaries 100 meters from the source would be scaled by 0.5. Thus, the beneficiaries at any point p are defined as the sum of the half-life scaled per-pixel beneficiaries on the downstream flowpath. Formally, the number of weighted downstream beneficiaries on a pixel p can be calculated as

$$B(p) = \sum_{i \in \{\text{[illegible]}\}} b(i) \cdot \left(\frac{1}{2}\right)^{d(p,i)/d_{hl}}$$

where

- *B(p)* are the number of beneficiaries calculated to be at point *p*
- *b(i)* is the number of people present at point *i* that could count as beneficiaries upstream
- *d(p,i)* is the flowpath distance between point *p* and *i*, and
- $d_{hl}$ is the half-life distance used to scale *b(i)* by half when $d(p,i)=d_{hl}$.

For the main analysis, we attenuate the importance of distant beneficiaries by setting the half-life to 500 km, but we also explore the sensitivity to this scaling factor by including an additional layer with a half-life of 50 km. These different scales do not have a major impact on the overall optimization, although they do differ when hydrologic NCP are optimized individually (see Section 22; Extended Data Table 5, row 1 vs. row 7 and rows 28 and 30).

*Benefiting areas mask*

We generate a “benefiting areas” raster mask to account for the population that benefits from upstream services. This raster mask contains values of “1” where there is a pixel on an upstream flowpath that contains a service pixel and “0” if no upstream pixel contains a service. Once this mask is generated we use it to filter a population raster such that pixels that lie outside of the benefiting area (i.e. “0” pixels on the benefiting area mask) are set to 0.

As a one-dimensional example, take a service area raster with a flow path starting from the left (upstream) to the right (downstream):

<u>Service Area Mask (input)</u>

| | | | | | | | | | | | |
|---|---|---|---|---|---|---|---|---|---|---|---|
| 0 | 0 | 0 | 0 | 1 | 0 | 0 | 1 | 0 | 0 | 0 | 0 |

The computed benefiting area mask is the following (a cell is “1” if any upstream flowpath cell is a service area):

<u>Benefiting areas mask raster (output)</u>

| | | | | | | | | | | | |
|---|---|---|---|---|---|---|---|---|---|---|---|
| 0 | 0 | 0 | 0 | 1 | 1 | 1 | 1 | 1 | 1 | 1 | 1 |

Once the benefiting areas mask raster is created, it can be used to mask out a population raster to show only the population that benefits from any upstream service area. Continuing the 1D example, if the base population is as follows:

| | | | | | | | | | | | |
|---|---|---|---|---|---|---|---|---|---|---|---|
| 358 | 565 | 148 | 126 | 914 | 423 | 402 | 549 | 691 | 415 | 991 | 60 |

Then the benefiting population is created by filtering any population with “0” pixels in the benefit area mask raster:

| | | | | | | | | | | | |
|---|---|---|---|---|---|---|---|---|---|---|---|
| 0 | 0 | 0 | 0 | 914 | 423 | 402 | 549 | 691 | 415 | 991 | 60 |

We use this technique to generate a downstream mask to mask the populations in critical natural asset areas selected by the optimization routines described in Section 22.

### 2. Sediment retention for water quality regulation

Erosion causes issues for land degradation and water quality, with sediments clogging waterways and often carrying diseases that can lead to water-borne illness. Here we model sediment retention provided by vegetation by adapting the InVEST Sediment Delivery Ratio (SDR) model, which maps overland sediment generation and delivery to the stream. Ideally this would be delineated for reservoirs, irrigation canals, or other water delivery infrastructure that is most impacted by sedimentation, but lacking a comprehensive global dataset identifying all such infrastructure, we again use the proxy of number of people downstream. Nature’s contribution to people for sediment retention is, therefore, expressed as sediment retention on natural and semi-natural pixels multiplied by the number of people downstream of those pixels.

**Sediment export**

The potential for erosion is determined by climate (specifically rain intensity), soil properties, topography, and vegetation. Land use practices can impact the amount of erosion and the amount of sediment that can be retained by vegetation before it reaches a stream. The magnitude of this effect is primarily determined by: i) the main sediment sources (vegetation has a smaller effect in catchments where sediments are not primarily coming from overland flow); and ii) the spatial distribution of sediment sources and sinks (vegetation has a larger effect if positioned between sediment sources and the stream). The InVEST SDR model is a spatially-explicit model working at the spatial resolution of the DEM raster. For each pixel (mapped here using ESA 2015; 10 arc-sec, ~300 m resolution), the model first computes the amount of annual soil loss from that pixel, then computes the SDR, the proportion of soil loss actually reaching the stream. The amount of annual soil loss from a pixel (in tons·ha−1yr−1), is given by the revised universal soil loss equation (RUSLE1), and the SDR for a pixel is then derived from a conductivity index based on the upslope and downslope areas of that pixel. See http://releases.naturalcapitalproject.org/invest-userguide/latest/sdr.html for more detail.

**Model changes necessary to calculate per-pixel sediment “retention” (sediment downslope deposition)**

This is retention in terms of the amount of total sediment that gets deposited on a pixel from on-pixel and upstream Universal Soil Loss Equation (USLE) sources.

As described above, sediment export *E* to stream from pixel *i* is defined as:

$$E_i = USLE_i \cdot SDR_i$$

In turn, there is a quantity that is the remainder of USLE that does not reach the stream. This sediment load must be deposited somewhere on the landscape along the flowpath to the stream:

$$E'_i = USLE_i \cdot (1 - SDR_i)$$

We can think about these two quantities as separate flows. Due to the nature of the calculation of SDR, the quantity $E_i$ has accounted for the downstream flow path and biophysical properties that filter sediment to stream. This means we can exclusively model the flow of E’ downstream independently of the flow of $E_i$.

To do this, we state the following properties about how $E'_i$ and SDR behave across a landscape:

- **Property A: SDR monotonically increases along a downhill flowpath:** As a flowpath is traced downhill, the value of SDR will monotonically increase since the amount of downstream flow distance decreases. Note there is the numerical possibility that a downstream pixel has the same SDR value as an upstream pixel. The implication in this case is that no on-pixel sediment flux deposition occurs along that step.

- **Property B: All non-exporting sediment flux on a boundary stream pixel is retained by that pixel:** If pixel *i* drains directly to the stream there is no opportunity for further downstream filtering of $E'_i$. Since $E'_i$ is the inverse of $E_i$, the implication is that the upstream flux (defined as $F_i$ *below)* must have been deposited on the pixel.

Given these two properties, the amount of $E'_i$ retained on a pixel must be a function of:

1) the absolute difference in SDR values from pixel *i* to the downstream pixel(s) drain, and
2) how numerically close the downstream SDR value is to 1.0 (the stream pixel).

These mechanics can be captured as a linear interpolation of the difference of pixel *i*'s SDR value with its downstream SDR counterpart with respect to the difference of pixel *i*'s difference with a theoretical maximum downstream SDR value 1.0. Formally,

$$dR_i = ([\sum_{k \in \{directly\ downstream\ from\ i\}} SDR_k \cdot p(i,k)] - SDR_i)/(1.0 - SDR_i)$$

The 'd' in 'dRi' indicates a delta difference and *p(i,k)* is the proportion of flow from pixel *i* to pixel *j*. This notation is meant to invoke the feeling of a derivative of $R_i$. Note the boundary conditions are satisfied:

- In the case of **Property A** (downstream $SDR_k = SDR_i$), the value of $dR_i = 0$ indicating no $F_i$ will be retained on the pixel.
- In the case of **Property B** (downstream $SDR_k = 1$ because it is a stream) the value of $dR_i=1$ indicating the remaining $F_i$ is retained on the pixel.

Now we define the amount of sediment flux that is retained on any pixel in the flowpath using $dR_i$ as a weighted flow of upstream flux:

$$R_i = dR_i \cdot (\sum_{j \in \{pixels\ that\ drain\ to\ i\}} F_j \cdot p(i,j) + E'_i)$$

where $F_j$ is the amount of sediment-export-that-does-not-reach-the stream "flux", defined as:

$$F_i = (1 - dR_i) \cdot ((\sum_{j \in \{pixels\ that\ drain\ to\ i\}} F_j \cdot p(i,j) + E'_i)$$

As described for nitrogen retention, the sediment retention on each pixel is then masked to all natural and semi-natural habitats (ESA classes 30-180; see Extended Data Table 3), though we include all land-uses (cropland, barren areas, etc.) in the modeling of sediment export that could arrive on a natural or semi-natural pixel.

**Downstream beneficiaries**

We follow the same process as described for nitrogen retention (Section 1) to delineate the number of people downstream of each pixel and the downstream area of the critical natural assets.

### 3. Crop pollination

Up to two-thirds of all crops require some degree of animal pollination to reach their maximum yields (Klein et al. 2007), and natural habitat around farmlands can support healthy populations of wild pollinators by providing them with foraging and nesting resources. We model the potential contribution of wild pollinators to nutrition production based on pollination sufficiency of habitat surrounding farmland and the pollination dependency of crops. NCP for crop pollination is expressed in terms of the average equivalent number of people fed by pollination-dependent crops, attributed to nearby ecosystems based on the area of pollinator habitat within pollinator flight distance of crops.

**Pollination on-farm**

Pollination sufficiency is based on the area of pollinator habitat around farmland. Agricultural pixels defined by ESA 2015 (10 arc-sec, ~300 m resolution) with >30% natural habitat in a 2 km area surrounding the farm are designated as receiving sufficient pollination for pollinator-dependent yields (according to Kremen et al. 2004); pixel values between 0 and 30% of natural habitat are interpolated between 0 and 1, respectively (i.e., 0% has 0 value, 15% has 0.5, and 30% has 1.0). Crop pollination dependency of 115 crops is defined as the percent by which yields are reduced for each crop with inadequate pollination (ranging from 0-95%) according to Klein et al. (2007). Spatially explicit global crop yields (metric tons/ha) and the harvested area as a proportion of each grid cell at 5 arc-min (~10 km) are taken from Monfreda et al. (2008) for 115 crops to determine the total potential production. The crop production estimates are multiplied by crop pollination dependency to calculate the pollination-dependent crop production for each pixel, and multiplied by sufficiency to calculate pollinated production, the amount of pollination-dependent production that is actually produced. See Chaplin-Kramer et al. (2019) for more detail.

**Equivalent people fed**

Crop content of critical macro and micronutrients (KJ energy/100 g, IU Vitamin A / 100 g, and mcg Folate / 100 g) from the US Department of Agriculture is multiplied by pollinated crop production, then summed across all crops to derive pollinated nutrient production for each nutrient at 10 arc seconds. Each of the three micronutrients is divided by their Average Annual Requirement based on dietary guidelines, then averaged to calculate an “equivalent people fed” metric (see Chaplin-Kramer et al. 2019 for more detail). This is not the number of people whose nutrition is supplemented by pollination-derived production, but only a metric to summarize the number of people whose nutritional needs could be fully met by this amount of production.

Table SM3.1 Average annual dietary requirement per capita, used to calculate "equivalent people fed" metric (Source: Chaplin-Kramer et al. 2019).

| Nutrient | Average Annual Requirement |
|---|---|
| Energy (KJ) | 3319921 |
| Folate (mcg) | 132654 |
| Vitamin A (IU) | 486980 |

**Mapping back to habitat**

The result of the above analysis is a raster layer whose pixels represent the equivalent people fed metric per pixel. We mapped this value back to the pollinator habitat by constructing a Gaussian weighted distance decay with the same sigma as the one used to model flight decay distance in the original pollination model. In essence we "fly" the value score back to the habitat that hosted the pollinators that created the value in the first place with one caveat. We cannot directly redistribute the value score with the same Gaussian kernel since it would overcount the value without taking into account the total habitat area that contributed to the score in the first place. Directly weighting the value score back to the habitat without normalization is the equivalent of splitting a pie among N people by giving each person the entire pie. Hence, we use the following steps to ensure the value score projected back to habitat is correctly distributed over the total area of habitat:

1) We calculate the weighted distance decay of the habitat mask using the same Gaussian weighted distance decay used in the pollination model. This results in a raster whose pixel values represent the distance weighted number of habitat pixels that affect that pixel.
2) We divide the value score by this distance weighted decay habitat mask resulting in a raster whose pixel values represent the weighted distance value score over all contributing habitat to that pixel.
3) We calculate the final habitat value score by calculating the distance weighted decay of the previous raster using the same Gaussian kernel in previous steps over habitat pixels only. The final pixel values of this raster represent the effective value score that each habitat pixel contributes to the original value score.

### 4. Fodder production for livestock

Globally, livestock contribute to the livelihoods and food security of nearly one billion people (Robinson et al. 2014). Certain ecosystems, including grasslands and shrublands, provide critical pasture and fodder for livestock. We identify the places important for grazing and browsing livestock using Version 3 of the Co$ting Nature model (Mulligan 2018). NCP as fodder production for livestock is expressed in terms of an index (0-1) based on rangeland productivity and livestock density.

To calculate the relative realized grazing and browsing service, Co$ting Nature includes the global gridded spatial distribution and density of livestock mapped by the Gridded Livestock of the World (GLW) database (Robinson et al. 2014). This gridded global spatial dataset, derived from the United Nations Food and Agriculture Organization (FAO) data, shows where free-roaming livestock consume vegetation in pastures and other non-cropland areas. Supply is calculated as tonnes of dry matter productivity for the non-cropland cover fraction (i.e., including pastures, forests and other non-cropland uses) using a 10-year average of dry matter productivity data from Copernicus Service Information (Buchhorn et al. 2019) at 1 km resolution, 2009-2018; see Section 18. Dry matter productivity represents the overall growth rate or dry biomass increase of the vegetation and is directly related to ecosystem net primary productivity, however, with units customized for agro-statistical purposes (kg/ha/day). The head count of livestock in a grid cell is then multiplied by the average amount of biomass one animal would need to eat (estimated to be equal to 12kg/day (EPA, 1990). This *per-capita* consumption is then compared against dry matter productivity, and the realized service is reported as the smaller of the two (if consumption exceeds productivity the gap is assumed to be met with feed). The output, in terms of kg/ha of biomass grazed/browsed, is rescaled to unitless relative terms, meaning the map scale ranges from 0 to 1, with 1 denoting areas of highest performance for that service globally.

As the original resolution was 5 arc-min (~10 km) and we resampled to 2 km for the optimization, and because the model assumes all dry matter productivity within a pixel can be utilized by livestock, we needed to remove vegetation types for which the majority of dry matter is inaccessible to livestock (i.e., forests). The Co$ting Nature output was, therefore, masked to ESA land cover classes 30 (mosaic cropland >50% / natural <50%), 40 (mosaic natural >50% / cropland < 50%), 100-153 (mosaic tree and herbaceous, shrublands, grassland, and sparse vegetation), and 180 (flooded shrub/herbaceous; see Extended Data Table 3).

### 5. Timber production (commercial and domestic)

Forests, both natural and managed, provide timber for construction as well as wood and paper products for domestic use and export. Globally, forests provide over USD 600 billion, or 1% of global GDP, through wood-based products (World Bank 2020). While essentially the same benefit, we modeled commercial (e.g., for trade/export) and domestic (e.g., for local use) timber separately, because they represent two different sets of beneficiaries. NCP for timber production is expressed as an index (0-1) based on forest productivity and accessibility for harvest.

We represent sustainable timber production with two spatially mutually exclusive layers, one for commercial timber and one for domestic. The two layers are calculated using Version 3 of the Co$ting Nature model (http://www.policysupport.org/costingnature). For the purposes of this work, a given grid cell on the planet can provide either domestic or commercial timber, but not both. The two layers each represent the “realized” yield of sustainable timber biomass, meaning the total potential sustainable supply available is scaled by demand, which in some places equals supply and in others is lower. The scale for each map ranges from 0 to 1, with 1 denoting areas of highest performance for that service globally. Because these two layers are mutually exclusive (pixels are either commercial or domestic, not both), keeping them separate ensures that the optimization will reach the target on each layer.

These two layers were generated using Co$ting Nature and updated data for dry matter productivity, above-ground carbon stock and fractional tree cover. First, the total potential sustainable supply of timber for any use, commercial or domestic, was estimated from the above-ground carbon stock map for the year 2000 based on data from Global Forest Watch (Bacchini et al, 2012). The proportion of carbon coming from trees in a pixel was calculated as the product of carbon stock and fractional tree cover (Buchhorn et al., 2019) for rural areas (Schneider et al., 2009) only (urban trees are considered not to be usable for timber). The sustainable harvest is considered to be the reciprocal of the number of years taken to develop the stock at the annual sequestration rate, according to dry matter productivity data based on a 10-year average dry matter productivity at 1 km (using data from the SPOT VGT and Proba-V satellites for the period 2009-2018 from Copernicus Service Information (2019) Dry Matter Productivity; see Section 18).

Both timber layers have an original resolution of 5 arc-min (~10 km), and we resampled to 2 km for the optimization. Because there may be non-forest pixels within the larger 5 arc-min grid cells, and timber is only attributable to forest, we masked these layers to ESA forest classes: 30-110, 150, 151, 160, and 170 (see Extended Data Table 3).

##### *(a) Commercial timber*

To generate a "realized" commercial timber map in terms of tonnes harvested, the model identifies the potential timber supply within six hours' travel time of a population center of >50K people (Nelson 2008) and on slope gradients <31.5 degrees (70%) (Lehner et al., 2008) considered to be workable for logging (Greulich et al., 1999). This accessibility requirement represents the availability of transport infrastructure for commercial timber harvesting. Timber mass defined as accessible is constrained by slope to reflect the higher cost of removal (and increased wastage) on steeper slopes using a linear decrease in timber availability (from the potential availability to zero) as slope increases from 0 to 90 degrees. All extractable commercial timber is assumed to be extracted (demand=supply). Any timber considered not sustainably harvestable (beyond the annual replacement rate) is assumed to remain unharvested. Some of this would be available for fuelwood (see next section). The realized timber consumption is rescaled to a relative map of 0-1.

##### *(b) Domestic timber*

Domestic timber is also modeled using Co$ting Nature v3 (http://www.policysupport.org/costingnature). This model is derived from the same total potential sustainable timber supply map as above and is also reported in "realized" terms. No domestic timber harvest is permitted in urban areas and in areas where commercial timber extraction is present. Urban areas are defined using Schneider and Potere (2009). Areas for commercial timber extraction were mapped as above (domestic and commercial are spatially mutually exclusive). Demand was calculated differently as well; rather than using travel time from population centers, in this case we assume harvest is done on foot by rural people. Thus, demand was modeled based on population within the same grid cell (in this case, within each 10 $km^2$ grid cell) (according to Landscan 2017: https://landscan.ornl.gov/) and accessibility (one tonne dry weight per person per year in flat areas, declining to zero at 90 degree slope). Per capita use of wood was estimated based on published estimates (e.g., Banks et al. 1996). Thus, the realized domestic timber service equals the domestic timber demand (where demand<supply) and the supply (if demand >supply), with unmet demand assumed to be supplied via markets. As with commercial timber, some timber biomass will be unharvested and available for collection as fuelwood (see next section).

### 6. Fuelwood production

Forests also provide vital sources of firewood and charcoal for cooking and heating, which an estimated 880 million people depend on worldwide (FAO and UNEP 2020). We represent realized fuelwood yield (calculated in tonnes, rescaled to 0-1) with another output from Version 3 of the Co$ting Nature model (http://www.policysupport.org/costingnature). As for timber, NCP for fuelwood production is represented as an index (0-1) based on forest productivity and accessibility for harvest, but in this case specifically by rural people.

This layer was generated using Co$ting Nature, but with updated data for aboveground carbon stock and dry matter productivity, as described in the methods for timber. The layer is calculated first in tonnes but converted to relative terms, meaning the map scale ranges from 0 to 1, with 1 denoting areas of highest performance globally, and the grid cell resolution is 5 arc-min (~10 km). As with the domestic and commercial realized timber layers above, the total potential sustainable supply of timber is also used as the starting point for the fuelwood yield layer. In this case, demand is calculated as a function of rural population (rather than total population) derived from Landscan 2017, with a per capita use of 3.65 tonnes per year (OECD, 2006). The model assumes a linear decrease in fuelwood accessibility with increase in topographic slope, from baseline demand on flat ground to zero demand at a slope of 90 degrees. Thus, the service is equal to the fuelwood demand (where supply ≥ demand) or is constrained to the supply where (supply < demand). Fuelwood can overlap spatially with domestic and commercial timber use, given that domestic and commercial timber harvest will not consume all sustainably available woody biomass in all places, due to the slope gradient limit and/or in places where demand is less than supply. Moreover, much fuelwood is gained from the waste of commercial and domestic timber consumption. Timber only consumes the main trunks, whereas fuelwood will consume branches and wastage. Much wood is initially used for timber then used for fuelwood (i.e., the same biomass has two uses).

This layer has an original resolution of 5 arc-min (~10 km) and we resampled to 2 km for the optimization. Because there may be herbaceous pixels within the larger 5 arc-min grid cells, and fuelwood is best attributable to woody vegetation, we masked this layer to ESA forest and shrubland classes (including sparse and flooded vegetation): 30-122, 150-152, 160, 170, and 180 (see Extended Data Table 3).

### 7. Flood regulation

Ecosystems regulate water flows by retaining water during wet periods and releasing it during dry periods. This regulation function can contribute to reducing the frequency and severity of flood events for people living downstream (e.g., Stürck et al. 2014). We modeled this contribution using the "realized influential green storage metric" from Version 2 of the WaterWorld model (http://www.policysupport.org/waterworld) (additional methods information available in Gunnell et al. 2019). To map nature's influence on flood risk reduction, we identify the upstream places where canopies, wetlands, and soils (green storage) retain and slowly release rainfall, to the benefit of downstream communities on floodplains. NCP for flood regulation is expressed as an index (0-1) based on "green" water storage multiplied by the number of people downstream.

Green storage in a pixel is reported as a proportion of the total local and downstream storage by vegetation. This indicates the significance of the storage in a given pixel for reducing flood risk experienced by downstream beneficiaries, relative to the storage in all other downstream pixels also providing flood risk reduction for those beneficiaries. The index is calculated for a pixel as the green storage in that pixel divided by downstream total green storage (the influence metric) multiplied by the downstream summed beneficiary metric (excluding beneficiaries in the pixel of the storage since its influence on flow is downstream). In pixels where there are no downstream beneficiaries, the storage unit is multiplied by zero. The beneficiary metric used here is all people downstream based on Landscan 2017 (see Extended Data Table 2).

This layer has an original resolution of 5 arc-min (~10 km), and we resampled to 2 km for the optimization. Because we are focused on the contribution of natural assets, we masked these layers to natural and semi-natural ESA (including natural/crop mosaic) classes: 30- 180 (see Extended Data Table 3).

### 8. Access to nature

Ecosystems provide numerous direct and indirect benefits to people, such as recreation, hunting and gathering, aesthetics, mental and physical health, cultural and traditional value, and sense of place. Some of these contributions depend on the ability of people to access nature. Therefore, we mapped nature that is accessible to people, as a proxy for nature's contributions to people such as gathering and recreational opportunities within nearby nature, following the logic of Schirpke et al. 2018 (but in reverse; instead of identifying places within access of residential areas, counting people within access of every "natural" pixel). This proxy-NCP is expressed as the number of people within 1 hour (or 6 hours, for sensitivity analysis) travel of natural and semi-natural lands.

The travel time is calculated based on a friction layer developed by Weiss et al. (2018) that represents the minutes per meter to travel across a pixel, based on information about roads, railroads, rivers, bodies of water, topographical conditions (elevation and slope angle), land cover, and national borders, combined so that the fastest mode of transport takes precedence. Therefore, the distance people may be from natural habitat may vary widely based on the available infrastructure and terrain. We include both 1 hour and 6 hours travel to test the sensitivity to assumptions of how long people are willing to travel to access nature; 1 hour may represent more casual recreation or daily gathering activities, while 6 hours may represent the time taken for a longer or more serious trip. This assumption does not significantly impact the total area selected in the optimization, but it doubles the area required when this NCP is optimized individually (see Section 22; Extended Data Table 5, row 1 vs. row 6, and row 27).

As for other NCP in this analysis, natural and semi-natural lands are defined as ESA Land Cover classes 30-180 (see Extended Data Table 3; 10 arc-sec, ~300 m resolution), which includes natural/crop mosaics. Population is taken from Landscan 2017, split into urban and rural population based on urban-rural catchment areas (URCA) delineated by Cattaneo et al. 2021. The URCA hierarchy classes 1 (large city, > 5 million people) to 7 (small cities and towns, 20,000-50,000 people) are defined as “urban” and classes 8 and higher (<1 hour to a large city up to hinterland) are defined as “rural”. The number of rural and urban people within the given travel time of each pixel of natural and semi-natural habitat are included as separate objectives in the optimization but optimized together (similar to commercial and domestic timber, Section 5) so that target levels of each are reached.

*Computational Method*

We calculate the number of people that can reach a pixel in a given amount of time by solving a modified Djikstra's least cost path algorithm (Dijkstra 1959) for every non-zero population pixel in the raster. We count the travel time between pixels as the average friction value multiplied by the distance between pixels and we terminate the algorithm when we can no longer travel to a pixel without exceeding the allotted total travel time. The result of this algorithm for a single pixel is a "reachable" mask that shows what pixels can be reached from the originating pixel. We add the population value of the originating pixel to a global raster that intersects this "reachable" mask. By running this algorithm for every pixel in the raster we end up with a global value of the number of people that can reach any pixel.

We also generate masks of "reachable areas" where we start with a 0/1 mask indicating some pixels of interest such as habitat and a global travel time and generate a 0/1 mask raster that is 1 where a pixel can reach any pixel of interest in the base raster. This requires a slight modification to the algorithm above that rather than accumulating a running total of population, we only record pixels which are covered in the given travel time as a single Boolean value; overlapping coverage doesn't otherwise change this value.

### 9. Riverine fish catch

Wild fish from rivers and lakes support the food security of 158 million people globally (McIntyre et al. 2016). We mapped global inland fisheries by updating the approach from McIntyre et al. (2016) to include recent progress in addressing catch underreporting and mapping harvests. Updates in the new maps fall into three categories: i) updates to catch estimates, using a longer time window, ii) updates to the mapping algorithm to account for wetlands, iii) finer spatial resolution. These maps can be used to assess the contribution of capture inland fisheries alongside rapidly improving representation of aquaculture and other food-production systems. NCP for riverine fish harvest is represented as metric tonnes of fish caught per $km^2$ per year, spatially disaggregated to the locations of the catch.

The map was generated from 40 training basins with catch estimates compiled from literature. We then fitted a multiple linear regression model of total catch predicted by population (in persons; CIESIN 2017), discharge (cubic liter per sec from HydroSHEDS; Lehner & Grill 2013), and percent wetland cover (Mean Annual Maximum inundated extent from Fluet-Chouinard et al. 2015). We predicted catch with this regression model across the global land surface. The absolute values of these predictions tend to overestimate actual, but this provides a surface over which catch estimates can be mapped. Finally, we distributed national catch within each countries' borders proportionally to the pixels. The national catch is a merger of FAO catch statistics for inland fisheries between 1997-2014, to which catch was revised using household surveys over 42 countries with known deficiencies in official statistics (Fluet-Chouinard et al. 2018).

For the analysis in this paper, we chose to truncate the fishery data which originally spans 15 orders of magnitude. We truncated catch estimates below 1 ton per 10x10 km cell, and then truncated the 99th percentile on the high end. As for all layers, we masked to ESA natural land cover classes, but included water bodies as an additional class for this layer (see Extended Data Table 3).

### 10. Marine fish catch

Marine fisheries also provide critical sources of food and income, supporting livelihoods for more than 260 million people and food security for over 524 million (Selig et al. 2019). We mapped marine fish catch using an update from Watson and Tidd (2018), which attributes tonnes (metric) of fish catch per sq km within 30 min grid cells across the ocean. As for riverine fish harvest, NCP for marine fish harvest is represented as metric tonnes of fish caught per $km^2$ per year, spatially disaggregated to the locations of the catch.

This catch data is based on data published by the Sea Around Us (Pauly et al. 2020). This mapping was further improved (Roberson et al. 2020) by using fine scale regional reporting such as the tuna regional management organization's (tuna rFMO) and observed patterns from satellite data from Global Fishing Watch's (GFW's) vessel Automatic Identification System (AIS)-based data (de Souza et al. 2016).

For this analysis, we used average annual total marine fish catch per grid cell from 2010-2014. The raw data is in units of tonnes per 30 min grid cell, including estimates of unreported and illegal catch by commercial and non-commercial sectors. However, because grid cells are of unequal size depending on where they occur on a spherical Earth, the catch was adjusted by the grid cell sizes (measured in sq km), so final units are "tonnes per sq km" within 30 min grid cells. Note that high values were not "clamped" as marine catches tend to be over-dispersed spatially compared to logbook data which has information on fishing hot spots. Thus, the annual catch rate per square km can be very high, as there are some very well-defined fishing spots, and high values can actually be an underestimate of fish catch in a given area.

Jurisdictional EEZ boundaries are used in the mapping to decide on access by national fleets. So if a country is not known to leave its own waters (EEZ claimed area) and if there are no other countries fishing in the area, mapped catches stop at this boundary (dictated also by other factors relevant to the species involved). But often the fleets that fish in international waters are reported as fishing nearby. If there is no evidence that they have permission to fish in the EEZ of a country then (depending on the species fished) their efforts are concentrated on the boundary. In reality they may fish deep into the EEZ area if there is no enforcement. Even sightings via satellite may not deter them if there are no vessels to apprehend them on the water. Meanwhile, many species have higher biomasses in shallower/inshore productive waters causing a strong incentive to fish nearer to shore. Combining these effects results in a phenomenon of foreign fishing creating 'haloes' around EEZ areas.

### 11. Coral reef tourism

Nature-based tourism provides important livelihoods for many people throughout the world. Here we focus specifically on tourism associated with coral reef habitat, using data from Spalding et al. 2017, who documented that coral reefs provide a total estimated value to the tourism sector of USD $36 billion worldwide. NCP for coral reef-associated tourism is expressed as dollars of tourism expenditure (in deciles 1-10), which is a way of assessing income or livelihoods derived from nature.

The value of coral reefs for travel and tourism was estimated as a proportion of national expenditure statistics for all coral reef jurisdictions. Figures include direct reef-use or "on-reef" tourism as well as expenditures associated with indirect or "reef-adjacent" benefits such as water quality, coral sand beaches and seafood. National expenditure data, taken from the World Travel and Tourism Council data (averaged from 2008-12 values), were distributed based on two independent sources of tourism distribution: hotel rooms from the commercial Global Accommodation Reference Database (GARD), and user-generated photos from the image-sharing website, Flickr. Once distributed expenditure values were extracted for reef-adjacent coastlines defined as coastal, non-urban areas, < 30 km from coral reefs). These expenditure numbers provided a starting point for assessing reef tourism values.

Reef-adjacent values were simply assigned as a simple estimate of 10% of the value of all reef-coast tourism to coral reefs, based on earlier literature reviews. On-reef tourism values were developed in a 2-stage process, first assigning national level importance to on-reef activities and then spreading the resulting value to reefs. The national importance statistics were informed by a combination of existing literature and by two statistics generated for each country: the abundance of dive shops relative to hotel rooms, and the abundance of underwater photographs relative to all photographs shared on Flickr. Based on these numbers, the literature and minor expert-informed interventions, a single figure for on-reef tourism value was assigned as a proportion of the remaining 90% of reef-coast tourism values (excluding the 10% assigned to reef-adjacent values) for each country.

Finally, both on-reef and reef-adjacent values were attributed to reefs on a 15 arc-sec (~500 m) gridded global reef map (Burke et al. 2011). Reef-adjacent tourism was spread with weighting driven by both the density of tourism (determined by hotel rooms and Flickr imagery) and by the distance of reefs (up to 30km) from this tourism density. On-reef tourism was spread to reefs using two georeferenced data sources representing on-reef activities: the location and density of underwater Flickr imagery and the location and density of dive-sites. The resulting maps thus linked expenditures to both on-reef and reef-adjacent use, which were combined to give a layer of total expenditure linked to these reefs.

### 12. Coastal risk reduction

Coastal habitats such as coral reefs, mangroves, salt marsh, or sea grass, attenuate waves and protect the shorelines from the impacts of storms, such as floods and erosion. These coastal protection benefits protect more than 76 million people globally (Selig et al. 2019). How much this attenuation matters depends on the physical exposure to coastal hazards (based on wind, waves, sea level rise, geomorphology, etc.) and the people exposed. To map coastal risk reduction to coastal habitats, we used methods based on the InVEST Coastal Vulnerability (http://releases.naturalcapitalproject.org/invest-userguide/latest/coastal_vulnerability.html) model as described in Chaplin-Kramer et al. 2019, with two major updates: 1) inclusion of new data, and 2) projecting the value back to the habitat. NCP for coastal risk reduction is expressed as a unitless index of the coastal risk reduced by habitat multiplied by the number of people within the protective distance of the habitat.

**Inclusion of new data**

In addition to the sources described in Chaplin-Kramer et al. 2019, we added terrestrial coastal habitat and included updates to offshore coastal habitat, and we included a global geomorphology layer in our calculation of coastal risk.

Terrestrial habitats were taken from ESA LULC, clipped to within 2 km of the shoreline (the maximum protective distance of any habitat). Although much of this terrestrial habitat is located behind the coastline and the storm surge that would hit it, including this habitat is important to represent mitigation of potential flooding. We also updated coastal habitat layers with more recent datasets for coral reefs, mangroves, and sea grass (the salt marsh habitat layer remains the same as in Chaplin-Kramer et al. 2019). We masked out mangroves and salt marsh that spatially overlapped with terrestrial coastal habitat so they were not double-counted. The risk ranks and protective distances of each habitat type are shown in Table SM12.1.  The data sources for each habitat are listed in Section 18 (Coastal habitats).

**Table SM12.1** Parameterizations for the InVEST Coastal Vulnerability model: risk rank (with 1 being the lowest) and maximum protective distances of the different habitat types.

| Habitat | Risk rank | Protective distance |
|---|---|---|
| Coastal forests | 1 | 2000 |
| Wetlands and swamps | 2 | 1000 |
| Coastal scrub | 2 | 2000 |
| Mosaic natural / cropland | 4 | 500 |
| Sparse vegetation | 4 | 500 |
| Coral Reefs | 1 | 2000 |
| Mangroves | 1 | 2000 |
| Sea grass | 4 | 500 |
| Salt marsh | 2 | 1000 |

Geomorphology or shoreline (substrate) type is another important variable used in determining coastal risk, as hard/rocky shorelines are less vulnerable than soft substrates. Acquisition of an unpublished global dataset for geomorphology (Vafeidis et al. 2008) allowed us to include this variable in this analysis, where it had been omitted in Chaplin-Kramer et al. 2019. We assigned all substrates a risk rank of 5 (the highest) except for the class defined as “unerodible,” which we assigned a rank of 1. Wherever the dataset was nodata, we calculated the geometric mean of the total coastal risk with one fewer variable. The original geomorphology dataset available at: https://storage.googleapis.com/critical-natural-capital-ecoshards/SedType_md5_d670148183cc7f817e26570e77ca3449.zip The processed dataset used in this analysis is at: https://storage.googleapis.com/critical-natural-capital-ecoshards/geomorphology_md5_e65eff55840e7a80cfcb11fdad2d02d7.gpkg

**Projecting coastal risk reduction value back onto habitat**

We projected the NCP onto the habitat that supplied it by mapping both the service values (or “nature’s contribution”) and the beneficiary values (delineating “to people”) for specific habitats. In order to calculate service values of a given habitat we subtracted the current risk (with all habitat) from the risk without that specific habitat (e.g., a coral reef, mangrove, or coastal wetland). This is necessarily different from Chaplin-Kramer et al. 2019 because it focuses on the difference from risk without a specific habitat instead of the risk without all habitat to attribute the marginal service value of that specific habitat. Each shore point is then buffered by the protective distance of the given habitat type, and the intersection of the buffered shore points with the habitat layer are rasterized. All overlapping buffered points are then added to the service value. Beneficiaries are defined as the population that benefits from a particular point on the shoreline is the count of people in pixels that are <10 m in height and within the protective distance of each habitat. These beneficiaries are attributed to habitat in a similar method as projecting NCP onto habitat except the population values are first divided by the area intersected with the habitat layer from the same circle used to buffer the protective distance in the NCP calculation. Normalizing by this area ensures that the sum of the distributed population on habitat equals the sum of the base population. Terrestrial vs. offshore coastal habitats were optimized separately, so targets could be reached in both realms.

### Global climate NCP

#### *13. Vulnerable terrestrial ecosystem carbon*

Ecosystems store large amounts of carbon in their aboveground biomass (e.g., vegetation), belowground biomass (e.g., roots), and soils. If properly managed, ecosystem carbon stocks can help mitigate the effects of global climate change by avoiding emissions of carbon dioxide to the atmosphere. Globally, ecosystems such as peatlands, mangroves, and old-growth forests and marshes contain at least 260 Gt of carbon (Goldstein et al. 2020). This analysis is based on vulnerable terrestrial and mangrove ecosystem carbon storage (aboveground, belowground, and soils), based on what carbon is likely to be released if the ecosystem were converted, by combining a variety of datasets, mapped by Noon et al. 2021, and building on the conceptual framework in Goldstein et al. (2020).

Noon et al. first created a “total terrestrial ecosystem carbon storage” map using the best available datasets (Table SM13.1) to serve as a baseline of total carbon, including both biomass and soil carbon. This dataset accounts for soil depths most likely to be affected by disturbances (100 cm depth for mangrove soil carbon and 30 cm depth everywhere else on land). The amount of total carbon was converted into the magnitude of vulnerable carbon, based on its initial stock, the conversion driver and the vulnerability of its carbon pools. Average carbon losses from biomass and soils due to the most common anthropogenic disturbances were quantified, including: conversion to agriculture in grasslands, wetlands and tropical forests; forestry in boreal and temperate forests; and aquaculture or built infrastructure in coastal ecosystems. The feasible loss events were only considered in terms of those that would alter the land cover (for example, forest to soy field) as opposed to activities that might reduce the carbon content but not constitute full conversion (for example, forest degradation due to charcoal collection or selective logging). Vulnerable carbon is therefore the portion that would be lost in a hypothetical but typical conversion event; it does not characterize the likelihood of that conversion event. The units of measurement for the resulting map are tonnes of carbon per ha, and the original resolution is 30 m.

**Table SM13.1** Data included in the map of terrestrial ecosystem carbon storage

| **Biomass** | **Resolution** | **Citation** |
|---|---|---|
| Above-ground biomass | 300m | Spawn et al. 2020 |

| | | |
|---|---|---|
| Below-ground biomass | 300m | Spawn et al. 2020 |
| Mangrove AGB | 30m | Simard, 2019 |
| Mangrove BGB | 30m | Hutchison et. al. 2013 |
| Mangrove extent | 0.8 arc seconds | Bunting et al., 2010; GMW, 2016 |
| Seagrass AGB | 30m | Fourqurean et al. 2012 |
| Seagrass BGB | 30m | Fourqurean et al. 2012 |
| Seagrass extent | 1:1,000,000 | UNEP-WCMC, 2018 |
| Salt marsh AGB | 30m | Byrd et al. 2018 |
| Salt marsh BGB | 30m | Byrd et al. 2018 |
| Salt marsh extent | Between 1:10,000 to 1:4,000 | McOwen et al. 2017 |
| **Soils** | **Resolution** | **Citation** |
| Soil Organic Carbon (SOC) | 250m | ISRIC, 2019; Hengl et al., 2017 |
| Mangrove SOC | 30m | Sanderman, 2018 |

The error associated with this dataset was derived by mosaicking the pixel-level uncertainty layers that accompanied each of the initial global input datasets (aboveground carbon, belowground carbon, soil organic carbon), and combined with uncertainty ranges around each of the emission and sequestration factors used, resulting in a spatial 'uncertainty map'.  The total uncertainty in vulnerable ecosystem carbon was found to be 332.54 +/- 152.98 for biomass carbon, 131.56 +/- 49.02 for soil carbon, and 464.07 +/- 176.98 for total carbon.

### 14. *Atmospheric moisture recycling*

Atmospheric moisture recycling is the process of water arising from the surface of the earth as evaporation, flowing through the atmosphere as water vapor, and returning to the surface of the earth as precipitation. In this case, evaporation includes canopy interception, soil interception, soil evaporation, vegetation transpiration, and open water evaporation. Likewise, precipitation includes all forms of liquid and solid precipitation. To quantify the moisture recycling benefits associated with intact vegetation globally, we calculated the volume of water evapo-transpired that falls on all rainfed productive land.

The rainfed productive lands are identified using the Anthromes dataset (Ellis and Ramankutty 2008). We selected all populated rainfed lands, including settlements, croplands, rangelands, and forested lands (irrigated lands were excluded since they are not reliant on rainfall, and wildlands, because they are not occupied social-ecological systems). Remote subcategories were included because, though population density may be low, those places may still be important for food production.

Nature's contribution to these productive lands is determined by where the moisture originates. In order to quantify the source of moisture, where it travels through the atmosphere, and where it falls out downwind, we employed an Eulerian moisture tracking model named the Water Accounting Model, 2 layers (hereafter referred to as the WAM-2layers) (Rudi J. van der Ent et al. 2010; R. J. van der Ent et al. 2014). Data for this analysis are from the European Centre for Mesoscale Weather Forecasting (ECMWF) climate reanalysis product, ERA-Interim (Dee et al. 2011). Reanalysis data are considered quasi-observational, since they combine global surface, air, and space-borne measurements into a global climate model. In this study, data were downloaded at a 1.5-degree spatial resolution. Six data types were used, including three-dimensional (i.e., full atmospheric column) zonal winds, meridional winds, and specific humidity; and, two-dimensional (i.e., surface) evaporation, precipitation, and surface pressure. The temporal resolution was sub-daily; surface pressure, winds, and specific humidity were downloaded at the 6-hourly time step; evaporation and precipitation were downloaded at the 3-hourly timestep. The analysis was run from the year 2000 to 2014.

The WAM-2layers tracks the daily flux of atmospheric moisture into and out of a hypothetical column of atmosphere (using an upper and lower atmospheric layer, to account for wind shear). Then, by identifying the precipitation that falls on the specific region (either the realized or critical regions), we can track the moisture backwards in time along the atmospheric trajectory that the water vapor flows, eventually to the evaporative origin of the moisture upwind. In this way, we generate moisture source regions for both the realized and critical regions. Some of these regions are over oceans, but much of the moisture originates on land.
We also want to identify the importance of vegetation specifically to the provision of moisture. So, using existing analysis that quantified the fraction of evaporation that is regulated by vegetation on land (relative to a hypothetical bare ground state) we quantify the fraction of this source moisture that comes from upwind vegetation (Wang-Erlandsson et al. 2014; Keys et al. 2016).

### Diversity

#### 15. Biodiversity: Species Area of Habitat

As a measure of biodiversity, we evaluated the overlap between areas optimized for NCP and species "Area of Habitat" (AOH) for 26,709 terrestrial vertebrate species. As described in Brooks et al. 2019, AOH areas are based on species range maps from IUCN, but refined using habitat preferences and elevational limits from IUCN Red List data. AOH are more specific than extent-of-occurrence (EOO) which can overestimate species range sizes (Hurlburt and Jetz 2007). AOH areas "exclude areas of unsuitable habitat from each species' range, which reduces commission errors and more closely approximates the actual occurrence of the species" (Soto-Navarro et al. 2020).

Species AOH ranges were produced for 10,774 species of birds, 5,219 mammals, 4,462 reptiles and 6,254 amphibians with available IUCN range polygon data following the procedure outlined in Brooks *et al*. 2019. Species range polygons obtained from the IUCN Red List spatial data portal (IUCN Red List and UN Environment Programme World Conservation Monitoring Centre 2020) and the Birdlife International spatial data zone (BirdLife International and Handbook of the Birds of the World 2019) were first filtered for 'extant' range then rasterized to a global 1 km grid in the Eckert IV equal area projection. Individual species range rasters were then modified to only include land cover classes that match the habitat associations for each species. Habitat associations were obtained from the IUCN Red List species habitat classification scheme and were matched to ESA land cover classes for the year 2018 following the crosswalk table presented in Santini *et al*. 2019. ESA land cover classification data was aggregated from its native 300 m resolution to match the global 1km grid using a majority rule. Species ranges were additionally filtered so that only areas within a species accepted elevational range were included. Global elevation data derived from SRTM was obtained from WorldClim v. 2 (Fick *et al*. 2017). For bird species, seasonal range codes 1-3 (1=year-round; 2=breeding range; 3=non-breeding range) were processed individually and stored as separate range files where applicable.

#### 16. Linguistic diversity

We analyzed the contributions of nature to cultural diversity through an analysis of languages associated with model solutions for various optimization scenarios. Languages serve as a useful proxy for cultures, being the primary mechanism for codifying the knowledge characterizing a given cultural system and as the means by which the systematic behavior patterns that compose cultures are conveyed from one generation to another. As an indicator of NCP to cultural diversity, the analysis presents the number of languages that co-occur with optimization model results.

The language analysis employed the most complete dataset available on global languages, a catalogue called *Ethnologue* compiled and frequently updated by SIL International (SIL n.d.). This constitutes the best and most comprehensive data on global languages available. The analysis used a geographic information system version of the language data created by SIL based on the 23rd edition of *Ethnologue* (Eberhard et al. 2020). This study only considers *Indigenous and non-migrant* languages—those traditionally linked with a particular sociocultural group and place—rather than languages whose locations and speakers have changed with migration, colonial expansion, and like processes (such as Portuguese in Angola, Brazil, Cape Verde, Guinea-Bissau, and Mozambique). The analysis focused on languages whose ranges appear as polygons, providing much more precise information on their co-occurrence with natural assets than point data. Language data remained in their original vector format, to enable identification of both number of languages co-occurring with an optimal natural asset solution and their individual identities; to enable the analysis, we converted natural asset data from their original raster format to vector format. Any instance where a language intersected an occurrence of natural assets, no matter the amount of overlap, was taken to signify a co-occurrence.

### Additional datasets

#### 17. Land cover

For all terrestrial datasets, we masked the original data using land cover classes from ESA 2015 land cover data (ESA Climate Change Initiative 2017, http://maps.elie.ucl.ac.be/CCI/viewer/download.php), excluding crop and urban areas primarily because accurately representing the contribution of these types of habitat to providing benefits to people depends on their management, and detailed spatial information on crop and urban nature management is not currently available globally. We also exclude bare areas and permanent snow and ice because the current models are not parameterized well to capture any role these ecosystems may be playing. For specific NCP that are modeled at a coarser resolution, we include additional masking to ensure that the value is attributed to the land cover classes that are thought to be providing that value (see Extended Data Table 3 for more detail).

#### 18. Coastal habitats

We rasterized all polygon data mapping offshore coastal habitat (coral reefs, mangroves, sea grass, salt marsh) and extracted relevant habitat types from ESA land cover (Section 17) for terrestrial coastal habitat. Coral reef habitat was used for modeling coastal risk reduction and coral reef tourism; mangrove habitat was used for modeling coastal risk reduction and carbon. All other habitats were used for the coastal risk reduction model; terrestrial habitats were clipped to within the protective distance of each habitat type for coastal risk reduction (Section 12). See Table SM18.1 for a list of habitat type data sources and availability.

**Table SM18.1** Coastal habitats used in this analysis.

| Habitat type | Original source | Data layer used in analysis |
|---|---|---|
| Coastal forests | ESA LULC classes | https://storage.googleapis.com/critical-natural-capital- |

| | | |
|---|---|---|
| | 50-100 | ecoshards/1_2000_mask_md5_91e7f997e1197e4a2abf064095e2179e.tif |
| Wetlands and swamps | ESA LULC classes 160-180 | https://storage.googleapis.com/critical-natural-capital-ecoshards/2_1000_mask_md5_f428433ea05cf1c7960ec7ff3995c5aa.tif |
| Coastal scrub | ESA LULC classes 110-140 | https://storage.googleapis.com/critical-natural-capital-ecoshards/2_2000_mask_md5_1ffc23cd09f748e1fe5a996b72df3757.tif |
| Crop/natural mosiac; sparse vegetation | ESA LULC classes 40, 150-153 | https://storage.googleapis.com/critical-natural-capital-ecoshards/4_500_mask_md5_6f48797ca1ab8e953e32288efd0536a4.tif |
| Coral Reefs | Reefs at Risk (Burke et al. 2011) | https://storage.googleapis.com/critical-natural-capital-ecoshards/ipbes-cv_reef_md5_5a90d55a505813b5aa9662faee351bf8.tif |
| Mangroves | Global Mangrove Watch (Bunting et al. 2018) | https://storage.googleapis.com/critical-natural-capital-ecoshards/ipbes-cv_mangrove_md5_2205f546ab3eb92f9901b3e57258b998.tif |
| Sea grass | UNEP-WCMC and Short (2018) | https://storage.googleapis.com/critical-natural-capital-ecoshards/ipbes-cv_seagrass_md5_a9cc6d922d2e74a14f74b4107c94a0d6.tif |
| Salt marsh | (Unchanged from Chaplin-Kramer et al) Mcowen et al. 2017 | https://storage.googleapis.com/critical-natural-capital-ecoshards/ipbes-cv_saltmarsh_md5_203d8600fd4b6df91f53f66f2a011bcd.tif |

### *19. Dry matter productivity*

We calculated a 10-year average of dry matter productivity (1 km resolution) using data from Copernicus Service Information (2019) Dry Matter Productivity for the years 2009-2018. Dry Matter Productivity (DMP) is an estimate of the total growth (in terms of increase in biomass) of vegetation. The units of measure are kg/ha/day, but for our analysis we calculated a 10-year average. The averaged 10-year DMP data and associated R code available at: https://drive.google.com/drive/folders/107DKplh5_D1_aFSmnv_ekHjqTmAIrXLy

### *20. Human population*

We used human population data from Landscan 2017 (Rose et al. 2018; data available from: https://landscan.ornl.gov/. LandScan 2017 population data is modeled using sub-national level census counts for each country, land cover, roads, slope, urban areas, village locations, and other high-resolution imagery. The population model predicts a “likelihood” coefficient for each grid cell and allocates total population from an area to each grid cell based on the coefficient. The results are reported in terms of “ambient or average day/night population count.” For more details, see https://landscan.ornl.gov/. The resolution of this dataset is 30 arc-sec (~1 km).

### 21. Land and ocean boundaries

We used country boundaries combined with their associated Exclusive Economic Zones (EEZ) from MarineRegions.org Version 3, released 2020-03-17, of the "marine and land zones" product (https://www.marineregions.org/sources.php#unioneezcountry). We combine distinct territories of sovereign nations together into a single feature and attribute any disputed territory to all its claimants. This treatment of disputed territories leads to these regions being counted multiple times, once for each claimant.

### Analysis

### 22. Optimization using prioritizr

To identify areas providing the highest value across different NCP, we conducted an optimization using a “minimum set” objective function (that is, identifying the grid cells which collectively provide the highest NCP values in the least amount of area.) We used integer linear programming (ILP), following methods similar to Schuster et al. 2019:

> “The general form of an ILP problem can be expressed in matrix notation as:
>
> $$\text{Minimize } cx \text{ subject to } Ax \geq b$$

where $x$ is a vector of decision variables (in our case, whether to prioritize an individual planning unit), $c$ and $b$ are vectors of known coefficients, and $A$ is the constraint matrix. In the minimum set cover problem, $c$ is a vector of costs for each planning unit, $b$, a vector of targets for each conservation feature, the relational operator would be ≥ for all features, and $A$ is the representation matrix with $A_{ij} = r_{ij}$, the representation level of feature $i$ in planning unit $j$. We set an objective to find the solution that fulfills all the targets and constraints for the smallest area, which we use as our measure of cost. This objective is similar to that used in Marxan, the most widely used spatial conservation planning tool, but has been shown to lead to more efficient solutions.”

##### *Code repository*

Analyses were conducted using prioritizr (https://prioritizr.net/) (Hanson et al. 2020a). For additional details on prioritizr and integer linear programming, please see Schuster et al. 2020 (Supplementary materials.) We have developed a repository with all the code used in the prioritizr analysis on the Open Science Framework (OSF): https://osf.io/r5xz7/?view_only=d611a688525f4ceb8db4ef4e7528b0e8

#### Targets

To explore the land/ocean area required to maintain different levels of NCP provision, we repeated the optimization using 20 different targets ranging from 5% to 100% of total NCP value, across all NCP, at 5% increments. In other words, we identified areas that collectively provided at least 5% of the total value of all NCP, 10% of all NCP, 15%, and so forth up to 100%. For example, the 90% target identifies areas that provide at least 90% each of total nitrogen retention, sediment retention, crop pollination, fodder, and so on.

#### Global and country-by-country optimization

We conducted the optimization at two spatial scales: globally and country-by-country. The global optimization identifies where the highest NCP values are worldwide, and therefore might be useful for international policy or decision-making. The country-by-country optimization identifies areas within each nation's land and marine jurisdiction (EEZ) (that is, areas providing the top 5%, 10%, etc. of all NCP within a given country), and therefore might be useful for government policy or decision making. For the country-by-country optimization, we ran the optimization within each country and then combined the outputs to produce a global map.

As described above, we used combined land and EEZ boundaries from MarineRegions.org, combining distinct territories of sovereign nations together into a single feature and attributing any disputed territory to all its claimants. Raster layers are converted to equal area grids (Eckert IV projection) at 2 km resolution and masked to the combined land + EEZ layer. In addition to these global layers, a country-specific layer for each ecosystem service is produced. Each ecosystem service within each country (and globally) is treated as a feature in the prioritization. For computational efficiency, all layers are converted to an aspatial format for prioritization; the prioritization solutions can then be converted back to spatial format.

#### Optimization objectives

We ran the following types of optimizations:

1. all "local" NCP (12 listed in Fig 1A) together, by country
2. global NCP (carbon and moisture) together, globally
3. scale sensitivity: all "local" NCP together, globally
4. scale sensitivity: all NCP (the 12 "local" and 2 global) together, globally
5. nature access sensitivity: all "local" NCP (substitute nature access within 1 hour travel for nature access within 6 hour travel; see Section 8), by country
6. hydrologic NCP sensitivity: all "local" NCP (substitute 500km flow attenuation for hydrologic NCP for 50km flow attenuation; see Section 1), by country
7. NCP set sensitivity: drop 1 NCP (drop each one of the 12 "local" NCP set), by country
8. NCP correspondence: each of the 12 "local" NCP by itself (including sensitivity NCP above, but not carbon or moisture), by country
9. NCP correspondence: each of the global NCP (carbon and moisture), by themselves, globally

The result from Optimization 1 is featured in Fig. 1 and compared to the result from Optimization 2 in Fig. 3 in the main text (Extended Data Table 5, rows 1 and 2). Optimizations 3 and 4 test the sensitivity of the solution to scale (Extended Data Fig. 4), along with Optimizations 5 and 6 for assumptions of scale of influence for specific NCP (Extended Data

Table 5, rows 4-7). The assumptions of scale for nature access (Optimization 5) and hydrologic NCP (Optimization 6), make very little difference to the overall area selected (Extended Data Table 5, rows 6 and 7, respectively), but optimizing the "local" NCP at a global scale requires vastly less land and EEZ area to reach 90% of all NCP (22.5% vs. 30.4% of land and 12.9% vs. 24.2% of EEZ; Extended Data Table 5, rows 4 and 1, respectively). However, this necessarily means that some countries do not retain close to their current NCP values, with the majority of area being consolidated in Asia and Africa (Extended Data Fig. 5). The difference in area required for reaching 90% of NCP through adding the solutions for Optimizations 1 and 2 together (44%, shown in Fig. 3 in main text, and row 3 of Extended Data Table 5) vs. through Optimization 4 which prioritizes all 14 NCP together globally (39.5%, row 5 of Extended Data Table 5) is smaller, meaning the efficiency gains are less when switching from local to global optimization for local NCP if global NCP are also included. However, the difference between Optimization 2 for global NCP only (38.5%, row 2 of Extended Data Table 5) and Optimization 4 for global and local NCP optimized globally together (39.5%, as above) is negligible; adding these additional NCP to a global optimization requires almost no additional area.

The sensitivity to the set of NCP included (Optimization 7) was tested in 12 different optimizations (listed in Extended Data Table 5, rows 8 through 19), dropping each individual NCP in turn from the set and solving for the 90% target of the remaining 11. The resulting binary (1/0) solution rasters were averaged together to calculate the area shared by all rasters (in 100% of the solutions) and the area shared by 11 of the 12 solutions (shown globally and broken down by country in SI Table 1). The correspondence between different pairs of NCP (shown in Extended Data Table 4) was calculated by taking the 90% target solutions for the 14 optimizations for individual NCP prioritized at their relevant scale, by country for local NCP (Optimization 8) and globally for global climate NCP (Optimization 9), multiplying the binary (1/0) solutions rasters of each pair, and calculating the area of overlap. Total areas for each solution were also calculated (shown in Extended Data Table 5, rows 20-32).

Our focus here is on NCP; ongoing analyses include optimizations for both NCP and biodiversity (>29,000 terrestrial vertebrate species). It was not possible to combine country-level prioritized local NCP with globally prioritized global NCP (carbon storage and atmospheric moisture regulation) because it would require each country to be treated as a separate feature, meaning 198 countries x 14 NCP maps (12 NCP, but timber split into domestic and commercial and coastal risk reduction split into coastal habitats and barrier reef) + 2 global NCP maps = 2774 features, which is not computationally feasible at a resolution of 2 km on conventional computers.

Additional global priorities (i.e., vertebrate biodiversity, cultural diversity; Fig. S1b) are not included in the optimization results reported here. Both were included in a separate analysis (see "Overlap analysis," below). For cultural diversity, we use the count of languages as a proxy (see "Languages," above). We originally included a combined linguistic diversity layer (0.5 degree) in preliminary optimization runs, but it greatly affected the results due to its relatively coarse spatial resolution (a significantly larger area was selected in the optimization, due to the large size of each grid cell relative to the other NCP maps). For vertebrate biodiversity, due to the large number of features (>26,000) and long processing times required, we were unable to include these in the optimization in time for this analysis. We are also in the process of conducting a follow-up optimization that includes both NCP and vertebrate biodiversity.

### 23. Overlap analysis

We analyzed the level of overlap between country-level NCP areas and global-scale priorities, including carbon, atmospheric moisture recycling, biodiversity, and cultural diversity, in several different ways. All analyses were conducted at 2 km resolution.

First, we examined the level of overlap between areas identified as critical (through optimization described in Section 22 above) for the 12 "local" NCP vs. the 2 global climate NCP. We made three rasters from the two critical natural asset masks (1/0 masks of the 90% target for the local NCP optimized by country and the global NCP optimized globally): showing where local NCP are optimized and global NCP are not, where global NCP are optimized and local NCP are not, and where both are optimized (the overlap).

We also considered how much biodiversity and cultural diversity are represented in critical natural assets for local NCP and for global NCP (Tables S6, S7). For biodiversity, we used data on terrestrial vertebrate species AOH, described above, and calculated the percentage of species AOH "represented" in critical natural assets. Since some species are widely distributed, and some species are narrowly distributed, "represented" was defined differently for species with different sizes of AOH. For biodiversity representation targets, we followed previous studies (Hanson et al. 2020, Larsen et al. 2011, Rodrigues et al. 2004) by setting minimum extent thresholds for suitable habitat using Area of Habitat (AOH) data for mammals, reptiles, amphibians, and birds (Brooks et al. 2019). Briefly, we assigned a 100% area target to species with less than 1,000 km$^2$ of total AOH, a 10% target to species with more than 250,000 km$^2$ of AOH, and log-linearly interpolated target for species with total AOH between 1,000 and 250,000 km$^2$. We also assigned a cap of 1,000,000 km$^2$ for species with very large AOH (i.e. >10,000,000 km$^2$). We reported this for birds, mammals, reptiles, amphibians and all species globally, and broken down into endemic species (AOH 100% within country) and species whose AOH falls >50% within a country to report at the country level (SI Table 3).

To evaluate how much cultural diversity is represented by priority areas for NCP, we used unique languages as a proxy for cultural diversity, following Gorenflo et al., 2012. As noted in Section 16, the language dataset, based on 23rd edition of languages reported in *Ethnologue* (Eberhard et al. 2020), contains information on 7,117 languages, though this analysis focuses on the 6,474 languages for which we have polygon representations of language ranges. To analyze the overlap with important areas for NCP, we counted the number of polygons representing unique indigenous and non-migrant languages that intersect with the critical natural assets for local NCP and for global NCP (SI Table 4). (Due to processing time, we didn't perform this analysis for all 20 target levels.)

### 24. Modeling limitations and data issues

##### *Effect of masking and resolution on percent of natural habitat*

Our original NCP layers were produced at a native resolution of 10 arc seconds (~300 m) or resampled to that resolution by masking to relevant natural habitat using ESA LULC data (see Extended Data Table 3). These layers were then resampled to an equal-area projection (Eckert) at 2km for computational feasibility of the optimization. We resampled using “near” rather than “average” or “mode” because this gave us the most consistently similar patterns and amounts of natural assets. We compare results of masking using near, average, mode at different resolutions (300 m, 2km, 4km, 10km) for calculating % natural assets in Table SM24.1 below.

**Table SM24.1** Percent natural assets under different resampling methods and resolutions

| Resampling method | Resolution | Percent natural assets |
|---|---|---|
| Average | 2 km | 79.4% |
| Mode | 2 km | 65.0% |
| Near | 2 km | 63.9% |
| Near | 4 km | 61.0% |
| Near | 10 km | 64.5% |

##### *No data values*

In our NCP maps, “no data” indicates that data is either 1) not attributed to that realm (for example, a terrestrial ecosystem service will show “no data” in the ocean, and vice versa), 2) unavailable for that location due to missing model inputs (most notably, the DEM used for nitrogen and sediment retention does not extend past 60 N; and data on flood mitigation, timber, fuelwood and grazing were not available for small island nations), masked from that location due to our definition of “natural assets” (i.e., excluding cropland, urban and bare areas, and permanent ice and snow).  “Zero” values indicate that the NCP could potentially be present in that location, but is not, either because the ecosystem doesn’t supply that particular NCP, or because there are no people benefitting from it (or both.) Each NCP map has different definitions of NCP supply and beneficiaries; therefore, “no data” and “zero” values can differ between individual maps. In the resulting (optimized) layers, “no data” indicates that none of the NCP maps have data for that location; “zero” indicates that a location did not contribute to the defined target level of any of the NCPs.

##### *Skewed and zero-inflated data*

Many of the NCP maps are both highly left-skewed and “zero-inflated,” meaning that for much of the planet’s area there is no or low provision of NCP and fewer places on the planet provide much higher levels of NCP. This is because ecosystems (and their contributions to people) are not evenly distributed across the planet, and also because beneficiaries are not evenly distributed. Thus, some ecosystems provide disproportionately higher levels of benefits than others, and some ecosystems provide benefits to vastly larger numbers of beneficiaries than others. For example, only 5% of marine grid cells provide 90% of marine fish catch, and the figures for riverine fish catch are quite similar. Fish, and the people catching them, tend to be concentrated over relatively small areas. This represents an opportunity: since more benefits are provided by relatively less area, it is possible to conserve the most important areas and maintain a disproportionate level of benefits from nature to humanity. It also presents a risk: extremely important benefits could be lost by inadequately maintaining a fairly small geographic area.

For the optimization, we wanted to retain this underlying information (the information about which areas provide disproportionate benefits.) Therefore, we selected linear integer programming which identifies areas that are providing a target level of each NCP across multiple maps, without having to transform the underlying maps (that is, the optimization handles highly skewed and zero-inflated data).

#### *Outliers*

Several of our datasets had outlier values which were a byproduct of modeling or post-processing. For a very small number of pixels, some models would produce nonsensical (e.g., negative, extremely low, or extremely high) values. For example, in Co$ting Nature, to avoid division by zero values, the model adds a very small number (0.000001) to the denominator in some of the final indices. These values should thus be ignored. Other small values (>0.000001) are meaningful, as they reflect that most of the world's land area has low human population densities (deserts, forests, snow and ice, rural). We consulted with the data providers to address these outliers, re-assigning negative values to zero, extremely small values to zero, or "capping" extremely high values at a maximum value, as appropriate for each case. Table SM24.2 provides the re-setting or capping applied to each layer.

**Table SM24.2** Data truncation rules applied to each dataset.

| Raster layer(s) | Value | Reset to | Rationale |
|---|---|---|---|
| Timber, fuelwood, fodder for livestock, flood mitigation | <0.001 | 0 | Values below this point are model artifacts to avoid divide by 0 errors |
| Freshwater fish | <0.001 | 0 | Minimum values are regression artifacts. Should be cut off at 0.001 T (1 kg) per $km^2$ |
| (Potential*) Nitrogen retention | >322 | 322 | Model produces rare but extreme outliers; 322 is the 99.9th percentile value for the global output |
| (Potential*) Sediment retention | >161 | 161 | Model produces rare but extreme outliers; 161is the 99.9th percentile value for the global output |
| Pollination | >38 | 38 | Model produces rare but extreme outliers; 38 is the 99.9th percentile value for the global output |

**for sediment and nitrogen retention the capping was done on the biophysical layer before combining with the downstream beneficiaries layer*

#### *Philosophical considerations*

There are a number of philosophical considerations in how and which values were ascribed for any given NCP. First, the approach we took of multiplying nature's contribution by the number of people benefitting presumes a world view that "the greatest good for the greatest number of people" has the highest value. However, high-dependence low-population places may be considered by many to be more deserving of attention for conservation. Places that support only a small number of people may still be irreplaceable for those few people (e.g., remote Indigenous tribes), due to their lack of access to substitutes or lack of capacity to substitute for many relational values. Second, the framing of *nature's contributions to people* is an anthropocentric worldview that fails to capture the intrinsic values of biodiversity, but nature may still be providing important contributions to biodiversity that are not captured by biodiversity maps alone. Regulating contributions in particular, including water quality regulation, natural hazards resilience, pollination, and atmospheric moisture recycling, maintain the conditions under which current biodiversity thrives. Delineating species or high biodiversity areas as the "beneficiaries" for all of these contributions may be an important step toward reflecting the intrinsic values of nature's contributions. Finally, applying equal targets to all NCP makes the implicit assumption that the values of all contributions are equal. In a stakeholder-led process, targets should be set according to importance ascribed by the people relying on nature's contributions, which may make certain areas within our "critical" natural assets more critical than others. As noted in the main text, our approach described here is really just a template for countries or decision-makers at the appropriate level of governance to follow to help identify conservation priorities, with input reflecting the values and knowledge of local people.

### References


Baccini, A. et al., Estimated carbon dioxide emissions from tropical deforestation improved by carbon-density maps. Nat. Clim. Chang. 2, 182–185 (2012).

Banks, D. I., et al., Wood supply and demand around two rural settlements in a semi-arid Savanna, South Africa. Biomass and Bioenergy. 11, 319–331 (1996).

BirdLife International and Handbook of the Birds of the World, "Bird species distribution maps of the world" (BirdLife International, Cambridge, UK, 2019).

Brooks, T. M., et al., Measuring terrestrial area of habitat (AOH) and its utility for the IUCN Red List. Trends Ecol. Evol. 34, 977–986 (2019).

Buchhorn, M., et al., Copernicus Global Land Service: Land Cover 100m: epoch 2015: Globe (2019), (available at https://zenodo.org/record/3243509).

Bunting, P., et al., The Global Mangrove Watch—A New 2010 Global Baseline of Mangrove Extent. Remote Sens. 10, 1669 (2018).

Burke, L., K. Reytar, M. Spalding, A. Perry, "Reefs at Risk Revisited" (World Resources Institute, 2011), (available at http://www.wri.org/publication/reefs-risk-revisited).

Byrd, K. B., et al., A remote sensing-based model of tidal marsh aboveground carbon stocks for the conterminous United States. ISPRS J. Photogramm. Remote Sens. 139, 255–271 (2018).

Chaplin-Kramer, R., et al., Global modeling of nature's contributions to people. Science (80-. ). 366, 255–258 (2019).

de Souza, E. N., K. Boerder, S. Matwin, B. Worm, Improving Fishing Pattern Detection from Satellite AIS Using Data Mining and Machine Learning. PLoS One. 11, e0158248 (2016).

Dee, D. P., et al., The ERA-Interim reanalysis: configuration and performance of the data assimilation system. Q. J. R. Meteorol. Soc. 137, 553–597 (2011).

Dijkstra, E.W. (1959). A note on two problems in connexion with graphs. Numerische Mathematik, 1(1), 269–271

Eberhard, D.M., G.F. Simons, and C.D. Fennig (eds.), 2020. *Ethnologue, Languages of the World*. 23rd edition, SIL International, Dallas, Texas. Online version: http://www.ethnologue.com.

Ellis, E. C., N. Ramankutty, Putting people in the map: anthropogenic biomes of the world. Front. Ecol. Environ. 6, 439–447 (2008).

EPA, "Methodology for assessing health risks associated with indirect exposure to combuster emissions" (1990), (available at https://ntrl.ntis.gov/NTRL/dashboard/searchResults.xhtml?searchQuery=PB90187055&starDB=GRAHIST#).

Fick, S. E., R. J. Hijmans, WorldClim 2: new 1-km spatial resolution climate surfaces for global land areas. Int. J. Climatol. 37, 4302–4315 (2017).

Fluet-Chouinard, E., S. Funge-Smith, P. B. McIntyre, Global hidden harvest of freshwater fish revealed by household surveys. Proc. Natl. Acad. Sci. 115, 7623–7628 (2018).

Fluet-Chouinard, E., et al., Development of a global inundation map at high spatial resolution from topographic downscaling of coarse-scale remote sensing data. Remote Sens. Environ. 158, 348–361 (2015).

J. W. Fourqurean et al., Seagrass ecosystems as a globally significant carbon stock. Nat. Geosci. 5, 505–509 (2012).

A. Goldstein et al., Protecting irrecoverable carbon in Earth's ecosystems. Nat. Clim. Chang. 10, 287–295 (2020).

L. J. Gorenflo, S. Romaine, R. A. Mittermeier, K. Walker-Painemilla, Co-occurrence of linguistic and biological diversity in biodiversity hotspots and high biodiversity wilderness areas. Proc. Natl. Acad. Sci. 109, 8032–8037 (2012).

Greulich, F. R., D. P. Hanley, J. F. McNeel, D. Baumgartner, "A Primer for Timber Harvesting" (Cooperative Extension, Washington State University, 1999), (available at http://faculty.washington.edu/greulich/Documents/eb1316.pdf).

Gunnell, K., M. Mulligan, R. A. Francis, D. G. Hole, Evaluating natural infrastructure for flood management within the watersheds of selected global cities. Sci. Total Environ. 670, 411–424 (2019).

Hanson, J. O., et al., Global conservation of species' niches. Nature. 580, 232–234 (2020).

Hanson, J. O., et al., Systematic Conservation Prioritization in R (2020), (available at https://cran.r-project.org/package=prioritizr).

Hengl, T., et al., SoilGrids250m: Global gridded soil information based on machine learning. PLoS One. 12, e0169748 (2017).

Hurlbert, A. H., W. Jetz, Species richness, hotspots, and the scale dependence of range maps in ecology and conservation. Proc. Natl. Acad. Sci. 104, 13384–13389 (2007).

Hutchison, J., et al., Predicting Global Patterns in Mangrove Forest Biomass. Conserv. Lett. 7, 233–240 (2014).

Copernicus Services Information, "Copernicus Global Land Operations "Vegetation and Energy""" (2019), (available at https://land.copernicus.eu/global/products/dmp).

Flanders Marine Institute, "Union of the ESRI Country shapefile and the Exclusive Economic Zones (version 3)" (2020), (available at https://www.marineregions.org/. https://doi.org/10.14284/403).

ISRIC, "Soil organic carbon stock (mean 0-30cm depth)" (2019), (available at https://soilgrids.org/).

IUCN, “A Global Standard for the Identification of Key Biodiversity Areas, Version 1.0” (IUCN, Gland, Switzerland, 2016), (available at https://portals.iucn.org/library/sites/library/files/documents/2016-048.pdf).

Keys, P. W., L. Wang-Erlandsson, L. J. Gordon, Revealing Invisible Water: Moisture Recycling as an Ecosystem Service. PLoS One. 11, e0151993 (2016).

Klein, A.-M., et al., Importance of pollinators in changing landscapes for world crops. Proc. R. Soc. B Biol. Sci. 274, 303–313 (2007).

Kremen, C., et al., The area requirements of an ecosystem service: crop pollination by native bee communities in California. Ecol. Lett. 7, 1109–1119 (2004).

Larsen, F. W., M. C. Londoño- Murcia, W. R. Turner, Global priorities for conservation of threatened species, carbon storage, and freshwater services: scope for synergy? Conserv. Lett. 4, 355–363 (2011).

Lehner, B., G. Grill, Global river hydrography and network routing: baseline data and new approaches to study the world’s large river systems: GLOBAL RIVER HYDROGRAPHY AND NETWORK ROUTING. Hydrol. Process. 27, 2171–2186 (2013).

Lehner, B., K. Verdin, A. Jarvis, New Global Hydrography Derived From Spaceborne Elevation Data. Eos, Trans. Am. Geophys. Union. 89, 93–94 (2008).

IUCN Red List, UNEP_WCMC, “Species Range Polygons” (2020), (available at https://www.iucnredlist.org/resources/other-spatial-downloads).

McIntyre, P. B., C.A.R. Liermann, C. Revenga, Linking freshwater fishery management to global food security and biodiversity conservation. Proc. Natl. Acad. Sci. 113, 12880–12885 (2016).

Mcowen, C., et al., A global map of saltmarshes. Biodivers. Data J. 5, e11764 (2017).

Monfreda, C., N. Ramankutty, J. A. Foley, Farming the planet: 2. Geographic distribution of crop areas, yields, physiological types, and net primary production in the year 2000. Global Biogeochem. Cycles. 22, GB1022 (2008).

Mulligan, M, “Documentation for the Co$tingNature Model V3” (King’s College London, London, 2018), (available at https://docs.google.com/document/d/136OvAO6PSyVBp0gNl9f0_pIAIg-h-4JZGArnf2V6-Us/edit).

Mulligan, M, “Documentation for the WaterWorld Model V2.” (King’s College London, London, 2016), (available at https://docs.google.com/document/d/1GKheQFp5_rsZyazwCJxCzeEStQF2jh04x-Dl_oG5yoY/edit).

Nelson, A, “Estimated travel time to the nearest city of 50,000 or more people in year 2000” (Global Environment Monitoring Unit-Joint Research Centre of the European Commission, Ispra Italy, 2008), (available at http://http//bioval. jrc. ec. europa. eu/products/gam/).

CIESIN, “Gridded Population of the World, Version 4 (GPWv4): Data Quality Indicators, Revision 11” (Columbia University and NASA Socioeconomic Data and Applications Center (SEDAC), Palisades, NY, 2017), (available at https://doi.org/10.7927/H42R3PN3.).

OECD, “World energy outlook 2006. Chapter 15 Energy for Cooking in Developing Countries” (OECD/IEA, 2006), (available at https://www.iea.org/reports/world-energy-outlook-2006).

Pauly, D., D. Zeller, Catch reconstructions reveal that global marine fisheries catches are higher than reported and declining. Nat. Commun. 7, ncomms10244 (2016).

Pauly, D., Zeller, D. & Palomares, M. D. Sea Around Us Concepts, Design and Data (2020). Available at: seaaroundus. org.

Porter, C., et al., “ArcticDEM v7” (Harvard Dataverse, V1, 2018), (available at https://doi.org/10.7910/DVN/OHHUKH).

ESA CCI project, “Land Cover CCI” (European Space Agency Climate Change Initiative, Belgium, 2017), (available at http://maps.elie.ucl.ac.be/CCI/viewer/download/ESACCI-LC-Ph2-PUGv2_2.0.pdf).

Roberson, L. A., R. A. Watson, C. J. Klein, Over 90 endangered fish and invertebrates are caught in industrial fisheries. Nat. Commun. 11, 4764 (2020).

Robinson, T. P., et al., Mapping the global distribution of livestock (2014).

Rodrigues, A. S. L., et al., Global Gap Analysis: Priority Regions for Expanding the Global Protected-Area Network. Bioscience. 54, 1092–1100 (2004).

Rose, A. N., J. J. McKee, M. L. Urban, E. A. Bright, LandScan 2017 (2018), (available at https://landscan.ornl.gov/).

Sanderman, J., et al., A global map of mangrove forest soil carbon at 30 m spatial resolution. Environ. Res. Lett. 13, 55002 (2018).

Santini, L., et al., Applying habitat and population-density models to land-cover time series to inform IUCN Red List assessments. Conserv. Biol. 33, 1084–1093 (2019).

Schneider, A., M. A. Friedl, D. Potere, A new map of global urban extent from MODIS satellite data. Environ. Res. Lett. 4, 44003 (2009).

Schuster, R., et al., Optimizing the conservation of migratory species over their full annual cycle. Nat. Commun. 10, 1754 (2019).

Selig, E. R., et al., Mapping global human dependence on marine ecosystems. Conserv. Lett. 12, e12617 (2019).

Sharp, R., et al., “InVEST 3.8.0 User’s Guide” (The Natural Capital Project, Stanford University, University of Minnesota, The Nature Conservancy, and World Wildlife Fund, 2020), (available at http://releases.naturalcapitalproject.org/invest-userguide/latest/).

Sharp, R., et al., “InVEST 3.5.0 User’s Guide” (The Natural Capital Project, Stanford University, University of Minnesota, The Nature Conservancy, and World Wildlife Fund, 2018), (available at http://data.naturalcapitalproject.org/invest-releases/3.5.0/userguide/index.html).

SIL, International. n.d. Ethnologue. Languages of the World. https://www.ethnologue.com/

Simard, M., et al., Global Mangrove Distribution, Aboveground Biomass, and Canopy Height. ORNL DAAC (2019), doi:https://doi.org/10.3334/ORNLDAAC/1665.

Soto-Navarro, C., et al., Mapping co-benefits for carbon storage and biodiversity to inform conservation policy and action. Philos. Trans. R. Soc. B Biol. Sci. 375, 20190128 (2020).

Spalding, M., et al., Mapping the global value and distribution of coral reef tourism. Mar. Policy. 82, 104–113 (2017).

Spawn, S. A., C. C. Sullivan, T. J. Lark, H. K. Gibbs, Harmonized global maps of above and belowground biomass carbon density in the year 2010. Sci. Data. 7, 112 (2020).

Stürck, J., A. Poortinga, P. H. Verburg, Mapping ecosystem services: The supply and demand of flood regulation services in Europe. Ecol. Indic. 38, 198–211 (2014).

Turner, W. R., D. S. Wilcove, H. M. Swain, Assessing the Effectiveness of Reserve Acquisition Programs in Protecting Rare and Threatened Species. Conserv. Biol. 20, 1657–1669 (2006).

FAO and UNEP, “The State of the World’s Forests 2020. Forests, biodiversity and people” (Rome, 2020), (available at https://doi.org/10.4060/ca8642en).

UNEP-WCMC, F. T. Short, “Global distribution of seagrasses (version 6.0)” (UN Environment World Conservation Monitoring Centre, Cambridge, UK, 2017), (available at http://data.unepwcmc.org/datasets/7).

Vafeidis, A. T., et al., A New Global Coastal Database for Impact and Vulnerability Analysis to Sea-Level Rise. J. Coast. Res. 24, 917–924 (2008).

van der Ent, R. J., L. Wang-Erlandsson, P. W. Keys, H. H. G. Savenije, Contrasting roles of interception and transpiration in the hydrological cycle – Part 2: Moisture recycling. Earth Syst. Dyn. 5, 471–489 (2014).

Wang-Erlandsson, L., R. J. van der Ent, L. J. Gordon, H. H. G. Savenije, Contrasting roles of interception and transpiration in the hydrological cycle – Part 1: Temporal characteristics over land. Earth Syst. Dyn. 5, 441–469 (2014).

Watson, R. A., A. Tidd, Mapping nearly a century and a half of global marine fishing: 1869–2015. Mar. Policy. 93, 171–177 (2018).

World Bank, Forests (2020), (available at https://www.worldbank.org/en/topic/forests).
