## Supplementary material for "Mapping the planet’s critical natural assets": SI Tables

### SI Table 1. Percentage of land and waters in critical natural assets for different analyses.

| Country (ranked by land area) | Total area (sq km) | Area of land (sq km) | Area of EEZ (sq km) | % of total area that is critical natural assets for local NCP (LCNA) | % of land that is natural assets | % of land that is LCNA | % of natural assets that are LCNA | % eez that is LCNA | % of land critical for global NCP (GCNA) | % overlap between LCNA and GCNA | % of LCNA not over-lapping with GCNA | % of GCNA not over-lapping with LCNA | Total % of land that contains LCNA and GCNA | LCNA % area selected across all 12 "drop 1" optimiza tions | LCNA % area selected across 11 of 12 "drop 1" optimiza tions |
|---|---|---|---|---|---|---|---|---|---|---|---|---|---|---|---|
| Global | 274,470,564 | 134,761,492 | 139,709,072 | 27.3% | 63.9% | 30.4% | 47.6% | 24.2% | 38.5% | 23.7% | 6.3% | 14.3% | 44.4% | 61.9% | 96.9% |
| Russia | 24,615,504 | 16,924,348 | 7,691,156 | 18.5% | 86.2% | 17.8% | 20.7% | 20.1% | 35.8% | 14.9% | 2.7% | 20.4% | 38.0% | 57.3% | 96.2% |
| Canada | 15,661,972 | 9,917,508 | 5,744,464 | 13.0% | 88.0% | 10.7% | 12.2% | 16.9% | 27.8% | 8.8% | 1.7% | 15.7% | 26.2% | 59.5% | 98.3% |
| United States of America | 21,628,792 | 9,467,504 | 12,161,288 | 24.7% | 75.8% | 37.4% | 49.3% | 14.8% | 52.9% | 34.1% | 3.0% | 18.1% | 55.2% | 56.5% | 96.4% |
| China | 10,701,884 | 9,393,892 | 1,307,992 | 29.3% | 51.3% | 25.4% | 49.5% | 57.2% | 38.1% | 20.0% | 5.1% | 18.0% | 43.1% | 54.9% | 96.9% |
| Brazil | 12,222,380 | 8,523,920 | 3,698,460 | 38.8% | 74.7% | 46.0% | 61.6% | 22.2% | 70.5% | 42.5% | 3.4% | 28.0% | 73.8% | 62.4% | 96.1% |
| Australia | 16,745,092 | 7,723,376 | 9,021,716 | 23.5% | 87.1% | 24.4% | 28.0% | 22.7% | 10.5% | 5.6% | 18.6% | 4.2% | 28.4% | 25.4% | 98.1% |
| India | 5,503,556 | 3,165,956 | 2,337,600 | 18.9% | 28.6% | 19.4% | 67.9% | 18.3% | 19.9% | 15.3% | 3.5% | 4.6% | 23.4% | 71.7% | 94.9% |
| Argentina | 3,863,420 | 2,790,952 | 1,072,468 | 50.0% | 71.4% | 44.2% | 61.9% | 65.4% | 23.0% | 18.8% | 25.2% | 4.3% | 48.2% | 48.3% | 97.1% |
| Kazakhstan | 2,826,860 | 2,712,492 | 114,368 | 33.8% | 64.3% | 35.2% | 54.8% | 0.3% | 55.3% | 30.9% | 3.8% | 24.2% | 58.8% | 40.4% | 94.8% |
| Democratic Republic of the Congo | 2,354,184 | 2,340,724 | 13,460 | 70.3% | 88.2% | 70.6% | 80.0% | 25.1% | 86.8% | 70.2% | 0.4% | 16.6% | 87.2% | 80.5% | 98.3% |
| Algeria | 2,448,776 | 2,317,420 | 131,356 | 4.1% | 5.1% | 2.8% | 54.8% | 27.4% | 3.0% | 1.8% | 0.9% | 1.2% | 3.9% | 69.8% | 95.4% |
| Greenland | 2,142,460 | 2,142,460 | - | 0.6% | 16.6% | 0.0% | 0.0% | | 0.0% | | | | 0.0% | | |
| Mexico | 5,168,872 | 1,966,228 | 3,202,644 | 29.0% | 81.1% | 49.1% | 60.6% | 16.6% | 33.0% | 24.3% | 24.5% | 8.7% | 57.5% | 55.9% | 96.9% |
| Saudi Arabia | 2,155,684 | 1,930,412 | 225,272 | 7.5% | 4.0% | 2.2% | 53.9% | 53.6% | 0.0% | 0.0% | 0.4% | 0.0% | 0.4% | 51.4% | 96.9% |
| Indonesia | 7,952,676 | 1,892,408 | 6,060,268 | 37.1% | 63.1% | 33.0% | 52.2% | 38.3% | 58.0% | 27.9% | 3.3% | 30.1% | 61.3% | 60.7% | 96.3% |
| Sudan | 1,950,240 | 1,867,248 | 82,992 | 20.2% | 26.7% | 18.6% | 69.4% | 57.1% | 14.5% | 12.0% | 6.5% | 2.5% | 21.0% | 75.1% | 96.6% |
| Libya | 1,995,616 | 1,630,216 | 365,400 | 4.5% | 2.6% | 0.8% | 32.6% | 20.9% | 0.0% | 0.0% | 0.2% | 0.0% | 0.2% | 51.0% | 98.2% |
| Iran | 1,843,292 | 1,627,164 | 216,128 | 12.6% | 14.8% | 8.8% | 59.6% | 41.4% | 11.1% | 7.3% | 1.2% | 3.7% | 12.2% | 59.6% | 97.1% |
| Mongolia | 1,563,920 | 1,563,920 | - | 29.7% | 49.4% | 29.7% | 60.2% | | 29.3% | 20.1% | 9.6% | 9.2% | 38.9% | 61.5% | 97.2% |
| Peru | 2,158,176 | 1,298,040 | 860,136 | 39.2% | 88.4% | 57.2% | 64.6% | 12.1% | 75.3% | 48.8% | 8.1% | 26.5% | 83.4% | 61.2% | 97.0% |
| Chad | 1,273,508 | 1,273,508 | - | 21.5% | 31.2% | 21.5% | 68.8% | | 14.5% | 12.2% | 9.3% | 2.3% | 23.8% | 63.2% | 99.6% |
| Mali | 1,259,524 | 1,259,524 | - | 18.5% | 23.4% | 18.5% | 79.2% | | 4.1% | 3.9% | 14.5% | 0.3% | 18.6% | 73.4% | 98.2% |
| Angola | 1,750,540 | 1,251,856 | 498,684 | 51.7% | 91.7% | 65.9% | 71.9% | 16.2% | 89.4% | 64.8% | 1.0% | 24.6% | 90.3% | 71.8% | 99.2% |
| South Africa | 2,774,500 | 1,224,140 | 1,550,360 | 31.5% | 84.8% | 59.3% | 69.9% | 9.5% | 12.7% | 9.4% | 49.8% | 3.3% | 62.5% | 59.6% | 99.3% |
| Niger | 1,187,828 | 1,187,828 | - | 18.2% | 24.3% | 18.2% | 74.8% | | 0.1% | 0.1% | 17.4% | 0.0% | 17.6% | 70.7% | 99.1% |
| Colombia | 1,866,912 | 1,142,680 | 724,232 | 33.6% | 87.3% | 47.9% | 54.8% | 11.0% | 85.5% | 46.4% | 1.3% | 39.1% | 86.8% | 67.9% | 96.2% |
| Ethiopia | 1,134,556 | 1,134,556 | - | 47.3% | 69.4% | 47.3% | 68.1% | | 61.5% | 43.1% | 4.1% | 18.4% | 65.6% | 61.9% | 97.7% |
| Bolivia | 1,092,884 | 1,092,884 | - | 60.5% | 84.6% | 60.5% | 71.6% | | 82.8% | 60.4% | 0.1% | 22.4% | 82.9% | 70.4% | 95.8% |
| Mauritania | 1,216,088 | 1,041,976 | 174,112 | 13.2% | 16.6% | 11.7% | 70.2% | 22.4% | 0.0% | 0.0% | 11.2% | 0.0% | 11.3% | 71.7% | 97.4% |
| Egypt | 1,250,212 | 1,006,044 | 244,168 | 2.6% | 3.2% | 0.7% | 22.3% | 10.4% | 0.1% | 0.1% | 0.2% | 0.0% | 0.3% | 39.7% | 97.6% |
| United Republic of Tanzania | 1,190,740 | 947,488 | 243,252 | 39.7% | 71.4% | 45.8% | 64.1% | 16.1% | 63.2% | 43.3% | 2.1% | 19.9% | 65.3% | 67.4% | 96.6% |
| Venezuela | 1,395,436 | 918,704 | 476,732 | 50.2% | 83.9% | 60.3% | 71.8% | 30.7% | 79.8% | 57.1% | 3.0% | 22.7% | 82.8% | 77.3% | 96.7% |
| Nigeria | 1,093,508 | 913,232 | 180,276 | 27.0% | 35.2% | 27.0% | 76.7% | 27.0% | 31.3% | 25.4% | 1.5% | 5.9% | 32.8% | 77.6% | 95.4% |
| Pakistan | 1,101,288 | 875,924 | 225,364 | 21.0% | 38.2% | 20.7% | 54.3% | 22.2% | 16.3% | 12.7% | 7.7% | 3.5% | 23.9% | 63.1% | 95.5% |
| Namibia | 1,391,648 | 826,760 | 564,888 | 46.2% | 78.1% | 56.5% | 72.3% | 31.2% | 34.0% | 25.3% | 31.1% | 8.7% | 65.0% | 50.6% | 98.8% |
| Mozambique | 1,361,736 | 792,764 | 568,972 | 43.1% | 84.6% | 61.8% | 73.1% | 16.9% | 72.9% | 53.1% | 8.5% | 19.8% | 81.4% | 72.0% | 96.1% |
| Turkey | 1,043,536 | 781,076 | 262,460 | 34.0% | 41.2% | 30.7% | 74.4% | 44.1% | 36.4% | 27.0% | 3.2% | 9.4% | 39.6% | 67.2% | 93.2% |
| Zambia | 756,488 | 756,488 | - | 68.3% | 83.0% | 68.3% | 82.3% | | 81.7% | 68.2% | 0.1% | 13.5% | 81.9% | 80.9% | 97.3% |
| Chile | 4,414,404 | 737,852 | 3,676,552 | 9.5% | 68.9% | 24.8% | 35.9% | 6.5% | 36.7% | 12.5% | 11.2% | 17.3% | 41.0% | 52.2% | 96.2% |
| Myanmar | 1,166,872 | 666,444 | 500,428 | 46.7% | 65.6% | 48.6% | 74.1% | 44.1% | 64.3% | 47.4% | 0.6% | 16.9% | 64.8% | 71.5% | 96.6% |
| Afghanistan | 643,804 | 643,804 | - | 27.7% | 48.4% | 27.7% | 57.2% | | 45.8% | 27.3% | 0.3% | 18.6% | 46.2% | 64.1% | 95.6% |
| France | 10,744,036 | 637,004 | 10,107,032 | 5.8% | 52.2% | 33.6% | 64.5% | 4.0% | 46.1% | 28.4% | 4.0% | 17.7% | 50.1% | 65.6% | 95.8% |
| South Sudan | 630,920 | 630,920 | - | 79.1% | 94.6% | 79.1% | 83.6% | | 94.6% | 79.0% | 0.0% | 15.6% | 94.6% | 88.4% | 98.9% |
| Central African Republic | 622,068 | 622,068 | - | 80.8% | 97.9% | 80.8% | 82.5% | | 97.9% | 80.8% | 0.0% | 17.1% | 97.9% | 81.6% | 98.5% |

| | | | | | | | | | | | | | | | |
|---|---|---|---|---|---|---|---|---|---|---|---|---|---|---|---|
| Madagascar | 1,842,036 | 596,124 | 1,245,912 | 32.4% | 84.9% | 65.0% | 76.6% | 16.8% | 64.4% | 48.5% | 15.9% | 16.0% | 80.4% | 69.8% | 95.6% |
| Morocco | 874,924 | 593,764 | 281,160 | 22.0% | 22.5% | 16.0% | 71.2% | 34.6% | 7.2% | 6.6% | 8.3% | 0.5% | 15.4% | 69.2% | 97.7% |
| Kenya | 754,488 | 589,712 | 164,776 | 39.8% | 71.9% | 48.2% | 67.0% | 9.9% | 65.8% | 46.2% | 1.8% | 19.6% | 67.6% | 55.4% | 98.4% |
| Botswana | 581,924 | 581,924 | - | 54.2% | 92.4% | 54.2% | 58.6% | | 56.7% | 29.5% | 24.7% | 27.2% | 81.4% | 47.9% | 98.7% |
| Ukraine | 707,652 | 571,372 | 136,280 | 23.3% | 20.0% | 16.3% | 81.4% | 52.6% | 18.0% | 15.2% | 0.6% | 2.8% | 18.6% | 74.7% | 94.9% |
| Thailand | 818,032 | 517,484 | 300,548 | 36.9% | 31.4% | 22.5% | 71.4% | 61.8% | 29.5% | 20.9% | 0.9% | 8.6% | 30.5% | 70.8% | 96.4% |
| Spain | 1,517,452 | 507,476 | 1,009,976 | 27.5% | 43.7% | 35.5% | 81.3% | 23.5% | 32.1% | 25.1% | 7.8% | 6.9% | 39.9% | 64.4% | 93.9% |
| Somalia | 1,262,120 | 474,972 | 787,148 | 29.4% | 83.0% | 61.1% | 73.6% | 10.3% | 58.4% | 47.8% | 13.0% | 10.6% | 71.4% | 65.2% | 98.6% |
| Turkmenistan | 532,768 | 471,460 | 61,308 | 10.4% | 21.0% | 11.8% | 56.2% | 0.1% | 13.3% | 10.0% | 1.7% | 3.3% | 15.0% | 48.5% | 98.9% |
| Papua New Guinea | 2,887,076 | 468,136 | 2,418,940 | 67.2% | 86.4% | 52.3% | 60.5% | 70.1% | 84.8% | 49.8% | 1.1% | 34.8% | 85.7% | 72.5% | 96.2% |
| Cameroon | 482,652 | 467,344 | 15,308 | 62.7% | 84.1% | 62.5% | 74.2% | 71.0% | 83.1% | 61.8% | 0.6% | 21.3% | 83.7% | 56.0% | 95.8% |
| Yemen | 986,256 | 455,628 | 530,628 | 14.2% | 16.1% | 9.9% | 61.5% | 17.8% | 0.0% | 0.0% | 7.2% | 0.0% | 7.3% | 70.2% | 96.2% |
| Uzbekistan | 448,200 | 448,200 | - | 11.3% | 23.1% | 11.3% | 48.8% | | 13.6% | 10.1% | 1.1% | 3.5% | 14.7% | 58.3% | 97.9% |
| Sweden | 600,012 | 444,624 | 155,388 | 43.6% | 89.9% | 33.0% | 36.7% | 73.7% | 78.6% | 30.9% | 1.4% | 47.7% | 80.0% | 64.5% | 96.5% |
| Iraq | 439,772 | 438,576 | 1,196 | 1.8% | 3.0% | 1.5% | 51.5% | 84.3% | 1.1% | 0.9% | 0.3% | 0.2% | 1.4% | 62.9% | 94.9% |
| Paraguay | 401,704 | 401,704 | - | 62.6% | 77.1% | 62.6% | 81.2% | | 76.6% | 62.4% | 0.2% | 14.2% | 76.8% | 80.8% | 99.8% |
| Zimbabwe | 391,384 | 391,384 | - | 49.6% | 62.3% | 49.6% | 79.5% | | 61.3% | 49.2% | 0.3% | 12.1% | 61.6% | 77.1% | 93.9% |
| Norway | 2,391,520 | 380,476 | 2,011,044 | 42.5% | 74.7% | 25.7% | 34.4% | 45.7% | 37.3% | 19.7% | 3.7% | 17.2% | 40.6% | 42.4% | 96.7% |
| Japan | 4,669,500 | 374,096 | 4,295,404 | 17.3% | 68.1% | 48.3% | 71.0% | 14.6% | 26.2% | 16.6% | 29.5% | 9.6% | 55.7% | 65.1% | 96.6% |
| Germany | 413,740 | 357,144 | 56,596 | 43.5% | 46.9% | 38.3% | 81.7% | 76.0% | 46.3% | 37.7% | 0.3% | 8.6% | 46.6% | 77.8% | 96.4% |
| Republic of the Congo | 381,200 | 347,180 | 34,020 | 64.9% | 89.9% | 68.3% | 76.0% | 30.4% | 89.2% | 68.0% | 0.2% | 21.2% | 89.4% | 77.9% | 99.0% |
| Finland | 412,496 | 331,656 | 80,840 | 48.4% | 90.9% | 41.4% | 45.5% | 77.2% | 83.6% | 38.8% | 1.9% | 44.7% | 85.5% | 50.9% | 97.7% |
| Vietnam | 1,086,328 | 330,732 | 755,596 | 41.2% | 50.5% | 38.8% | 76.9% | 42.2% | 42.7% | 32.6% | 5.0% | 10.1% | 47.7% | 71.0% | 97.3% |
| Malaysia | 844,336 | 330,060 | 514,276 | 60.2% | 65.7% | 44.6% | 67.8% | 70.3% | 56.5% | 36.3% | 4.8% | 20.2% | 61.2% | 66.9% | 98.2% |
| Ivory Coast | 495,728 | 322,812 | 172,916 | 27.9% | 47.4% | 37.3% | 78.6% | 10.3% | 46.8% | 36.9% | 0.3% | 10.0% | 47.2% | 75.8% | 96.0% |
| Poland | 342,868 | 312,896 | 29,972 | 36.8% | 39.1% | 32.3% | 82.5% | 84.2% | 38.3% | 31.5% | 0.5% | 6.8% | 38.9% | 73.8% | 94.9% |
| Oman | 872,260 | 312,840 | 559,420 | 11.4% | 6.1% | 4.0% | 65.4% | 15.6% | 0.0% | 0.0% | 0.6% | 0.0% | 0.6% | 48.4% | 97.7% |
| Italy | 838,760 | 301,348 | 537,412 | 38.1% | 39.7% | 31.7% | 79.9% | 41.7% | 33.0% | 25.8% | 5.0% | 7.2% | 38.0% | 67.4% | 94.8% |
| Philippines | 2,277,832 | 295,028 | 1,982,804 | 30.0% | 35.8% | 27.2% | 76.1% | 30.4% | 25.4% | 15.7% | 6.7% | 9.7% | 32.1% | 65.1% | 96.8% |
| Burkina Faso | 274,488 | 274,488 | - | 22.6% | 27.7% | 22.6% | 81.6% | | 5.7% | 4.7% | 17.8% | 1.0% | 23.5% | 83.6% | 97.0% |
| New Zealand | 6,105,660 | 268,704 | 5,836,956 | 9.1% | 89.9% | 60.5% | 67.3% | 6.7% | 62.9% | 37.0% | 21.4% | 22.0% | 80.4% | 45.8% | 98.3% |
| Gabon | 464,852 | 261,768 | 203,084 | 45.8% | 94.7% | 66.1% | 69.7% | 19.7% | 93.9% | 65.6% | 0.3% | 28.2% | 94.1% | 76.8% | 98.5% |
| Ecuador | 1,351,112 | 256,720 | 1,094,392 | 16.3% | 81.8% | 49.4% | 60.5% | 8.5% | 76.3% | 45.4% | 3.5% | 30.9% | 79.8% | 71.9% | 98.6% |
| Guinea | 348,736 | 245,884 | 102,852 | 52.1% | 76.7% | 63.4% | 82.6% | 25.2% | 63.0% | 52.5% | 10.8% | 10.5% | 73.8% | 77.5% | 97.5% |
| Uganda | 243,632 | 243,632 | - | 31.9% | 50.7% | 31.9% | 62.9% | | 36.2% | 30.5% | 0.7% | 5.7% | 36.9% | 78.3% | 96.9% |
| United Kingdom | 6,127,288 | 243,144 | 5,884,144 | 13.2% | 63.6% | 47.9% | 75.3% | 11.7% | 51.6% | 40.7% | 5.1% | 10.9% | 56.7% | 74.1% | 98.4% |
| Ghana | 469,140 | 240,176 | 228,964 | 31.0% | 51.6% | 43.1% | 83.4% | 18.3% | 43.8% | 36.9% | 5.6% | 6.9% | 49.5% | 77.6% | 97.6% |
| Romania | 265,976 | 236,388 | 29,588 | 35.8% | 42.4% | 35.2% | 82.8% | 40.6% | 41.0% | 34.4% | 0.6% | 6.5% | 41.6% | 76.8% | 96.0% |
| Laos | 229,312 | 229,312 | - | 68.5% | 81.4% | 68.5% | 84.2% | | 80.4% | 67.7% | 0.7% | 12.6% | 81.1% | 79.7% | 96.5% |
| Guyana | 352,220 | 212,656 | 139,564 | 34.3% | 98.2% | 40.7% | 41.5% | 24.4% | 97.9% | 40.3% | 0.2% | 57.6% | 98.1% | 52.7% | 94.7% |
| Belarus | 207,080 | 207,080 | - | 38.3% | 46.3% | 38.3% | 82.8% | | 45.9% | 38.1% | 0.2% | 7.8% | 46.2% | 79.4% | 95.5% |
| Kyrgyzstan | 199,084 | 199,084 | - | 43.0% | 62.5% | 43.0% | 68.8% | | 55.3% | 41.5% | 1.2% | 13.8% | 56.5% | 69.3% | 94.8% |
| Senegal | 356,688 | 197,232 | 159,456 | 38.9% | 67.4% | 53.1% | 78.8% | 21.2% | 4.3% | 3.1% | 49.7% | 1.1% | 53.9% | 68.2% | 95.9% |
| Syria | 196,616 | 186,328 | 10,288 | 5.7% | 6.5% | 3.8% | 59.5% | 38.4% | 2.8% | 1.9% | 0.8% | 0.9% | 3.6% | 63.9% | 94.2% |
| Cambodia | 231,168 | 182,144 | 49,024 | 45.2% | 55.5% | 41.8% | 75.3% | 57.9% | 45.3% | 34.0% | 7.4% | 11.2% | 52.7% | 82.7% | 97.6% |
| Uruguay | 321,248 | 177,824 | 143,424 | 49.8% | 79.7% | 67.0% | 84.0% | 28.5% | 6.5% | 5.6% | 61.1% | 0.9% | 67.6% | 74.7% | 97.0% |
| Somaliland | 168,536 | 168,536 | - | 51.4% | 76.2% | 51.4% | 67.4% | | 7.4% | 6.2% | 44.5% | 1.2% | 51.9% | 74.7% | 98.0% |
| Tunisia | 256,980 | 157,060 | 99,920 | 35.5% | 11.9% | 8.1% | 67.9% | 78.8% | 3.5% | 3.1% | 3.4% | 0.4% | 6.9% | 55.3% | 97.1% |
| Nepal | 147,612 | 147,612 | - | 51.7% | 65.4% | 51.7% | 79.1% | | 65.2% | 51.6% | 0.1% | 13.6% | 65.3% | 78.4% | 98.3% |
| Suriname | 280,288 | 146,080 | 134,208 | 36.2% | 99.2% | 42.2% | 42.5% | 29.6% | 97.9% | 41.5% | 0.6% | 56.3% | 98.4% | 80.1% | 97.9% |
| Tajikistan | 142,464 | 142,464 | - | 33.6% | 53.7% | 33.6% | 62.6% | | 36.4% | 31.1% | 2.2% | 5.2% | 38.6% | 68.8% | 97.8% |
| Bangladesh | 250,524 | 137,544 | 112,980 | 30.0% | 23.4% | 17.8% | 75.9% | 44.9% | 15.9% | 11.6% | 2.6% | 4.4% | 18.5% | 60.1% | 97.0% |
| Greece | 615,444 | 131,648 | 483,796 | 36.9% | 54.4% | 44.6% | 82.0% | 34.8% | 41.7% | 33.9% | 8.9% | 7.8% | 50.5% | 65.7% | 95.2% |
| Nicaragua | 344,648 | 129,436 | 215,212 | 39.3% | 67.1% | 43.4% | 64.7% | 36.8% | 38.5% | 23.7% | 18.7% | 14.8% | 57.2% | 64.7% | 96.2% |
| Eritrea | 202,040 | 123,240 | 78,800 | 34.3% | 38.5% | 23.0% | 59.9% | 52.0% | 11.8% | 10.7% | 12.0% | 1.0% | 23.7% | 79.6% | 96.3% |

| | | | | | | | | | | | | | | | |
|---|---|---|---|---|---|---|---|---|---|---|---|---|---|---|---|
| North Korea | 237,080 | 122,588 | 114,492 | 40.7% | 64.1% | 48.8% | 76.2% | 32.0% | 25.8% | 19.0% | 28.9% | 6.7% | 54.6% | 70.9% | 94.7% |
| Malawi | 120,160 | 120,160 | - | 25.1% | 47.7% | 25.1% | 52.6% | | 17.3% | 14.0% | 9.8% | 3.3% | 27.1% | 80.1% | 96.8% |
| Benin | 152,516 | 116,756 | 35,760 | 37.5% | 57.8% | 45.6% | 78.7% | 11.1% | 49.6% | 40.5% | 4.8% | 9.1% | 54.4% | 79.1% | 96.4% |
| Honduras | 324,760 | 112,912 | 211,848 | 41.4% | 76.1% | 51.4% | 67.6% | 36.1% | 52.4% | 31.5% | 19.5% | 20.9% | 71.8% | 60.8% | 95.9% |
| Bulgaria | 147,612 | 112,852 | 34,760 | 31.7% | 41.2% | 32.2% | 78.1% | 30.3% | 40.0% | 31.5% | 0.6% | 8.5% | 40.6% | 69.9% | 93.8% |
| Cuba | 464,372 | 110,404 | 353,968 | 34.2% | 50.5% | 38.4% | 76.0% | 32.9% | 18.2% | 11.5% | 25.3% | 6.5% | 43.3% | 55.1% | 95.5% |
| Guatemala | 220,676 | 109,388 | 111,288 | 30.0% | 71.8% | 49.3% | 68.7% | 11.1% | 55.7% | 36.6% | 12.6% | 19.2% | 68.3% | 71.4% | 96.4% |
| Iceland | 861,860 | 102,000 | 759,860 | 39.1% | 68.3% | 21.7% | 31.7% | 41.4% | 0.0% | 0.0% | 16.4% | 0.0% | 16.4% | 27.9% | 96.5% |
| South Korea | 530,172 | 98,696 | 431,476 | 56.6% | 51.2% | 39.3% | 76.8% | 60.6% | 11.0% | 7.3% | 30.3% | 3.6% | 41.3% | 59.4% | 96.5% |
| Liberia | 349,380 | 95,896 | 253,484 | 13.7% | 46.0% | 30.8% | 67.1% | 7.2% | 45.3% | 30.0% | 0.6% | 15.2% | 45.8% | 79.3% | 97.3% |
| Hungary | 93,076 | 93,076 | - | 22.9% | 27.6% | 22.9% | 82.8% | | 26.4% | 22.2% | 0.5% | 4.2% | 26.9% | 74.3% | 97.2% |
| Portugal | 1,823,500 | 91,544 | 1,731,956 | 10.3% | 43.5% | 37.2% | 85.5% | 8.9% | 24.7% | 21.1% | 7.9% | 3.6% | 32.6% | 64.8% | 95.3% |
| Western Sahara | 90,700 | 90,700 | - | 0.0% | 0.0% | 0.0% | 100.0% | | 0.0% | 0.0% | 0.0% | 0.0% | 0.0% | 20.0% | 100.0% |
| Jordan | 89,236 | 89,140 | 96 | 1.5% | 2.4% | 1.5% | 60.9% | 83.3% | 0.8% | 0.7% | 0.0% | 0.1% | 0.8% | 60.8% | 95.1% |
| Azerbaijan | 167,012 | 86,320 | 80,692 | 13.2% | 34.1% | 25.5% | 74.7% | 0.2% | 25.7% | 21.7% | 2.7% | 4.0% | 28.4% | 68.1% | 95.4% |
| Austria | 83,992 | 83,992 | - | 59.2% | 71.4% | 59.2% | 82.9% | | 71.2% | 59.1% | 0.1% | 12.1% | 71.2% | 81.8% | 95.3% |
| Czechia | 78,736 | 78,736 | - | 39.0% | 45.7% | 39.0% | 85.4% | | 45.4% | 38.8% | 0.1% | 6.6% | 45.6% | 76.2% | 97.1% |
| Republic of Serbia | 77,532 | 77,532 | - | 35.1% | 41.0% | 35.1% | 85.6% | | 40.5% | 34.6% | 0.4% | 5.9% | 40.9% | 78.7% | 95.6% |
| Panama | 407,864 | 74,964 | 332,900 | 20.3% | 67.2% | 44.2% | 65.8% | 14.9% | 59.5% | 37.5% | 5.9% | 21.6% | 65.0% | 69.0% | 96.9% |
| Sierra Leone | 233,772 | 72,084 | 161,688 | 15.2% | 32.1% | 26.0% | 81.0% | 10.3% | 23.2% | 18.0% | 7.7% | 5.2% | 30.9% | 79.0% | 94.9% |
| United Arab Emirates | 129,352 | 71,384 | 57,968 | 35.6% | 4.7% | 4.0% | 84.8% | 74.6% | 0.0% | 0.0% | 0.3% | 0.0% | 0.3% | 61.5% | 98.2% |
| Georgia | 92,524 | 69,564 | 22,960 | 43.8% | 60.7% | 41.8% | 68.9% | 49.8% | 60.6% | 41.4% | 0.1% | 19.1% | 60.6% | 64.0% | 95.5% |
| Ireland | 495,468 | 69,244 | 426,224 | 46.0% | 90.0% | 73.8% | 82.0% | 41.4% | 75.1% | 65.0% | 6.6% | 10.2% | 81.7% | 81.1% | 98.0% |
| Sri Lanka | 603,828 | 66,740 | 537,088 | 13.0% | 49.2% | 39.5% | 80.2% | 9.7% | 32.5% | 25.3% | 12.3% | 7.2% | 44.8% | 62.2% | 97.6% |
| Lithuania | 71,656 | 64,796 | 6,860 | 40.5% | 41.5% | 35.6% | 85.6% | 87.3% | 41.0% | 35.0% | 0.4% | 6.0% | 41.4% | 75.9% | 94.3% |
| Latvia | 92,552 | 64,364 | 28,188 | 66.4% | 69.2% | 58.0% | 83.7% | 85.8% | 66.4% | 55.5% | 2.1% | 10.9% | 68.5% | 62.6% | 95.9% |
| Togo | 72,892 | 57,348 | 15,544 | 35.9% | 49.8% | 42.3% | 84.9% | 12.4% | 45.2% | 38.6% | 3.5% | 6.6% | 48.7% | 77.5% | 95.4% |
| Croatia | 110,564 | 55,028 | 55,536 | 60.8% | 54.3% | 43.3% | 79.7% | 78.1% | 48.6% | 37.8% | 4.1% | 10.8% | 52.7% | 63.9% | 94.6% |
| Bosnia and Herzegovina | 51,832 | 51,820 | 12 | 57.2% | 69.8% | 57.1% | 81.9% | 100.0% | 68.7% | 56.5% | 0.6% | 12.2% | 69.3% | 74.0% | 94.4% |
| Costa Rica | 637,336 | 51,480 | 585,856 | 14.0% | 71.9% | 55.5% | 77.2% | 10.4% | 52.7% | 39.1% | 15.6% | 13.6% | 68.3% | 66.1% | 95.2% |
| Dominican Republic | 401,300 | 48,768 | 352,532 | 14.4% | 64.7% | 49.0% | 75.7% | 9.6% | 33.6% | 22.9% | 25.1% | 10.7% | 58.8% | 63.9% | 95.5% |
| Slovakia | 48,488 | 48,488 | - | 44.6% | 52.3% | 44.6% | 85.3% | | 52.1% | 44.5% | 0.1% | 7.6% | 52.2% | 81.2% | 96.0% |
| Estonia | 82,040 | 45,732 | 36,308 | 68.7% | 72.7% | 57.9% | 79.6% | 82.3% | 58.7% | 47.4% | 9.0% | 11.3% | 67.7% | 66.0% | 96.8% |
| Denmark | 2,670,696 | 42,528 | 2,628,168 | 31.3% | 21.7% | 20.7% | 95.2% | 31.5% | 6.4% | 5.4% | 6.7% | 1.0% | 13.1% | 65.2% | 97.7% |
| Switzerland | 41,392 | 41,392 | - | 56.8% | 71.7% | 56.8% | 79.2% | | 70.1% | 56.3% | 0.1% | 13.7% | 70.2% | 78.8% | 95.7% |
| Bhutan | 40,488 | 40,488 | - | 70.9% | 91.6% | 70.9% | 77.4% | | 91.6% | 70.9% | 0.0% | 20.7% | 91.6% | 74.9% | 97.5% |
| Netherlands | 182,376 | 37,300 | 145,076 | 40.2% | 46.5% | 45.1% | 97.0% | 39.0% | 43.1% | 36.9% | 2.2% | 6.2% | 45.3% | 67.6% | 97.1% |
| Taiwan | 467,724 | 36,420 | 431,304 | 21.9% | 62.4% | 39.7% | 63.6% | 20.4% | 23.3% | 9.6% | 29.0% | 13.8% | 52.3% | 65.1% | 97.3% |
| Moldova | 33,204 | 33,204 | - | 7.0% | 8.9% | 7.0% | 79.5% | | 8.0% | 6.4% | 0.5% | 1.5% | 8.4% | 71.4% | 94.7% |
| Guinea-Bissau | 140,552 | 33,048 | 107,504 | 29.1% | 73.4% | 56.8% | 77.3% | 20.5% | 38.4% | 25.5% | 29.9% | 12.9% | 68.3% | 68.7% | 96.8% |
| Belgium | 34,128 | 30,636 | 3,492 | 37.4% | 37.7% | 32.3% | 85.7% | 82.6% | 35.8% | 30.4% | 1.3% | 5.3% | 37.0% | 81.2% | 95.1% |
| Lesotho | 30,232 | 30,232 | - | 63.5% | 73.7% | 63.5% | 86.2% | | 4.1% | 4.0% | 59.6% | 0.1% | 63.7% | 84.3% | 98.5% |
| Armenia | 29,644 | 29,644 | - | 28.9% | 39.3% | 28.9% | 73.5% | | 35.9% | 28.6% | 0.1% | 7.3% | 36.0% | 61.4% | 95.6% |
| Albania | 40,520 | 28,352 | 12,168 | 49.2% | 46.7% | 38.6% | 82.6% | 74.0% | 41.1% | 33.7% | 4.2% | 7.4% | 45.3% | 74.7% | 94.3% |
| Solomon Islands | 1,643,080 | 27,356 | 1,615,724 | 48.2% | 91.7% | 49.5% | 54.0% | 48.2% | 83.0% | 42.2% | 1.9% | 40.7% | 84.8% | 63.6% | 97.1% |
| Burundi | 27,196 | 27,196 | - | 30.6% | 42.0% | 30.6% | 72.8% | | 32.5% | 28.4% | 1.7% | 4.1% | 34.3% | 78.5% | 97.1% |
| Haiti | 145,048 | 27,016 | 118,032 | 28.0% | 78.4% | 65.4% | 83.4% | 19.4% | 22.1% | 16.7% | 47.1% | 5.4% | 69.2% | 73.5% | 97.1% |
| Equatorial Guinea | 333,112 | 26,864 | 306,248 | 8.2% | 89.1% | 71.8% | 80.6% | 2.6% | 89.0% | 71.1% | 0.0% | 17.9% | 89.1% | 81.0% | 97.7% |
| Rwanda | 25,512 | 25,512 | - | 22.2% | 30.5% | 22.2% | 72.8% | | 23.4% | 19.7% | 2.3% | 3.7% | 25.8% | 75.3% | 94.1% |
| Macedonia | 25,380 | 25,380 | - | 46.9% | 59.6% | 46.9% | 78.8% | | 58.0% | 46.6% | 0.2% | 11.3% | 58.1% | 74.8% | 94.3% |
| Belize | 56,948 | 22,472 | 34,476 | 55.2% | 86.3% | 60.2% | 69.8% | 51.9% | 79.7% | 54.2% | 4.6% | 25.4% | 84.2% | 59.3% | 96.2% |
| Djibouti | 29,252 | 21,988 | 7,264 | 24.0% | 12.0% | 6.4% | 53.1% | 77.3% | 2.7% | 1.7% | 3.6% | 1.0% | 6.3% | 49.1% | 91.4% |
| Israel | 46,672 | 21,960 | 24,712 | 16.7% | 8.8% | 6.0% | 68.5% | 26.1% | 3.4% | 2.8% | 1.5% | 0.5% | 4.8% | 66.0% | 94.9% |
| El Salvador | 116,360 | 20,664 | 95,696 | 27.6% | 73.9% | 61.8% | 83.7% | 20.2% | 16.0% | 12.3% | 48.8% | 3.6% | 64.8% | 71.4% | 92.8% |
| Slovenia | 20,516 | 20,316 | 200 | 57.6% | 70.9% | 57.3% | 80.8% | 90.0% | 70.8% | 57.1% | 0.0% | 13.7% | 70.8% | 78.3% | 95.6% |

| | | | | | | | | | | | | | | | |
|---|---|---|---|---|---|---|---|---|---|---|---|---|---|---|---|
| New Caledonia | 18,956 | 18,956 | - | 38.3% | 75.9% | 38.3% | 50.4% | | 13.4% | 4.7% | 32.7% | 8.6% | 46.0% | 4.1% | 94.9% |
| Fiji | 1,300,116 | 18,944 | 1,281,172 | 11.3% | 64.2% | 60.8% | 94.8% | 10.5% | 16.3% | 13.5% | 35.8% | 2.8% | 52.0% | 58.8% | 97.3% |
| Kuwait | 28,772 | 17,536 | 11,236 | 28.9% | 1.2% | 1.2% | 100.0% | 72.2% | 0.0% | 0.0% | 0.2% | 0.0% | 0.2% | 43.6% | 94.5% |
| eSwatini | 17,196 | 17,196 | - | 52.8% | 61.6% | 52.8% | 85.8% | | 45.8% | 39.8% | 13.0% | 6.0% | 58.8% | 80.7% | 96.0% |
| East Timor | 93,164 | 15,184 | 77,980 | 20.9% | 15.7% | 12.0% | 76.2% | 22.7% | 12.7% | 8.2% | 2.3% | 4.5% | 15.0% | 66.3% | 95.6% |
| Montenegro | 20,144 | 13,764 | 6,380 | 61.5% | 81.4% | 64.5% | 79.2% | 55.2% | 80.6% | 63.5% | 0.2% | 17.1% | 80.8% | 75.0% | 93.2% |
| The Bahamas | 635,316 | 12,652 | 622,664 | 40.3% | 85.5% | 53.2% | 62.2% | 40.0% | 51.7% | 27.6% | 16.0% | 19.7% | 63.3% | 35.0% | 97.1% |
| Vanuatu | 639,236 | 12,332 | 626,904 | 19.1% | 86.2% | 48.4% | 56.1% | 18.5% | 65.4% | 30.8% | 11.1% | 34.3% | 76.2% | 48.3% | 94.6% |
| Falkland Islands | 11,488 | 11,488 | - | 6.3% | 0.0% | 0.0% | 0.0% | | 0.0% | 0.0% | 0.0% | 0.0% | 0.0% | 0.0% | 100.0% |
| Qatar | 42,848 | 11,220 | 31,628 | 32.7% | 1.8% | 1.8% | 100.0% | 43.5% | 0.0% | 0.0% | 0.3% | 0.0% | 0.3% | 29.2% | 98.5% |
| Jamaica | 284,904 | 11,056 | 273,848 | 16.1% | 78.3% | 63.7% | 81.4% | 14.1% | 43.2% | 32.8% | 28.9% | 10.4% | 72.1% | 66.0% | 94.8% |
| Kosovo | 10,928 | 10,928 | - | 35.9% | 48.8% | 35.9% | 73.6% | | 48.8% | 35.9% | 0.0% | 12.9% | 48.8% | 74.1% | 92.0% |
| Gambia | 33,912 | 10,672 | 23,240 | 28.2% | 32.6% | 26.9% | 82.7% | 28.8% | 11.5% | 8.0% | 17.2% | 3.4% | 28.7% | 78.4% | 94.4% |
| Lebanon | 30,268 | 10,044 | 20,224 | 27.4% | 32.4% | 28.7% | 88.6% | 26.7% | 4.4% | 3.8% | 16.8% | 0.6% | 21.2% | 60.4% | 90.4% |
| Puerto Rico | 9,052 | 9,052 | - | 69.9% | 72.2% | 69.9% | 96.8% | | 41.9% | 37.0% | 28.8% | 4.8% | 70.6% | 52.8% | 99.1% |
| French Southern and Antarctic Lan | 7,284 | 7,284 | - | 1.2% | 0.0% | 0.0% | 0.0% | | 0.0% | 0.0% | 0.0% | 0.0% | 0.0% | 0.0% | 100.0% |
| Palestine | 6,292 | 6,292 | - | 22.1% | 28.2% | 22.1% | 78.4% | | 14.0% | 11.2% | 4.6% | 2.8% | 18.6% | 74.7% | 96.0% |
| Brunei | 49,132 | 5,752 | 43,380 | 30.8% | 86.5% | 67.0% | 77.4% | 26.0% | 85.3% | 65.0% | 0.3% | 20.3% | 85.7% | 64.9% | 95.5% |
| Cyprus | 104,100 | 5,420 | 98,680 | 23.2% | 48.8% | 45.4% | 93.0% | 22.0% | 6.6% | 5.6% | 32.8% | 1.0% | 39.5% | 65.4% | 93.2% |
| Trinidad and Tobago | 82,164 | 5,144 | 77,020 | 34.8% | 73.3% | 58.0% | 79.2% | 33.3% | 69.4% | 50.9% | 3.4% | 18.6% | 72.9% | 68.8% | 96.4% |
| South Georgia and the Islands | 3,908 | 3,908 | - | 0.2% | 0.0% | 0.0% | 0.0% | | 0.0% | 0.0% | 0.0% | 0.0% | 0.0% | 0.0% | 100.0% |
| Cabo Verde | 810,336 | 3,888 | 806,448 | 62.8% | 0.0% | 0.0% | 0.0% | 63.1% | 0.0% | 0.0% | 0.0% | 0.0% | 0.0% | 5.0% | 100.0% |
| French Polynesia | 3,428 | 3,428 | - | 19.3% | 0.0% | 0.0% | 0.0% | | 0.0% | 0.0% | 0.0% | 0.0% | 0.0% | 30.3% | 89.1% |
| Northern Cyprus | 3,340 | 3,340 | - | 31.1% | 37.5% | 31.1% | 83.1% | | 1.2% | 0.8% | 13.4% | 0.4% | 14.6% | 56.9% | 93.8% |
| Samoa | 134,052 | 2,776 | 131,276 | 12.4% | 0.0% | 0.0% | 0.0% | 12.4% | 0.0% | 0.0% | 0.0% | 0.0% | 0.0% | 39.1% | 90.6% |
| Luxembourg | 2,620 | 2,620 | - | 49.9% | 57.1% | 49.9% | 87.4% | | 57.1% | 49.9% | 0.0% | 7.2% | 57.1% | 80.1% | 95.1% |
| Siachen Glacier | 2,064 | 2,064 | - | 0.0% | 10.1% | 0.0% | 0.0% | | 1.0% | 0.0% | 0.0% | 1.0% | 1.0% | #N/A | #N/A |
| Mauritius | 1,942,436 | 2,012 | 1,940,424 | 13.9% | 0.0% | 0.0% | 0.0% | 13.9% | 0.0% | 0.0% | 0.0% | 0.0% | 0.0% | 38.3% | 97.9% |
| Comoros | 167,140 | 1,640 | 165,500 | 10.3% | 0.0% | 0.0% | 0.0% | 10.2% | 0.0% | 0.0% | 0.0% | 0.0% | 0.0% | 35.2% | 100.0% |
| Faroe Islands | 1,328 | 1,328 | - | 35.8% | 74.1% | 35.8% | 48.4% | | 0.0% | 0.0% | 3.3% | 0.0% | 3.3% | 5.0% | 97.5% |
| Hong Kong S.A.R. | 1,048 | 1,048 | - | 51.5% | 51.5% | 51.5% | 100.0% | | 5.0% | 2.7% | 35.5% | 2.3% | 40.5% | 17.0% | 94.1% |
| São Tomé and Príncipe | 167,496 | 1,032 | 166,464 | 28.0% | 0.0% | 0.0% | 0.0% | 28.1% | 0.0% | 0.0% | 0.0% | 0.0% | 0.0% | 0.0% | 100.0% |
| Kiribati | 3,462,532 | 988 | 3,461,544 | 38.9% | 0.0% | 0.0% | 0.0% | 38.9% | 0.0% | 0.0% | 0.0% | 0.0% | 0.0% | 23.3% | 99.2% |
| Aland | 920 | 920 | - | 44.8% | 73.9% | 44.8% | 60.6% | | 8.7% | 5.7% | 31.3% | 3.0% | 40.0% | 42.7% | 94.2% |
| Dominica | 29,480 | 744 | 28,736 | 29.1% | 94.6% | 87.6% | 92.6% | 27.6% | 91.9% | 81.7% | 2.7% | 10.2% | 94.6% | 55.8% | 86.5% |
| Federated States of Micronesia | 3,030,868 | 624 | 3,030,244 | 72.4% | 0.0% | 0.0% | 0.0% | 72.4% | 0.0% | 0.0% | 0.0% | 0.0% | 0.0% | 13.0% | 95.7% |
| Saint Lucia | 16,124 | 624 | 15,500 | 46.1% | 84.6% | 69.2% | 81.8% | 45.2% | 82.1% | 60.3% | 2.6% | 21.8% | 84.6% | 56.5% | 87.0% |
| Bahrain | 8,140 | 596 | 7,544 | 76.1% | 5.4% | 5.4% | 100.0% | 80.4% | 0.7% | 0.7% | 0.7% | 0.0% | 1.3% | 56.3% | 81.3% |
| Tonga | 670,056 | 592 | 669,464 | 8.2% | 0.0% | 0.0% | 0.0% | 8.2% | 0.0% | 0.0% | 0.0% | 0.0% | 0.0% | 31.6% | 96.5% |
| Isle of Man | 580 | 580 | - | 46.2% | 79.3% | 46.2% | 58.3% | | 77.9% | 42.8% | 1.4% | 35.2% | 79.3% | 76.1% | 91.0% |
| Northern Mariana Islands | 580 | 580 | - | 21.4% | 0.0% | 0.0% | 0.0% | | 0.0% | 0.0% | 0.0% | 0.0% | 0.0% | 32.3% | 96.8% |
| Guam | 560 | 560 | - | 30.7% | 0.0% | 0.0% | 0.0% | | 0.0% | 0.0% | 0.0% | 0.0% | 0.0% | 27.9% | 100.0% |
| Singapore | 1,216 | 520 | 696 | 59.2% | 13.1% | 13.1% | 100.0% | 90.2% | 9.2% | 8.5% | 3.8% | 0.8% | 13.1% | 47.8% | 100.0% |
| Palau | 619,300 | 500 | 618,800 | 63.2% | 0.0% | 0.0% | 0.0% | 63.2% | 0.0% | 0.0% | 0.0% | 0.0% | 0.0% | 43.6% | 100.0% |
| Antigua and Barbuda | 112,620 | 464 | 112,156 | 13.4% | 78.4% | 78.4% | 100.0% | 13.1% | 18.1% | 10.3% | 55.2% | 7.8% | 73.3% | 69.2% | 94.5% |
| Curaçao | 456 | 456 | - | 50.9% | 78.1% | 50.9% | 65.2% | | 5.3% | 3.5% | 36.8% | 1.8% | 42.1% | 13.8% | 91.4% |
| Andorra | 448 | 448 | - | 82.1% | 93.8% | 82.1% | 87.6% | | 93.8% | 82.1% | 0.0% | 11.6% | 93.8% | 71.7% | 92.4% |
| Seychelles | 1,350,692 | 436 | 1,350,256 | 19.6% | 0.0% | 0.0% | 0.0% | 19.6% | 0.0% | 0.0% | 0.0% | 0.0% | 0.0% | 21.4% | 97.1% |
| Turks and Caicos Islands | 432 | 432 | - | 63.9% | 75.9% | 63.9% | 84.1% | | 43.5% | 29.6% | 25.0% | 12.0% | 66.7% | 2.9% | 98.6% |
| Barbados | 186,556 | 428 | 186,128 | 7.5% | 13.1% | 13.1% | 100.0% | 7.5% | 8.4% | 7.5% | 4.7% | 0.9% | 13.1% | 58.8% | 82.4% |
| Heard Island and McDonald Island: | 392 | 392 | - | 11.2% | 0.0% | 0.0% | 0.0% | | 0.0% | 0.0% | 0.0% | 0.0% | 0.0% | 0.0% | 100.0% |
| Saint Helena | 368 | 368 | - | 0.0% | 0.0% | 0.0% | 0.0% | | 0.0% | 0.0% | 0.0% | 0.0% | 0.0% | #N/A | #N/A |
| United States Virgin Islands | 360 | 360 | - | 76.7% | 58.9% | 58.9% | 100.0% | | 32.2% | 27.8% | 23.3% | 4.4% | 55.6% | 33.3% | 98.6% |
| Grenada | 26,068 | 348 | 25,720 | 43.7% | 90.8% | 78.2% | 86.1% | 43.3% | 89.7% | 69.0% | 1.1% | 20.7% | 90.8% | 52.9% | 86.8% |
| Saint Vincent and the Grenadines | 36,832 | 348 | 36,484 | 38.7% | 81.6% | 59.8% | 73.2% | 38.5% | 80.5% | 50.6% | 1.1% | 29.9% | 81.6% | 44.2% | 92.3% |

| | | | | | | | | | | | | | | | |
|---|---|---|---|---|---|---|---|---|---|---|---|---|---|---|---|
| Cyprus No Mans Area | 324 | 324 | - | 32.1% | 37.0% | 32.1% | 86.7% | | 0.0% | 0.0% | 14.8% | 0.0% | 14.8% | 80.8% | 100.0% |
| Malta | 53,360 | 324 | 53,036 | 52.2% | 2.5% | 2.5% | 100.0% | 52.4% | 2.5% | 2.5% | 0.0% | 0.0% | 2.5% | 9.1% | 100.0% |
| Cayman Islands | 312 | 312 | - | 87.2% | 70.5% | 70.5% | 100.0% | | 56.4% | 51.3% | 14.1% | 5.1% | 70.5% | 33.8% | 100.0% |
| Saint Kitts and Nevis | 9,844 | 272 | 9,572 | 44.5% | 69.1% | 69.1% | 100.0% | 43.3% | 26.5% | 25.0% | 36.8% | 1.5% | 63.2% | 52.5% | 84.7% |
| Saint Pierre and Miquelon | 224 | 224 | - | 17.9% | 85.7% | 17.9% | 20.8% | | 21.4% | 0.0% | 10.7% | 19.6% | 30.4% | 30.0% | 100.0% |
| Niue | 216 | 216 | - | 16.7% | 0.0% | 0.0% | 0.0% | | 0.0% | 0.0% | 0.0% | 0.0% | 0.0% | 0.0% | 66.7% |
| Cook Islands | 188 | 188 | - | 23.4% | 0.0% | 0.0% | 0.0% | | 0.0% | 0.0% | 0.0% | 0.0% | 0.0% | 27.3% | 100.0% |
| Aruba | 172 | 172 | - | 60.5% | 74.4% | 60.5% | 81.3% | | 0.0% | 0.0% | 46.5% | 0.0% | 46.5% | 42.3% | 84.6% |
| American Samoa | 164 | 164 | - | 24.4% | 0.0% | 0.0% | 0.0% | | 0.0% | 0.0% | 0.0% | 0.0% | 0.0% | 0.0% | 70.0% |
| British Virgin Islands | 152 | 152 | - | 86.8% | 73.7% | 73.7% | 100.0% | | 52.6% | 47.4% | 15.8% | 5.3% | 68.4% | 48.5% | 100.0% |
| Liechtenstein | 148 | 148 | - | 56.8% | 64.9% | 56.8% | 87.5% | | 64.9% | 56.8% | 0.0% | 8.1% | 64.9% | 52.4% | 90.5% |
| Marshall Islands | 2,014,328 | 140 | 2,014,188 | 71.1% | 0.0% | 0.0% | 0.0% | 71.1% | 0.0% | 0.0% | 0.0% | 0.0% | 0.0% | 15.2% | 100.0% |
| Wallis and Futuna | 136 | 136 | - | 55.9% | 0.0% | 0.0% | 0.0% | | 0.0% | 0.0% | 0.0% | 0.0% | 0.0% | 15.8% | 100.0% |
| Jersey | 128 | 128 | - | 21.9% | 12.5% | 12.5% | 100.0% | | 3.1% | 0.0% | 3.1% | 3.1% | 6.3% | 0.0% | 85.7% |
| Dhekelia Sovereign Base Area | 120 | 120 | - | 16.7% | 6.7% | 6.7% | 100.0% | | 0.0% | 0.0% | 0.0% | 0.0% | 0.0% | 20.0% | 100.0% |
| Indian Ocean Territories | 104 | 104 | - | 42.3% | 0.0% | 0.0% | 0.0% | | 0.0% | 0.0% | 0.0% | 0.0% | 0.0% | 0.0% | 100.0% |
| Akrotiri Sovereign Base Area | 104 | 104 | - | 23.1% | 19.2% | 19.2% | 100.0% | | 11.5% | 7.7% | 3.8% | 3.8% | 15.4% | 0.0% | 66.7% |
| Montserrat | 100 | 100 | - | 60.0% | 84.0% | 60.0% | 71.4% | | 60.0% | 28.0% | 24.0% | 32.0% | 84.0% | 20.0% | 86.7% |
| Maldives | 926,852 | 96 | 926,756 | 11.3% | 0.0% | 0.0% | 0.0% | 11.3% | 0.0% | 0.0% | 0.0% | 0.0% | 0.0% | 80.0% | 100.0% |
| Anguilla | 76 | 76 | - | 73.7% | 78.9% | 73.7% | 93.3% | | 0.0% | 0.0% | 57.9% | 0.0% | 57.9% | 35.7% | 100.0% |
| Guernsey | 72 | 72 | - | 44.4% | 5.6% | 5.6% | 100.0% | | 0.0% | 0.0% | 0.0% | 0.0% | 0.0% | 12.5% | 100.0% |
| Saint Martin | 72 | 72 | - | 94.4% | 66.7% | 66.7% | 100.0% | | 16.7% | 16.7% | 44.4% | 0.0% | 61.1% | 29.4% | 100.0% |
| Bermuda | 68 | 68 | - | 70.6% | 0.0% | 0.0% | 0.0% | | 0.0% | 0.0% | 0.0% | 0.0% | 0.0% | 58.3% | 100.0% |
| San Marino | 64 | 64 | - | 0.0% | 0.0% | 0.0% | 0.0% | | 0.0% | 0.0% | 0.0% | 0.0% | 0.0% | | |
| US Naval Base Guantanamo Bay | 64 | 64 | - | 37.5% | 93.8% | 37.5% | 40.0% | | 87.5% | 25.0% | 6.3% | 62.5% | 93.8% | 50.0% | 100.0% |
| Pitcairn Islands | 44 | 44 | - | 0.0% | 0.0% | 0.0% | 0.0% | | 0.0% | 0.0% | 0.0% | 0.0% | 0.0% | | |
| Macao S.A.R | 40 | 40 | - | 50.0% | 30.0% | 30.0% | 100.0% | | 20.0% | 20.0% | 0.0% | 0.0% | 20.0% | 20.0% | 100.0% |
| British Indian Ocean Territory | 36 | 36 | - | 44.4% | 0.0% | 0.0% | 0.0% | | 0.0% | 0.0% | 0.0% | 0.0% | 0.0% | 0.0% | 100.0% |
| Norfolk Island | 32 | 32 | - | 12.5% | 12.5% | 12.5% | 100.0% | | 0.0% | 0.0% | 0.0% | 0.0% | 0.0% | 0.0% | 100.0% |
| Nauru | 311,364 | 32 | 311,332 | 71.4% | 0.0% | 0.0% | 0.0% | 71.4% | 0.0% | 0.0% | 0.0% | 0.0% | 0.0% | 33.3% | 100.0% |
| United States Minor Outlying Islanc | 32 | 32 | - | 0.0% | 25.0% | 0.0% | 0.0% | | 0.0% | 0.0% | 0.0% | 0.0% | 0.0% | | |
| Saint Barthelemy | 28 | 28 | - | 100.0% | 42.9% | 42.9% | 100.0% | | 0.0% | 0.0% | 42.9% | 0.0% | 42.9% | 85.7% | 100.0% |
| Sint Maarten | 24 | 24 | - | 83.3% | 66.7% | 66.7% | 100.0% | | 16.7% | 16.7% | 50.0% | 0.0% | 66.7% | 40.0% | 100.0% |
| Tuvalu | 726,192 | 20 | 726,172 | 66.1% | 0.0% | 0.0% | 0.0% | 66.1% | 0.0% | 0.0% | 0.0% | 0.0% | 0.0% | 0.0% | 100.0% |
| Monaco | 300 | 16 | 284 | 33.3% | 0.0% | 0.0% | 0.0% | 32.4% | 0.0% | 0.0% | 0.0% | 0.0% | 0.0% | 0.0% | 100.0% |
| Spratly Islands | 12 | 12 | - | 100.0% | 0.0% | 0.0% | 0.0% | | 0.0% | 0.0% | 0.0% | 0.0% | 0.0% | 0.0% | 100.0% |
| Scarborough Reef | 8 | 8 | - | 100.0% | 0.0% | 0.0% | 0.0% | | 0.0% | 0.0% | 0.0% | 0.0% | 0.0% | 0.0% | 100.0% |
| Serranilla Bank | 8 | 8 | - | 100.0% | 0.0% | 0.0% | 0.0% | | 0.0% | 0.0% | 0.0% | 0.0% | 0.0% | 0.0% | 100.0% |
| Bajo Nuevo Bank (Petrel Is.) | 4 | 4 | - | 100.0% | 0.0% | 0.0% | 0.0% | | 0.0% | 0.0% | 0.0% | 0.0% | 0.0% | 0.0% | 100.0% |
| Clipperton Island | 4 | 4 | - | 0.0% | 0.0% | 0.0% | 0.0% | | 0.0% | 0.0% | 0.0% | 0.0% | 0.0% | | |
| Coral Sea Islands | 4 | 4 | - | 0.0% | 0.0% | 0.0% | 0.0% | | 0.0% | 0.0% | 0.0% | 0.0% | 0.0% | | |
| Gibraltar | 388 | 4 | 384 | 100.0% | 0.0% | 0.0% | 0.0% | 100.0% | 0.0% | 0.0% | 0.0% | 0.0% | 0.0% | 100.0% | 100.0% |
| Vatican | 4 | 4 | - | 0.0% | 0.0% | 0.0% | 0.0% | | 0.0% | 0.0% | 0.0% | 0.0% | 0.0% | | |

### SI Table 2. Human population benefitting from and living on critical natural assets.

| Country (ranked by population) | Total population (Landscan 2017) | Population on land that is critical natural assets for local NCP (LCNA) | Population on land critical for global NCP (GCNA) | Percent of population living on LCNA | Percent of population living on GCNA | Ratio of % pop to % land for LCNA | Ratio of % pop to % land for GCNA | Population benefitting from LCNA | Population benefitting from GCNA | Percent of population benefitting from LCNA | Percent of population benefitting from GCNA | Ratio of pop benefiting to pop living on LCNA | Ratio of pop benefiting to pop living on GNCA |
|---|---|---|---|---|---|---|---|---|---|---|---|---|---|
| Global | 7,341,941,858 | 1,168,934,232 | 771,148,868 | 15.9% | 10.5% | 0.523 | 0.273 | 6,389,108,220 | 5,369,617,498 | 87.0% | 73.1% | 5.466 | 6.963 |
| China | 1,370,730,445 | 136,967,856 | 82,980,379 | 10.0% | 6.1% | 0.394 | 0.159 | 1,121,101,594 | 890,449,284 | 81.8% | 10.0% | 8.185 | 1.651 |
| India | 1,278,133,079 | 88,485,930 | 50,633,003 | 6.9% | 4.0% | 0.357 | 0.199 | 921,797,950 | 655,768,365 | 72.1% | 6.9% | 10.417 | 1.748 |
| United States of America | 323,806,117 | 92,689,394 | 80,664,054 | 28.6% | 24.9% | 0.766 | 0.471 | 320,403,145 | 311,306,286 | 98.9% | 28.6% | 3.457 | 1.149 |
| Indonesia | 254,421,809 | 37,878,004 | 16,280,107 | 14.9% | 6.4% | 0.452 | 0.110 | 244,939,654 | 230,594,870 | 96.3% | 14.9% | 6.467 | 2.327 |
| Brazil | 204,578,040 | 53,052,869 | 35,002,397 | 25.9% | 17.1% | 0.564 | 0.243 | 201,914,446 | 199,952,596 | 98.7% | 25.9% | 3.806 | 1.516 |
| Pakistan | 204,166,120 | 16,198,897 | 10,576,391 | 7.9% | 5.2% | 0.383 | 0.319 | 155,734,897 | 115,600,400 | 76.3% | 7.9% | 9.614 | 1.532 |
| Nigeria | 188,649,700 | 30,174,379 | 27,748,061 | 16.0% | 14.7% | 0.592 | 0.471 | 153,898,199 | 126,478,041 | 81.6% | 16.0% | 5.100 | 1.087 |
| Bangladesh | 157,117,529 | 13,698,772 | 5,998,490 | 8.7% | 3.8% | 0.490 | 0.240 | 137,103,359 | 111,984,561 | 87.3% | 8.7% | 10.008 | 2.284 |
| Russia | 144,144,677 | 28,516,962 | 23,774,810 | 19.8% | 16.5% | 1.110 | 0.461 | 138,137,303 | 136,757,367 | 95.8% | 19.8% | 4.844 | 1.199 |
| Japan | 125,239,921 | 15,017,870 | 3,528,044 | 12.0% | 2.8% | 0.248 | 0.107 | 123,941,148 | 119,894,505 | 99.0% | 12.0% | 8.253 | 4.257 |
| Mexico | 123,993,577 | 25,401,136 | 9,870,496 | 20.5% | 8.0% | 0.417 | 0.241 | 122,195,434 | 93,782,223 | 98.5% | 20.5% | 4.811 | 2.573 |
| Ethiopia | 105,206,039 | 32,855,419 | 33,405,019 | 31.2% | 31.8% | 0.661 | 0.516 | 90,897,374 | 90,311,138 | 86.4% | 31.2% | 2.767 | 0.984 |
| Egypt | 98,727,615 | 3,812,238 | 145,998 | 3.9% | 0.1% | 5.435 | 1.698 | 95,344,349 | 44,418,846 | 96.6% | 3.9% | 25.010 | 26.112 |
| Philippines | 97,898,021 | 15,988,760 | 3,096,444 | 16.3% | 3.2% | 0.600 | 0.124 | 93,484,786 | 82,187,345 | 95.5% | 16.3% | 5.847 | 5.164 |
| Vietnam | 95,036,501 | 11,495,970 | 5,867,111 | 12.1% | 6.2% | 0.311 | 0.145 | 92,859,430 | 84,897,422 | 97.7% | 12.1% | 8.078 | 1.959 |
| Democratic Republic of the Congo | 82,863,685 | 29,354,443 | 30,967,493 | 35.4% | 37.4% | 0.502 | 0.431 | 74,755,318 | 76,216,225 | 90.2% | 35.4% | 2.547 | 0.948 |
| Iran | 81,452,008 | 4,310,973 | 3,224,677 | 5.3% | 4.0% | 0.601 | 0.358 | 71,015,774 | 61,520,235 | 87.2% | 5.3% | 16.473 | 1.337 |
| Germany | 80,593,192 | 16,166,956 | 16,508,367 | 20.1% | 20.5% | 0.523 | 0.442 | 80,544,646 | 80,502,951 | 99.9% | 20.1% | 4.982 | 0.979 |
| Turkey | 80,143,741 | 12,441,988 | 8,512,473 | 15.5% | 10.6% | 0.506 | 0.292 | 76,682,318 | 75,576,769 | 95.7% | 15.5% | 6.163 | 1.462 |
| Thailand | 68,142,105 | 4,238,765 | 2,238,970 | 6.2% | 3.3% | 0.277 | 0.111 | 66,102,827 | 61,558,169 | 97.0% | 6.2% | 15.595 | 1.893 |
| France | 65,735,954 | 12,515,094 | 9,930,964 | 19.0% | 15.1% | 0.566 | 0.328 | 65,331,520 | 63,169,365 | 99.4% | 19.0% | 5.220 | 1.260 |
| United Kingdom | 64,098,136 | 15,403,218 | 13,682,601 | 24.0% | 21.3% | 0.501 | 0.413 | 63,471,562 | 62,736,916 | 99.0% | 24.0% | 4.121 | 1.126 |
| Italy | 61,240,480 | 7,929,664 | 5,085,296 | 12.9% | 8.3% | 0.408 | 0.252 | 60,132,414 | 56,320,521 | 98.2% | 12.9% | 7.583 | 1.559 |
| South Africa | 54,657,349 | 19,895,337 | 5,092,550 | 36.4% | 9.3% | 0.614 | 0.735 | 53,838,279 | 41,179,075 | 98.5% | 36.4% | 2.706 | 3.907 |
| United Republic of Tanzania | 53,633,880 | 16,774,742 | 15,310,375 | 31.3% | 28.5% | 0.684 | 0.452 | 49,533,541 | 46,719,072 | 92.4% | 31.3% | 2.953 | 1.096 |
| Myanmar | 53,484,510 | 6,819,231 | 5,226,354 | 12.7% | 9.8% | 0.262 | 0.152 | 42,544,083 | 35,522,508 | 79.5% | 12.7% | 6.239 | 1.305 |
| South Korea | 50,315,468 | 9,238,749 | 672,164 | 18.4% | 1.3% | 0.468 | 0.122 | 49,901,414 | 46,863,091 | 99.2% | 18.4% | 5.401 | 13.745 |
| Spain | 48,031,946 | 9,769,735 | 4,413,926 | 20.3% | 9.2% | 0.573 | 0.287 | 46,945,353 | 42,382,637 | 97.7% | 20.3% | 4.805 | 2.213 |
| Colombia | 47,355,574 | 13,046,700 | 12,495,809 | 27.6% | 26.4% | 0.575 | 0.308 | 46,594,767 | 45,802,441 | 98.4% | 27.6% | 3.571 | 1.044 |
| Kenya | 47,102,899 | 12,165,359 | 11,551,089 | 25.8% | 24.5% | 0.536 | 0.373 | 42,446,651 | 41,731,590 | 90.1% | 25.8% | 3.489 | 1.053 |
| Argentina | 44,284,392 | 7,302,668 | 3,327,867 | 16.5% | 7.5% | 0.373 | 0.326 | 42,807,246 | 37,050,399 | 96.7% | 16.5% | 5.862 | 2.194 |
| Ukraine | 41,708,795 | 3,363,002 | 2,780,221 | 8.1% | 6.7% | 0.495 | 0.370 | 38,750,223 | 38,058,925 | 92.9% | 8.1% | 11.523 | 1.210 |
| Algeria | 40,728,545 | 7,037,439 | 2,645,410 | 17.3% | 6.5% | 6.125 | 2.130 | 35,804,305 | 24,464,776 | 87.9% | 17.3% | 5.088 | 2.660 |
| Iraq | 40,335,504 | 969,294 | 215,113 | 2.4% | 0.5% | 1.566 | 0.466 | 31,931,815 | 23,861,691 | 79.2% | 2.4% | 32.943 | 4.506 |
| Uganda | 39,730,276 | 4,586,510 | 4,562,523 | 11.5% | 11.5% | 0.362 | 0.317 | 33,962,261 | 32,914,413 | 85.5% | 11.5% | 7.405 | 1.005 |
| Poland | 38,548,752 | 5,969,387 | 6,095,178 | 15.5% | 15.8% | 0.480 | 0.412 | 38,536,972 | 38,539,364 | 100.0% | 15.5% | 6.456 | 0.979 |
| Sudan | 37,353,956 | 10,342,147 | 5,037,275 | 27.7% | 13.5% | 1.492 | 0.928 | 32,026,917 | 20,814,860 | 85.7% | 27.7% | 3.097 | 2.053 |
| Canada | 34,476,073 | 7,195,593 | 4,390,118 | 20.9% | 12.7% | 1.952 | 0.459 | 33,896,058 | 32,737,887 | 98.3% | 20.9% | 4.711 | 1.639 |
| Afghanistan | 34,095,851 | 8,937,048 | 9,619,524 | 26.2% | 28.2% | 0.948 | 0.616 | 31,892,732 | 32,587,792 | 93.5% | 26.2% | 3.569 | 0.929 |
| Morocco | 34,044,087 | 6,483,979 | 1,504,189 | 19.0% | 4.4% | 1.189 | 0.616 | 31,796,604 | 22,334,090 | 93.4% | 19.0% | 4.904 | 4.311 |
| Peru | 30,978,783 | 11,105,547 | 6,537,870 | 35.8% | 21.1% | 0.627 | 0.280 | 30,135,549 | 18,088,017 | 97.3% | 35.8% | 2.714 | 1.699 |
| Venezuela | 30,880,169 | 9,131,402 | 5,173,372 | 29.6% | 16.8% | 0.491 | 0.210 | 30,229,025 | 29,112,431 | 97.9% | 29.6% | 3.310 | 1.765 |

| | | | | | | | | | | | | | |
|---|---|---|---|---|---|---|---|---|---|---|---|---|---|
| Nepal | 30,593,400 | 6,987,071 | 7,053,216 | 22.8% | 23.1% | 0.442 | 0.354 | 27,279,292 | 27,225,436 | 89.2% | 22.8% | 3.904 | 0.991 |
| Malaysia | 30,011,529 | 3,983,654 | 1,755,630 | 13.3% | 5.8% | 0.298 | 0.104 | 29,287,853 | 27,658,771 | 97.6% | 13.3% | 7.352 | 2.269 |
| Uzbekistan | 29,799,358 | 946,323 | 776,507 | 3.2% | 2.6% | 0.281 | 0.191 | 25,466,443 | 22,242,395 | 85.5% | 3.2% | 26.911 | 1.219 |
| Angola | 29,183,564 | 14,700,041 | 12,895,337 | 50.4% | 44.2% | 0.765 | 0.494 | 28,295,356 | 27,161,838 | 97.0% | 50.4% | 1.925 | 1.140 |
| Saudi Arabia | 28,124,932 | 1,956,729 | 1,684 | 7.0% | 0.0% | 3.202 | 0.466 | 16,732,351 | 137,639 | 59.5% | 7.0% | 8.551 | 1,161.953 |
| Yemen | 27,895,190 | 8,108,278 | 25,988 | 29.1% | 0.1% | 2.928 | 3.659 | 20,286,191 | 1,025,233 | 72.7% | 29.1% | 2.502 | 312.001 |
| Ghana | 27,287,268 | 2,640,184 | 1,829,416 | 9.7% | 6.7% | 0.225 | 0.153 | 20,374,440 | 16,544,090 | 74.7% | 9.7% | 7.717 | 1.443 |
| Mozambique | 26,560,095 | 12,582,904 | 10,107,029 | 47.4% | 38.1% | 0.766 | 0.522 | 25,762,102 | 22,104,895 | 97.0% | 47.4% | 2.047 | 1.245 |
| Cameroon | 24,953,119 | 7,287,435 | 7,467,623 | 29.2% | 29.9% | 0.468 | 0.360 | 22,841,594 | 22,567,703 | 91.5% | 29.2% | 3.134 | 0.976 |
| Madagascar | 24,819,901 | 13,785,553 | 10,547,279 | 55.5% | 42.5% | 0.854 | 0.659 | 23,783,289 | 21,089,185 | 95.8% | 55.5% | 1.725 | 1.307 |
| Ivory Coast | 23,599,114 | 2,953,176 | 2,870,149 | 12.5% | 12.2% | 0.335 | 0.260 | 14,656,774 | 14,139,262 | 62.1% | 12.5% | 4.963 | 1.029 |
| North Korea | 23,401,940 | 4,246,197 | 682,155 | 18.1% | 2.9% | 0.372 | 0.113 | 22,773,176 | 14,234,123 | 97.3% | 18.1% | 5.363 | 6.225 |
| Taiwan | 23,085,648 | 2,030,945 | 358,142 | 8.8% | 1.6% | 0.222 | 0.066 | 23,015,392 | 22,824,713 | 99.7% | 8.8% | 11.332 | 5.671 |
| Australia | 22,849,333 | 5,705,191 | 2,576,895 | 25.0% | 11.3% | 1.024 | 1.077 | 22,417,286 | 21,405,879 | 98.1% | 25.0% | 3.929 | 2.214 |
| Sri Lanka | 22,137,848 | 7,376,273 | 3,908,519 | 33.3% | 17.7% | 0.844 | 0.544 | 21,979,776 | 21,675,401 | 99.3% | 33.3% | 2.980 | 1.887 |
| Romania | 21,459,263 | 2,884,618 | 2,899,927 | 13.4% | 13.5% | 0.382 | 0.330 | 20,730,683 | 20,541,668 | 96.6% | 13.4% | 7.187 | 0.995 |
| Burkina Faso | 20,160,229 | 2,650,836 | 387,533 | 13.1% | 1.9% | 0.582 | 0.337 | 12,726,056 | 3,793,231 | 63.1% | 13.1% | 4.801 | 6.840 |
| Niger | 19,236,051 | 7,093,159 | 7,744 | 36.9% | 0.0% | 2.029 | 0.315 | 17,585,819 | 127,116 | 91.4% | 36.9% | 2.479 | 915.955 |
| Malawi | 19,172,271 | 2,911,717 | 1,752,459 | 15.2% | 9.1% | 0.605 | 0.528 | 18,365,152 | 15,909,947 | 95.8% | 15.2% | 6.307 | 1.662 |
| Kazakhstan | 18,469,884 | 3,547,563 | 3,334,665 | 19.2% | 18.1% | 0.546 | 0.327 | 17,190,748 | 17,049,572 | 93.1% | 19.2% | 4.846 | 1.064 |
| Syria | 18,078,901 | 952,754 | 562,306 | 5.3% | 3.1% | 1.371 | 1.114 | 15,833,615 | 13,227,577 | 87.6% | 5.3% | 16.619 | 1.694 |
| Mali | 17,924,669 | 3,901,798 | 567,724 | 21.8% | 3.2% | 1.174 | 0.768 | 14,235,256 | 6,603,424 | 79.4% | 21.8% | 3.648 | 6.873 |
| Chile | 17,538,346 | 5,804,057 | 1,570,091 | 33.1% | 9.0% | 1.336 | 0.244 | 17,250,202 | 15,478,976 | 98.4% | 33.1% | 2.972 | 3.697 |
| Netherlands | 16,952,999 | 4,249,024 | 3,322,790 | 25.1% | 19.6% | 0.556 | 0.455 | 16,906,288 | 16,805,413 | 99.7% | 25.1% | 3.979 | 1.279 |
| Cambodia | 16,167,046 | 1,383,540 | 753,404 | 8.6% | 4.7% | 0.205 | 0.103 | 14,977,765 | 13,546,661 | 92.6% | 8.6% | 10.826 | 1.836 |
| Zambia | 15,901,762 | 6,702,292 | 7,063,437 | 42.1% | 44.4% | 0.617 | 0.543 | 15,594,987 | 15,664,511 | 98.1% | 42.1% | 2.327 | 0.949 |
| Ecuador | 15,625,817 | 5,359,966 | 4,152,070 | 34.3% | 26.6% | 0.694 | 0.348 | 14,964,415 | 14,613,910 | 95.8% | 34.3% | 2.792 | 1.291 |
| Guatemala | 15,469,543 | 8,253,278 | 5,169,708 | 53.4% | 33.4% | 1.082 | 0.600 | 15,274,990 | 14,792,750 | 98.7% | 53.4% | 1.851 | 1.596 |
| Senegal | 14,495,435 | 2,364,104 | 125,418 | 16.3% | 0.9% | 0.307 | 0.203 | 6,408,641 | 624,241 | 44.2% | 16.3% | 2.711 | 18.850 |
| Zimbabwe | 13,748,107 | 4,199,488 | 4,679,260 | 30.5% | 34.0% | 0.616 | 0.555 | 13,206,804 | 13,254,149 | 96.1% | 30.5% | 3.145 | 0.897 |
| South Sudan | 12,974,180 | 8,276,157 | 8,894,975 | 63.8% | 68.6% | 0.807 | 0.725 | 12,676,631 | 12,943,655 | 97.7% | 63.8% | 1.532 | 0.930 |
| Guinea | 12,340,694 | 4,929,610 | 3,729,441 | 39.9% | 30.2% | 0.630 | 0.479 | 12,041,815 | 10,773,549 | 97.6% | 39.9% | 2.443 | 1.322 |
| Chad | 12,175,771 | 3,588,821 | 1,356,201 | 29.5% | 11.1% | 1.373 | 0.768 | 10,154,030 | 6,022,361 | 83.4% | 29.5% | 2.829 | 2.646 |
| Rwanda | 11,963,961 | 1,591,662 | 1,410,268 | 13.3% | 11.8% | 0.600 | 0.503 | 4,477,667 | 3,514,880 | 37.4% | 13.3% | 2.813 | 1.129 |
| Belgium | 11,537,984 | 1,593,287 | 1,459,625 | 13.8% | 12.7% | 0.428 | 0.354 | 11,537,502 | 11,537,381 | 100.0% | 13.8% | 7.241 | 1.092 |
| Burundi | 11,504,302 | 2,452,579 | 2,399,609 | 21.3% | 20.9% | 0.696 | 0.641 | 11,163,503 | 10,901,545 | 97.0% | 21.3% | 4.552 | 1.022 |
| Tunisia | 11,222,162 | 946,281 | 157,609 | 8.4% | 1.4% | 1.046 | 0.405 | 10,372,409 | 3,152,730 | 92.4% | 8.4% | 10.961 | 6.004 |
| Benin | 11,107,123 | 1,545,576 | 1,275,130 | 13.9% | 11.5% | 0.305 | 0.231 | 9,512,825 | 8,837,572 | 85.6% | 13.9% | 6.155 | 1.212 |
| Bolivia | 11,032,684 | 3,409,936 | 3,677,689 | 30.9% | 33.3% | 0.511 | 0.402 | 10,688,546 | 10,946,719 | 96.9% | 30.9% | 3.135 | 0.927 |
| Cuba | 10,983,207 | 2,278,618 | 325,491 | 20.7% | 3.0% | 0.541 | 0.163 | 10,941,840 | 8,911,485 | 99.6% | 20.7% | 4.802 | 7.001 |
| Czechia | 10,627,182 | 1,852,064 | 2,031,222 | 17.4% | 19.1% | 0.447 | 0.421 | 10,627,182 | 10,627,182 | 100.0% | 17.4% | 5.738 | 0.912 |
| Portugal | 10,535,151 | 2,151,436 | 551,744 | 20.4% | 5.2% | 0.549 | 0.212 | 10,302,456 | 7,996,576 | 97.8% | 20.4% | 4.789 | 3.899 |
| Dominican Republic | 10,455,716 | 2,478,040 | 564,098 | 23.7% | 5.4% | 0.484 | 0.160 | 10,064,962 | 8,879,523 | 96.3% | 23.7% | 4.062 | 4.393 |
| Greece | 10,399,857 | 1,762,910 | 724,722 | 17.0% | 7.0% | 0.380 | 0.167 | 10,290,822 | 9,575,140 | 99.0% | 17.0% | 5.837 | 2.433 |
| Haiti | 10,207,934 | 5,364,900 | 1,460,629 | 52.6% | 14.3% | 0.804 | 0.648 | 10,102,912 | 8,414,268 | 99.0% | 52.6% | 1.883 | 3.673 |
| Azerbaijan | 9,922,452 | 644,103 | 424,576 | 6.5% | 4.3% | 0.255 | 0.166 | 9,163,779 | 8,468,038 | 92.4% | 6.5% | 14.227 | 1.517 |
| Hungary | 9,861,803 | 912,881 | 875,976 | 9.3% | 8.9% | 0.405 | 0.337 | 9,825,901 | 9,828,569 | 99.6% | 9.3% | 10.764 | 1.042 |
| Belarus | 9,555,690 | 1,192,055 | 1,175,570 | 12.5% | 12.3% | 0.325 | 0.268 | 9,541,249 | 9,551,242 | 99.8% | 12.5% | 8.004 | 1.014 |
| Sweden | 9,512,654 | 4,154,165 | 3,511,365 | 43.7% | 36.9% | 1.321 | 0.470 | 8,870,504 | 8,881,920 | 93.2% | 43.7% | 2.135 | 1.183 |
| Jordan | 9,225,528 | 289,148 | 151,580 | 3.1% | 1.6% | 2.156 | 2.034 | 8,702,319 | 7,810,837 | 94.3% | 3.1% | 30.096 | 1.908 |

| | | | | | | | | | | | | | |
|---|---|---|---|---|---|---|---|---|---|---|---|---|---|
| Honduras | 8,950,218 | 2,608,031 | 952,404 | 29.1% | 10.6% | 0.566 | 0.203 | 8,849,393 | 8,283,556 | 98.9% | 29.1% | 3.393 | 2.738 |
| Austria | 8,781,279 | 2,669,998 | 2,898,054 | 30.4% | 33.0% | 0.514 | 0.464 | 8,773,419 | 8,781,213 | 99.9% | 30.4% | 3.286 | 0.921 |
| Tajikistan | 8,509,930 | 727,363 | 749,831 | 8.5% | 8.8% | 0.255 | 0.242 | 8,072,251 | 8,037,731 | 94.9% | 8.5% | 11.098 | 0.970 |
| Switzerland | 8,301,755 | 2,585,189 | 2,820,431 | 31.1% | 34.0% | 0.548 | 0.485 | 8,290,655 | 8,301,406 | 99.9% | 31.1% | 3.207 | 0.917 |
| Somalia | 7,992,709 | 3,828,754 | 2,390,691 | 47.9% | 29.9% | 0.784 | 0.512 | 7,605,098 | 4,815,115 | 95.2% | 47.9% | 1.986 | 1.602 |
| Togo | 7,982,507 | 1,256,002 | 994,683 | 15.7% | 12.5% | 0.372 | 0.276 | 6,380,166 | 5,516,143 | 79.9% | 15.7% | 5.080 | 1.263 |
| Israel | 7,629,219 | 508,231 | 99,622 | 6.7% | 1.3% | 1.102 | 0.388 | 7,366,358 | 3,359,366 | 96.6% | 6.7% | 14.494 | 5.102 |
| Republic of Serbia | 7,163,415 | 644,937 | 694,465 | 9.0% | 9.7% | 0.257 | 0.240 | 7,078,879 | 7,021,272 | 98.8% | 9.0% | 10.976 | 0.929 |
| Bulgaria | 7,106,711 | 764,233 | 709,872 | 10.8% | 10.0% | 0.334 | 0.250 | 6,786,060 | 6,693,985 | 95.5% | 10.8% | 8.880 | 1.077 |
| Laos | 7,073,997 | 2,084,199 | 2,097,934 | 29.5% | 29.7% | 0.430 | 0.369 | 6,742,622 | 6,799,999 | 95.3% | 29.5% | 3.235 | 0.993 |
| Paraguay | 6,894,793 | 1,814,242 | 2,030,354 | 26.3% | 29.4% | 0.420 | 0.384 | 6,856,637 | 6,888,175 | 99.4% | 26.3% | 3.779 | 0.894 |
| Hong Kong S.A.R. | 6,849,951 | 2,881,166 | 44,970 | 42.1% | 0.7% | 0.816 | 0.132 | 6,269,084 | 5,733,951 | 91.5% | 42.1% | 2.176 | 64.069 |
| Libya | 6,731,508 | 563,938 | 6,284 | 8.4% | 0.1% | 9.879 | 3.044 | 6,146,587 | 337,977 | 91.3% | 8.4% | 10.899 | 89.742 |
| Papua New Guinea | 6,466,968 | 3,238,004 | 2,841,604 | 50.1% | 43.9% | 0.958 | 0.518 | 6,111,465 | 6,097,351 | 94.5% | 50.1% | 1.887 | 1.139 |
| Lebanon | 6,200,285 | 1,805,177 | 227,518 | 29.1% | 3.7% | 1.015 | 0.838 | 6,147,984 | 5,038,301 | 99.2% | 29.1% | 3.406 | 7.934 |
| El Salvador | 6,133,006 | 2,217,125 | 175,518 | 36.2% | 2.9% | 0.585 | 0.179 | 6,118,290 | 5,618,695 | 99.8% | 36.2% | 2.760 | 12.632 |
| Nicaragua | 5,988,125 | 1,796,236 | 584,101 | 30.0% | 9.8% | 0.691 | 0.253 | 5,881,397 | 5,387,183 | 98.2% | 30.0% | 3.274 | 3.075 |
| Sierra Leone | 5,934,755 | 1,230,334 | 533,348 | 20.7% | 9.0% | 0.797 | 0.387 | 5,315,207 | 4,364,883 | 89.6% | 20.7% | 4.320 | 2.307 |
| Eritrea | 5,887,782 | 2,133,453 | 749,195 | 36.2% | 12.7% | 1.574 | 1.083 | 5,481,449 | 4,534,726 | 93.1% | 36.2% | 2.569 | 2.848 |
| Kyrgyzstan | 5,722,700 | 728,242 | 766,025 | 12.7% | 13.4% | 0.296 | 0.242 | 5,678,312 | 5,681,381 | 99.2% | 12.7% | 7.797 | 0.951 |
| Singapore | 5,606,590 | 500,881 | 157,043 | 8.9% | 2.8% | 0.505 | 0.303 | 5,493,693 | 5,218,573 | 98.0% | 8.9% | 10.968 | 3.189 |
| Central African Republic | 5,603,671 | 3,449,326 | 3,544,657 | 61.6% | 63.3% | 0.762 | 0.646 | 5,572,435 | 5,601,786 | 99.4% | 61.6% | 1.616 | 0.973 |
| United Arab Emirates | 5,501,365 | 457,274 | 23 | 8.3% | 0.0% | 2.075 | 0.012 | 4,779,657 | 172,563 | 86.9% | 8.3% | 10.453 | ######## |
| Slovakia | 5,429,724 | 817,470 | 945,823 | 15.1% | 17.4% | 0.338 | 0.334 | 5,424,595 | 5,425,368 | 99.9% | 15.1% | 6.636 | 0.864 |
| Denmark | 5,351,816 | 1,333,497 | 126,759 | 24.9% | 2.4% | 1.205 | 0.370 | 5,277,334 | 4,486,513 | 98.6% | 24.9% | 3.958 | 10.520 |
| Turkmenistan | 5,331,160 | 272,305 | 158,775 | 5.1% | 3.0% | 0.433 | 0.224 | 4,207,239 | 2,700,081 | 78.9% | 5.1% | 15.450 | 1.715 |
| Finland | 5,308,910 | 2,966,577 | 2,681,338 | 55.9% | 50.5% | 1.350 | 0.604 | 5,113,781 | 5,189,877 | 96.3% | 55.9% | 1.724 | 1.106 |
| Palestine | 5,257,405 | 409,933 | 151,082 | 7.8% | 2.9% | 0.352 | 0.205 | 4,770,632 | 1,738,892 | 90.7% | 7.8% | 11.638 | 2.713 |
| Republic of the Congo | 4,995,844 | 1,633,168 | 1,258,645 | 32.7% | 25.2% | 0.479 | 0.282 | 4,841,650 | 4,706,341 | 96.9% | 32.7% | 2.965 | 1.298 |
| Costa Rica | 4,918,985 | 1,435,492 | 559,949 | 29.2% | 11.4% | 0.526 | 0.216 | 4,898,733 | 4,859,068 | 99.6% | 29.2% | 3.413 | 2.564 |
| Georgia | 4,888,034 | 863,572 | 820,924 | 17.7% | 16.8% | 0.422 | 0.277 | 4,799,121 | 4,824,696 | 98.2% | 17.7% | 5.557 | 1.052 |
| Ireland | 4,886,468 | 2,655,786 | 2,178,225 | 54.3% | 44.6% | 0.736 | 0.593 | 4,858,876 | 4,747,318 | 99.4% | 54.3% | 1.830 | 1.219 |
| Norway | 4,564,747 | 2,113,438 | 1,317,449 | 46.3% | 28.9% | 1.800 | 0.773 | 4,123,817 | 3,928,572 | 90.3% | 46.3% | 1.951 | 1.604 |
| Liberia | 4,326,013 | 820,020 | 465,885 | 19.0% | 10.8% | 0.615 | 0.238 | 3,265,752 | 3,108,127 | 75.5% | 19.0% | 3.983 | 1.760 |
| New Zealand | 4,066,381 | 1,942,277 | 644,354 | 47.8% | 15.8% | 0.790 | 0.252 | 3,911,698 | 3,632,233 | 96.2% | 47.8% | 2.014 | 3.014 |
| Croatia | 4,022,835 | 757,206 | 547,245 | 18.8% | 13.6% | 0.435 | 0.280 | 3,882,896 | 3,797,386 | 96.5% | 18.8% | 5.128 | 1.384 |
| Bosnia and Herzegovina | 3,886,733 | 895,166 | 1,007,049 | 23.0% | 25.9% | 0.403 | 0.377 | 3,875,403 | 3,881,522 | 99.7% | 23.0% | 4.329 | 0.889 |
| Mauritania | 3,767,659 | 1,689,921 | 2,317 | 44.9% | 0.1% | 3.839 | 2.465 | 3,347,098 | 58,646 | 88.8% | 44.9% | 1.981 | 729.357 |
| Panama | 3,656,505 | 965,393 | 474,478 | 26.4% | 13.0% | 0.597 | 0.218 | 3,619,795 | 3,532,853 | 99.0% | 26.4% | 3.750 | 2.035 |
| Moldova | 3,415,404 | 106,241 | 99,321 | 3.1% | 2.9% | 0.442 | 0.365 | 2,908,086 | 2,900,367 | 85.1% | 3.1% | 27.373 | 1.070 |
| Oman | 3,389,456 | 439,409 | - | 13.0% | 0.0% | 3.231 | | 2,788,494 | - | 82.3% | 13.0% | 6.346 | |
| Uruguay | 3,354,933 | 694,467 | 70,065 | 20.7% | 2.1% | 0.309 | 0.320 | 3,263,163 | 2,581,882 | 97.3% | 20.7% | 4.699 | 9.912 |
| Puerto Rico | 3,309,835 | 1,439,071 | 462,468 | 43.5% | 14.0% | 0.622 | 0.333 | 3,305,405 | 3,299,759 | 99.9% | 43.5% | 2.297 | 3.112 |
| Somaliland | 3,204,281 | 1,632,447 | 58,751 | 50.9% | 1.8% | 0.991 | 0.248 | 3,099,928 | 1,142,552 | 96.7% | 50.9% | 1.899 | 27.786 |
| Mongolia | 3,073,670 | 1,039,039 | 670,406 | 33.8% | 21.8% | 1.137 | 0.745 | 2,800,905 | 2,805,765 | 91.1% | 33.8% | 2.696 | 1.550 |
| Armenia | 3,030,410 | 196,721 | 199,043 | 6.5% | 6.6% | 0.225 | 0.183 | 2,998,862 | 2,998,128 | 99.0% | 6.5% | 15.244 | 0.988 |
| Albania | 3,029,644 | 341,638 | 270,951 | 11.3% | 8.9% | 0.292 | 0.218 | 2,990,849 | 2,791,399 | 98.7% | 11.3% | 8.754 | 1.261 |
| Jamaica | 2,893,649 | 1,026,859 | 290,166 | 35.5% | 10.0% | 0.557 | 0.232 | 2,841,701 | 2,755,103 | 98.2% | 35.5% | 2.767 | 3.539 |
| Kuwait | 2,852,295 | 126,841 | - | 4.4% | 0.0% | 3.545 | 0.000 | 2,379,930 | 8 | 83.4% | 4.4% | 18.763 | |
| Lithuania | 2,818,468 | 467,998 | 466,740 | 16.6% | 16.6% | 0.467 | 0.404 | 2,815,317 | 2,815,297 | 99.9% | 16.6% | 6.016 | 1.003 |

| | | | | | | | | | | | | | |
|---|---|---|---|---|---|---|---|---|---|---|---|---|---|
| Namibia | 2,494,956 | 1,303,297 | 673,342 | 52.2% | 27.0% | 0.925 | 0.795 | 2,423,723 | 2,036,850 | 97.1% | 52.2% | 1.860 | 1.936 |
| Qatar | 2,228,738 | 93,322 | - | 4.2% | 0.0% | 1.807 | | 2,108,786 | - | 94.6% | 4.2% | 22.597 | |
| Botswana | 2,189,399 | 1,181,943 | 796,135 | 54.0% | 36.4% | 0.996 | 0.642 | 2,158,948 | 2,098,935 | 98.6% | 54.0% | 1.827 | 1.485 |
| Macedonia | 2,108,556 | 232,900 | 242,807 | 11.0% | 11.5% | 0.235 | 0.199 | 2,088,409 | 2,103,366 | 99.0% | 11.0% | 8.967 | 0.959 |
| Gambia | 1,997,898 | 205,183 | 27,520 | 10.3% | 1.4% | 0.381 | 0.120 | 1,918,208 | 1,552,235 | 96.0% | 10.3% | 9.349 | 7.456 |
| Slovenia | 1,987,022 | 562,328 | 632,161 | 28.3% | 31.8% | 0.494 | 0.449 | 1,984,832 | 1,984,417 | 99.9% | 28.3% | 3.530 | 0.890 |
| Lesotho | 1,953,018 | 702,643 | 16,581 | 36.0% | 0.8% | 0.566 | 0.206 | 1,934,752 | 824,693 | 99.1% | 36.0% | 2.754 | 42.376 |
| Latvia | 1,933,565 | 550,716 | 488,407 | 28.5% | 25.3% | 0.491 | 0.380 | 1,931,377 | 1,917,314 | 99.9% | 28.5% | 3.507 | 1.128 |
| Kosovo | 1,890,703 | 157,436 | 202,081 | 8.3% | 10.7% | 0.232 | 0.219 | 1,883,698 | 1,890,703 | 99.6% | 8.3% | 11.965 | 0.779 |
| Guinea-Bissau | 1,746,002 | 661,910 | 194,476 | 37.9% | 11.1% | 0.668 | 0.290 | 1,661,447 | 1,221,221 | 95.2% | 37.9% | 2.510 | 3.404 |
| Gabon | 1,738,438 | 837,197 | 412,298 | 48.2% | 23.7% | 0.729 | 0.253 | 1,672,469 | 1,700,572 | 96.2% | 48.2% | 1.998 | 2.031 |
| eSwatini | 1,456,471 | 521,795 | 359,320 | 35.8% | 24.7% | 0.678 | 0.539 | 1,454,833 | 1,446,994 | 99.9% | 35.8% | 2.788 | 1.452 |
| Mauritius | 1,326,444 | 278,406 | - | 21.0% | 0.0% | 1.123 | | 1,268,232 | - | 95.6% | 21.0% | 4.555 | |
| East Timor | 1,255,920 | 129,202 | 63,789 | 10.3% | 5.1% | 0.860 | 0.399 | 1,075,754 | 933,269 | 85.7% | 10.3% | 8.326 | 2.025 |
| Estonia | 1,221,733 | 442,504 | 293,109 | 36.2% | 24.0% | 0.626 | 0.408 | 1,214,456 | 1,199,253 | 99.4% | 36.2% | 2.745 | 1.510 |
| Trinidad and Tobago | 1,189,509 | 464,267 | 188,626 | 39.0% | 15.9% | 0.673 | 0.228 | 1,181,365 | 1,163,757 | 99.3% | 39.0% | 2.545 | 2.461 |
| Bahrain | 1,038,211 | 295,148 | 524 | 28.4% | 0.1% | 1.324 | 0.075 | 1,025,903 | 847,945 | 98.8% | 28.4% | 3.476 | 563.260 |
| Bhutan | 870,394 | 547,289 | 587,242 | 62.9% | 67.5% | 0.887 | 0.737 | 863,279 | 870,153 | 99.2% | 62.9% | 1.577 | 0.932 |
| Djibouti | 821,637 | 339,968 | 4,794 | 41.4% | 0.6% | 6.499 | 0.220 | 691,391 | 22,949 | 84.1% | 41.4% | 2.034 | 70.915 |
| Fiji | 813,134 | 416,920 | 61,601 | 51.3% | 7.6% | 0.843 | 0.466 | 764,013 | 565,666 | 94.0% | 51.3% | 1.833 | 6.768 |
| Guyana | 716,848 | 369,757 | 219,607 | 51.6% | 30.6% | 1.267 | 0.313 | 703,511 | 701,780 | 98.1% | 51.6% | 1.903 | 1.684 |
| Cyprus | 700,731 | 96,551 | 2,299 | 13.8% | 0.3% | 0.304 | 0.049 | 676,970 | 462,343 | 96.6% | 13.8% | 7.012 | 41.997 |
| Equatorial Guinea | 680,451 | 440,111 | 248,675 | 64.7% | 36.5% | 0.900 | 0.410 | 646,274 | 672,947 | 95.0% | 64.7% | 1.468 | 1.770 |
| Comoros | 678,746 | 417,477 | - | 61.5% | 0.0% | 2.866 | | 583,383 | - | 86.0% | 61.5% | 1.397 | |
| Montenegro | 606,998 | 219,118 | 202,229 | 36.1% | 33.3% | 0.560 | 0.413 | 604,210 | 605,488 | 99.5% | 36.1% | 2.757 | 1.084 |
| Luxembourg | 596,003 | 158,107 | 187,193 | 26.5% | 31.4% | 0.531 | 0.550 | 591,096 | 595,858 | 99.2% | 26.5% | 3.739 | 0.845 |
| Suriname | 584,869 | 235,434 | 128,597 | 40.3% | 22.0% | 0.954 | 0.225 | 578,641 | 582,624 | 98.9% | 40.3% | 2.458 | 1.831 |
| Macao S.A.R | 564,113 | 396,532 | 333,915 | 70.3% | 59.2% | 1.406 | 2.960 | 555,588 | 484,778 | 98.5% | 70.3% | 1.401 | 1.188 |
| Cabo Verde | 452,747 | 151,128 | - | 33.4% | 0.0% | 3.245 | | 328,105 | - | 72.5% | 33.4% | 2.171 | |
| Brunei | 419,129 | 125,571 | 76,165 | 30.0% | 18.2% | 0.447 | 0.213 | 413,190 | 417,104 | 98.6% | 30.0% | 3.290 | 1.649 |
| Malta | 409,763 | 48,602 | 558 | 11.9% | 0.1% | 0.873 | 0.055 | 401,267 | 395,230 | 97.9% | 11.9% | 8.256 | 87.100 |
| Solomon Islands | 400,839 | 287,013 | 205,198 | 71.6% | 51.2% | 1.447 | 0.616 | 348,275 | 339,105 | 86.9% | 71.6% | 1.213 | 1.399 |
| Northern Cyprus | 383,550 | 65,548 | 217 | 17.1% | 0.1% | 0.549 | 0.047 | 372,626 | 286,893 | 97.2% | 17.1% | 5.685 | 302.065 |
| Belize | 338,621 | 154,697 | 107,874 | 45.7% | 31.9% | 0.759 | 0.400 | 315,679 | 306,368 | 93.2% | 45.7% | 2.041 | 1.434 |
| Iceland | 306,970 | 238,873 | 2 | 77.8% | 0.0% | 3.586 | 0.055 | 274,901 | 29 | 89.6% | 77.8% | 1.151 | ######## |
| The Bahamas | 289,725 | 146,320 | 14,240 | 50.5% | 4.9% | 0.950 | 0.095 | 283,726 | 233,410 | 97.9% | 50.5% | 1.939 | 10.275 |
| Barbados | 247,885 | 86,227 | 4,957 | 34.8% | 2.0% | 1.095 | 0.238 | 241,653 | 228,013 | 97.5% | 34.8% | 2.803 | 17.395 |
| New Caledonia | 243,095 | 126,420 | 15,399 | 52.0% | 6.3% | 1.359 | 0.473 | 228,874 | 161,268 | 94.2% | 52.0% | 1.810 | 8.210 |
| Vanuatu | 216,480 | 185,630 | 50,705 | 85.7% | 23.4% | 1.773 | 0.358 | 204,728 | 170,441 | 94.6% | 85.7% | 1.103 | 3.661 |
| French Polynesia | 203,189 | 109,532 | - | 53.9% | 0.0% | 2.800 | | 179,927 | - | 88.6% | 53.9% | 1.643 | |
| São Tomé and Príncipe | 188,186 | 55,261 | - | 29.4% | 0.0% | 2.048 | | 183,842 | - | 97.7% | 29.4% | 3.327 | |
| Guam | 163,406 | 90,631 | - | 55.5% | 0.0% | 1.806 | | 161,052 | - | 98.6% | 55.5% | 1.777 | |
| Saint Lucia | 157,880 | 118,161 | 62,153 | 74.8% | 39.4% | 1.081 | 0.480 | 156,947 | 157,312 | 99.4% | 74.8% | 1.328 | 1.901 |
| Maldives | 157,785 | 155,935 | - | 98.8% | 0.0% | 1.186 | | - | - | 0.0% | 98.8% | - | |
| Curaçao | 147,442 | 59,581 | 164 | 40.4% | 0.1% | 0.794 | 0.021 | 147,429 | 6,455 | 100.0% | 40.4% | 2.474 | 363.299 |
| Samoa | 146,519 | 67,695 | - | 46.2% | 0.0% | 5.010 | | 131,994 | - | 90.1% | 46.2% | 1.950 | |
| Cyprus No Mans Area | 120,703 | 7,102 | - | 5.9% | 0.0% | 0.183 | | 120,703 | 114,413 | 100.0% | 5.9% | 16.996 | |
| United States Virgin Islands | 104,176 | 74,879 | 15,982 | 71.9% | 15.3% | 0.938 | 0.476 | 102,926 | 96,306 | 98.8% | 71.9% | 1.375 | 4.685 |
| Grenada | 95,383 | 84,389 | 65,800 | 88.5% | 69.0% | 1.132 | 0.769 | 94,567 | 92,842 | 99.1% | 88.5% | 1.121 | 1.283 |
| Seychelles | 88,552 | 40,430 | - | 45.7% | 0.0% | 0.711 | | 76,634 | - | 86.5% | 45.7% | 1.895 | |

| | | | | | | | | | | | | | |
|---|---|---|---|---|---|---|---|---|---|---|---|---|---|
| Antigua and Barbuda | 85,449 | 74,608 | 1,042 | 87.3% | 1.2% | 1.113 | 0.067 | 85,417 | 84,898 | 100.0% | 87.3% | 1.145 | 71.601 |
| Isle of Man | 85,235 | 21,721 | 33,092 | 25.5% | 38.8% | 0.552 | 0.498 | 71,053 | 83,612 | 83.4% | 25.5% | 3.271 | 0.656 |
| Andorra | 84,995 | 68,752 | 70,641 | 80.9% | 83.1% | 0.985 | 0.887 | 84,995 | 84,995 | 100.0% | 80.9% | 1.236 | 0.973 |
| Aruba | 80,565 | 52,649 | - | 65.3% | 0.0% | 1.081 | | 80,562 | - | 100.0% | 65.3% | 1.530 | |
| Saint Vincent and the Grenadines | 71,274 | 60,619 | 27,840 | 85.1% | 39.1% | 1.423 | 0.485 | 71,025 | 68,043 | 99.7% | 85.1% | 1.172 | 2.177 |
| Bermuda | 63,562 | 48,869 | - | 76.9% | 0.0% | 1.089 | | - | - | 0.0% | 76.9% | - | |
| Guernsey | 61,621 | 31,938 | - | 51.8% | 0.0% | 1.166 | | 35,840 | - | 58.2% | 51.8% | 1.122 | |
| Jersey | 60,305 | 3,497 | 933 | 5.8% | 1.5% | 0.265 | 0.495 | 50,871 | 50,221 | 84.4% | 5.8% | 14.547 | 3.748 |
| Dominica | 59,013 | 54,652 | 33,176 | 92.6% | 56.2% | 1.057 | 0.611 | 58,490 | 58,571 | 99.1% | 92.6% | 1.070 | 1.647 |
| Cayman Islands | 57,107 | 56,103 | 10,551 | 98.2% | 18.5% | 1.127 | 0.328 | 56,979 | 37,326 | 99.8% | 98.2% | 1.016 | 5.317 |
| Tonga | 51,480 | 21,039 | - | 40.9% | 0.0% | 1.061 | | 45,983 | - | 89.3% | 40.9% | 2.186 | |
| Northern Mariana Islands | 50,350 | 37,918 | - | 75.3% | 0.0% | 3.523 | | 47,492 | - | 94.3% | 75.3% | 1.252 | |
| American Samoa | 45,555 | 16,809 | - | 36.9% | 0.0% | 1.513 | | 40,739 | - | 89.4% | 36.9% | 2.424 | |
| Turks and Caicos Islands | 44,265 | 27,501 | 18,297 | 62.1% | 41.3% | 0.972 | 0.950 | 40,358 | 41,251 | 91.2% | 62.1% | 1.468 | 1.503 |
| Federated States of Micronesia | 41,989 | 29,832 | - | 71.0% | 0.0% | 2.409 | | 38,077 | - | 90.7% | 71.0% | 1.276 | |
| Saint Martin | 41,301 | 35,163 | 711 | 85.1% | 1.7% | 0.901 | 0.103 | 41,050 | 23,182 | 99.4% | 85.1% | 1.167 | 49.456 |
| Western Sahara | 40,247 | 852 | - | 2.1% | 0.0% | 96.003 | | 1,727 | - | 4.3% | 2.1% | 2.027 | |
| Monaco | 40,195 | 32,366 | - | 80.5% | 0.0% | 1.610 | | 40,195 | - | 100.0% | 80.5% | 1.242 | |
| Saint Kitts and Nevis | 39,936 | 35,673 | 740 | 89.3% | 1.9% | 1.030 | 0.070 | 39,855 | 39,855 | 99.8% | 89.3% | 1.117 | 48.207 |
| Faroe Islands | 37,249 | 24,819 | - | 66.6% | 0.0% | 1.859 | | 32,659 | - | 87.7% | 66.6% | 1.316 | |
| Liechtenstein | 34,641 | 12,587 | 14,603 | 36.3% | 42.2% | 0.640 | 0.650 | 34,640 | 34,641 | 100.0% | 36.3% | 2.752 | 0.862 |
| British Virgin Islands | 31,118 | 31,108 | 6,413 | 100.0% | 20.6% | 1.151 | 0.392 | 30,766 | 27,454 | 98.9% | 100.0% | 0.989 | 4.851 |
| San Marino | 30,139 | - | - | 0.0% | 0.0% | | | 2,020 | 2,020 | 6.7% | 0.0% | | |
| Kiribati | 26,147 | 22,089 | - | 84.5% | 0.0% | 1.569 | | 15,244 | - | 58.3% | 84.5% | 0.690 | |
| Aland | 24,202 | 16,354 | 910 | 67.6% | 3.8% | 1.509 | 0.432 | 16,923 | 16,272 | 69.9% | 67.6% | 1.035 | 17.971 |
| Sint Maarten | 22,221 | 19,133 | 1,779 | 86.1% | 8.0% | 1.033 | 0.480 | 22,221 | 12,257 | 100.0% | 86.1% | 1.161 | 10.755 |
| Dhekelia Sovereign Base Area | 16,445 | 1,244 | - | 7.6% | 0.0% | 0.454 | | 14,582 | 14 | 88.7% | 7.6% | 11.722 | |
| Greenland | 16,269 | 8,845 | - | 54.4% | 0.0% | | | 10,579 | - | 65.0% | 54.4% | 1.196 | |
| Palau | 14,500 | 12,462 | - | 85.9% | 0.0% | 2.755 | | 13,545 | - | 93.4% | 85.9% | 1.087 | |
| Anguilla | 14,113 | 12,681 | - | 89.9% | 0.0% | 1.219 | | 13,168 | - | 93.3% | 89.9% | 1.038 | |
| Gibraltar | 9,176 | 9,176 | - | 100.0% | 0.0% | 1.000 | | 9,176 | - | 100.0% | 100.0% | 1.000 | |
| Wallis and Futuna | 9,060 | 5,431 | - | 59.9% | 0.0% | 1.073 | | 7,123 | - | 78.6% | 59.9% | 1.312 | |
| Nauru | 7,641 | 6,939 | - | 90.8% | 0.0% | 1.211 | | 7,436 | - | 97.3% | 90.8% | 1.072 | |
| Saint Helena | 6,998 | - | - | 0.0% | 0.0% | | | - | - | 0.0% | 0.0% | | |
| Saint Barthelemy | 6,772 | 6,772 | - | 100.0% | 0.0% | 1.000 | | 6,772 | - | 100.0% | 100.0% | 1.000 | |
| Cook Islands | 5,987 | 4,798 | - | 80.1% | 0.0% | 3.424 | | 3,753 | - | 62.7% | 80.1% | 0.782 | |
| Saint Pierre and Miquelon | 5,456 | 4,827 | 109 | 88.5% | 2.0% | 4.954 | 0.093 | 5,404 | 127 | 99.0% | 88.5% | 1.120 | 44.284 |
| Akrotiri Sovereign Base Area | 5,389 | 1,359 | 187 | 25.2% | 3.5% | 1.093 | 0.301 | 4,663 | 3,234 | 86.5% | 25.2% | 3.431 | 7.267 |
| Montserrat | 4,247 | 3,877 | 835 | 91.3% | 19.7% | 1.521 | 0.328 | 4,245 | 4,247 | 100.0% | 91.3% | 1.095 | 4.643 |
| Marshall Islands | 4,024 | 1,755 | - | 43.6% | 0.0% | 0.463 | | 3,453 | - | 85.8% | 43.6% | 1.968 | |
| Falkland Islands | 2,816 | 33 | - | 1.2% | 0.0% | 0.185 | | 2,748 | - | 97.6% | 1.2% | 83.273 | |
| Norfolk Island | 2,180 | 88 | - | 4.0% | 0.0% | 0.323 | | 1,780 | - | 81.7% | 4.0% | 20.227 | |
| Niue | 1,071 | 728 | - | 68.0% | 0.0% | 4.078 | | 1,026 | - | 95.8% | 68.0% | 1.409 | |
| Vatican | 1,000 | - | - | 0.0% | 0.0% | | | 1,000 | 1,000 | 100.0% | 0.0% | | |
| Tuvalu | 971 | 564 | - | 58.1% | 0.0% | 0.968 | | 917 | - | 94.4% | 58.1% | 1.626 | |
| Siachen Glacier | 239 | - | - | 0.0% | 0.0% | | 0.000 | - | 239 | 0.0% | 0.0% | | |
| Indian Ocean Territories | 217 | 217 | - | 100.0% | 0.0% | 2.364 | | 217 | - | 100.0% | 100.0% | 1.000 | |
| US Naval Base Guantanamo Bay | 175 | 28 | 151 | 16.0% | 86.3% | 0.427 | 0.986 | 127 | 175 | 72.6% | 16.0% | 4.536 | 0.185 |
| Spratly Islands | 69 | 32 | - | 46.4% | 0.0% | 0.464 | | - | - | 0.0% | 46.4% | - | |
| Pitcairn Islands | 15 | - | - | 0.0% | 0.0% | | | - | - | 0.0% | 0.0% | | |

| | | | | | | | | | |
|---|---|---|---|---|---|---|---|---|---|
| United States Minor Outlying Islands | 15 | - | - | 0.0% | 0.0% | - | - | 0.0% | 0.0% |
| French Southern and Antarctic Lanc | - | - | - | | | - | - | 0.0% | 0.0% |
| South Georgia and the Islands | - | - | - | | | - | - | 0.0% | 0.0% |
| Heard Island and McDonald Islands | - | - | - | | | - | - | 0.0% | 0.0% |
| British Indian Ocean Territory | - | - | - | | | - | - | 0.0% | 0.0% |
| Clipperton Island | - | - | - | | | - | - | 0.0% | 0.0% |
| Scarborough Reef | - | - | - | | | - | - | 0.0% | 0.0% |
| Serranilla Bank | - | - | - | | | - | - | 0.0% | 0.0% |
| Coral Sea Islands | - | - | - | | | - | - | 0.0% | 0.0% |
| Bajo Nuevo Bank (Petrel Is.) | - | - | - | | | - | - | 0.0% | 0.0% |

**SI Table 3. Overlap of critical natural assets with biodiversity.**

| | | Number of species | Percent of species within LCNA | Percent of species within GCNA |
|---|---|---|---|---|
| Global totals | birds | 13077 | 73% | 73% |
| | mammals | 5102 | 66% | 70% |
| | reptiles | 4031 | 41% | 41% |
| | amphibians | 5967 | 41% | 46% |
| | all vertebrates | 28177 | 60% | 62% |
| Endemic species (AOH 100% within country) | birds | 3435 | 45% | 50% |
| | mammals | 2027 | 41% | 47% |
| | reptiles | 2484 | 26% | 26% |
| | amphibians | 4033 | 26% | 30% |
| | all vertebrates | 11979 | 34% | 37% |
| Species whose AOH falls >80% within country | birds | 5484 | 55% | 58% |
| | mammals | 2796 | 50% | 54% |
| | reptiles | 3048 | 30% | 31% |
| | amphibians | 4725 | 30% | 35% |
| | all vertebrates | 16053 | 42% | 45% |
| Species whose AOH falls >50% within country | birds | 9638 | 61% | 65% |
| | mammals | 4160 | 57% | 61% |
| | reptiles | 3748 | 34% | 36% |
| | amphibians | 5583 | 34% | 39% |
| | all vertebrates | 23129 | 50% | 53% |

| Country (ranked by number of endemic + majority AOH species) | # Endemic species (whose area of habitat (AOH) is entirely contained within country) | Percent of endemic species within critical natural assets for local NCP (LCNA) | Percent of endemic species within critical natural assets for global NCP (GCNA) | # Species whose AOH falls >50% within country | Percent of >50% AOH species within LCNA | Percent of >50% AOH species within GCNA |
|---|---|---|---|---|---|---|
| Global | 11979 | 34% | 37% | 23129 | 50% | 53% |
| Brazil | 976 | 61% | 62% | 2360 | 82% | 83% |
| Indonesia | 982 | 31% | 58% | 1729 | 51% | 71% |
| Mexico | 880 | 40% | 25% | 1433 | 55% | 42% |
| Australia | 960 | 54% | 48% | 1150 | 57% | 51% |
| Madagascar | 931 | 46% | 54% | 965 | 47% | 55% |
| United States of America | 444 | 55% | 64% | 1208 | 69% | 76% |
| Colombia | 485 | 25% | 40% | 962 | 48% | 63% |
| China | 362 | 52% | 49% | 1063 | 66% | 68% |
| Peru | 449 | 35% | 35% | 921 | 57% | 54% |
| India | 470 | 27% | 26% | 877 | 40% | 38% |
| Philippines | 596 | 31% | 52% | 619 | 32% | 52% |
| Papua New Guinea | 405 | 21% | 55% | 785 | 47% | 71% |
| Argentina | 252 | 29% | 8% | 568 | 56% | 32% |
| Venezuela | 267 | 24% | 32% | 480 | 43% | 51% |
| Ecuador | 248 | 17% | 28% | 497 | 28% | 47% |
| Democratic Republic of the Congo | 92 | 33% | 41% | 426 | 72% | 77% |
| United Republic of Tanzania | 160 | 4% | 10% | 285 | 25% | 34% |
| Russia | 46 | 9% | 20% | 369 | 62% | 76% |
| South Africa | 111 | 58% | 1% | 292 | 77% | 16% |
| Malaysia | 109 | 16% | 30% | 291 | 44% | 64% |

| | | | | | | |
|---|---|---|---|---|---|---|
| Bolivia | 102 | 38% | 55% | 264 | 61% | 74% |
| Chile | 103 | 31% | 14% | 245 | 36% | 36% |
| Japan | 132 | 39% | 30% | 180 | 44% | 36% |
| Vietnam | 89 | 6% | 20% | 200 | 22% | 47% |
| Sri Lanka | 141 | 30% | 8% | 146 | 30% | 8% |
| Costa Rica | 61 | 5% | 5% | 207 | 7% | 20% |
| Cuba | 111 | 29% | 0% | 143 | 34% | 0% |
| Canada | 9 | 22% | 22% | 231 | 50% | 74% |
| Ethiopia | 75 | 21% | 31% | 164 | 54% | 59% |
| Cameroon | 76 | 14% | 18% | 161 | 41% | 45% |
| New Caledonia | 108 | 0% | 0% | 120 | 0% | 0% |
| Solomon Islands | 86 | 15% | 43% | 129 | 12% | 57% |
| New Zealand | 92 | 18% | 30% | 112 | 21% | 28% |
| Panama | 63 | 10% | 13% | 141 | 16% | 31% |
| Guatemala | 58 | 21% | 26% | 122 | 30% | 34% |
| Kenya | 49 | 12% | 18% | 125 | 37% | 49% |
| Myanmar | 43 | 9% | 35% | 130 | 45% | 67% |
| Honduras | 66 | 2% | 6% | 97 | 12% | 12% |
| Dominican Republic | 33 | 3% | 6% | 119 | 18% | 6% |
| Iran | 42 | 2% | 2% | 108 | 6% | 12% |
| Taiwan | 67 | 40% | 4% | 76 | 38% | 4% |
| Jamaica | 71 | 37% | 0% | 71 | 37% | 0% |
| Spain | 36 | 6% | 8% | 90 | 42% | 38% |
| Angola | 38 | 50% | 50% | 84 | 69% | 69% |
| Turkey | 32 | 3% | 3% | 83 | 27% | 33% |
| Yemen | 38 | 0% | 0% | 72 | 13% | 0% |
| France | 34 | 0% | 6% | 72 | 14% | 25% |
| Fiji | 45 | 49% | 0% | 60 | 48% | 0% |
| Thailand | 29 | 0% | 14% | 72 | 7% | 28% |
| Puerto Rico | 40 | 38% | 0% | 58 | 33% | 0% |
| Haiti | 38 | 0% | 0% | 56 | 4% | 0% |
| São Tomé and Príncipe | 44 | 0% | 0% | 45 | 0% | 0% |
| Mozambique | 12 | 0% | 0% | 67 | 58% | 63% |
| Morocco | 25 | 12% | 4% | 50 | 14% | 2% |
| Pakistan | 9 | 11% | 22% | 59 | 22% | 22% |
| Liberia | | | | 68 | 69% | 96% |
| Guyana | 25 | 4% | 16% | 40 | 5% | 33% |
| Italy | 21 | 24% | 19% | 44 | 30% | 30% |
| Namibia | 11 | 27% | 0% | 53 | 49% | 32% |
| Comoros | 28 | 0% | 0% | 35 | 0% | 0% |
| Nepal | 14 | 21% | 36% | 47 | 36% | 40% |
| Kazakhstan | 6 | 50% | 33% | 54 | 74% | 76% |
| Nigeria | 14 | 0% | 7% | 42 | 24% | 31% |
| Seychelles | 26 | 0% | 0% | 26 | 0% | 0% |
| Laos | 9 | 33% | 56% | 43 | 47% | 58% |
| Federated States of Micronesia | 24 | 0% | 0% | 27 | 0% | 0% |
| Algeria | 7 | 14% | 0% | 43 | 12% | 0% |
| Somalia | 11 | 45% | 36% | 38 | 63% | 55% |
| Zambia | 11 | 36% | 0% | 36 | 58% | 0% |
| Cambodia | 7 | 0% | 0% | 40 | 25% | 48% |
| Nicaragua | 6 | 0% | 0% | 41 | 15% | 44% |
| Cabo Verde | 22 | 0% | 0% | 23 | 0% | 0% |
| Vanuatu | 19 | 0% | 53% | 26 | 0% | 54% |
| Trinidad and Tobago | 18 | 0% | 11% | 21 | 0% | 14% |
| Saudi Arabia | 4 | 0% | 0% | 35 | 0% | 0% |

| | | | | | | |
|---|---|---|---|---|---|---|
| Palau | 17 | 0% | 0% | 20 | 0% | 0% |
| The Bahamas | 16 | 6% | 0% | 21 | 5% | 0% |
| Greece | 13 | 8% | 15% | 24 | 8% | 13% |
| French Polynesia | 16 | 0% | 0% | 20 | 0% | 0% |
| Gabon | 8 | 13% | 25% | 28 | 68% | 71% |
| Ivory Coast | 8 | 0% | 13% | 28 | 4% | 18% |
| Uganda | 7 | 0% | 14% | 28 | 0% | 4% |
| Egypt | 13 | 0% | 0% | 21 | 0% | 0% |
| Ghana | 8 | 0% | 0% | 24 | 8% | 21% |
| Mongolia | 1 | 0% | 0% | 31 | 39% | 48% |
| Oman | 9 | 0% | 0% | 20 | 0% | 0% |
| Sudan | 5 | 20% | 0% | 23 | 52% | 35% |
| French Southern and Antarctic Lanc | 4 | 0% | 0% | 23 | 0% | 0% |
| Samoa | 10 | 0% | 0% | 16 | 0% | 0% |
| Malawi | 9 | 0% | 0% | 16 | 0% | 0% |
| Uruguay | 6 | 0% | 0% | 19 | 42% | 0% |
| Guinea | 8 | 0% | 0% | 16 | 6% | 6% |
| Falkland Islands | 7 | 0% | 0% | 17 | 0% | 0% |
| Zimbabwe | 3 | 0% | 0% | 21 | 33% | 0% |
| Afghanistan | 3 | 33% | 33% | 20 | 45% | 55% |
| Somaliland | 6 | 67% | 0% | 14 | 64% | 0% |
| Paraguay | 3 | 33% | 33% | 17 | 59% | 76% |
| Mali | 2 | 50% | 0% | 16 | 75% | 0% |
| Cayman Islands | 8 | 0% | 0% | 8 | 0% | 0% |
| Mauritius | 8 | 0% | 0% | 8 | 0% | 0% |
| South Korea | 5 | 0% | 0% | 11 | 45% | 0% |
| Equatorial Guinea | 7 | 0% | 0% | 8 | 0% | 0% |
| Saint Vincent and the Grenadines | 7 | 0% | 0% | 8 | 0% | 0% |
| Suriname | 5 | 20% | 40% | 10 | 60% | 70% |
| Senegal | 2 | 0% | 0% | 13 | 23% | 0% |
| East Timor | 1 | 0% | 0% | 14 | 0% | 0% |
| Indian Ocean Territories | 7 | 0% | 0% | 7 | 0% | 0% |
| Northern Mariana Islands | 6 | 0% | 0% | 8 | 0% | 0% |
| Turkmenistan | 2 | 50% | 50% | 12 | 8% | 8% |
| Iraq | 1 | 0% | 0% | 13 | 0% | 0% |
| Portugal | 6 | 0% | 0% | 7 | 0% | 0% |
| Grenada | 5 | 0% | 0% | 7 | 0% | 0% |
| Central African Republic | 4 | 25% | 25% | 8 | 63% | 63% |
| Belize | 2 | 0% | 50% | 10 | 0% | 20% |
| United Kingdom | 2 | 50% | 50% | 10 | 20% | 30% |
| South Sudan | 1 | 0% | 0% | 11 | 73% | 91% |
| Republic of the Congo | 4 | 25% | 25% | 7 | 43% | 57% |
| Chad | 3 | 0% | 0% | 8 | 25% | 25% |
| Dominica | 5 | 0% | 0% | 5 | 0% | 0% |
| Montserrat | 5 | 0% | 0% | 5 | 0% | 0% |
| Norfolk Island | 5 | 0% | 0% | 5 | 0% | 0% |
| Saint Lucia | 5 | 0% | 0% | 5 | 0% | 0% |
| Sierra Leone | 4 | 0% | 0% | 6 | 0% | 0% |
| Libya | 3 | 0% | 0% | 7 | 0% | 0% |
| Mauritania | 2 | 0% | 0% | 7 | 29% | 0% |
| Norway | 2 | 0% | 0% | 7 | 14% | 14% |
| Armenia | 3 | 0% | 0% | 5 | 0% | 0% |
| Hong Kong S.A.R. | 3 | 0% | 0% | 5 | 0% | 0% |
| United States Virgin Islands | 3 | 0% | 0% | 5 | 0% | 0% |
| Armenia | 3 | 0% | 0% | 5 | 0% | 0% |

| | | | | | | |
|---|---|---|---|---|---|---|
| Rwanda | 2 | 0% | 0% | 6 | 0% | 0% |
| Syria | 2 | 0% | 0% | 6 | 0% | 0% |
| Cyprus | 1 | 0% | 0% | 7 | 0% | 0% |
| Georgia | 1 | 0% | 0% | 7 | 0% | 14% |
| Bhutan | | | | 8 | 38% | 38% |
| Antigua and Barbuda | 3 | 0% | 0% | 4 | 0% | 0% |
| Botswana | 1 | 0% | 0% | 6 | 83% | 83% |
| Israel | 1 | 0% | 0% | 6 | 0% | 0% |
| Benin | 3 | 0% | 0% | 3 | 0% | 0% |
| Cook Islands | 3 | 0% | 0% | 3 | 0% | 0% |
| Singapore | 3 | 0% | 0% | 3 | 0% | 0% |
| Turks and Caicos Islands | 3 | 0% | 0% | 3 | 0% | 0% |
| Netherlands | 2 | 0% | 0% | 4 | 0% | 0% |
| Aruba | 2 | 0% | 0% | 3 | 0% | 0% |
| Burundi | 2 | 0% | 0% | 3 | 0% | 0% |
| Eritrea | 2 | 0% | 0% | 3 | 0% | 0% |
| Tunisia | 2 | 0% | 0% | 3 | 0% | 0% |
| Curaçao | 2 | 0% | 0% | 3 | 0% | 0% |
| Ukraine | 2 | 0% | 0% | 3 | 33% | 33% |
| South Georgia and the Islands | 1 | 0% | 0% | 4 | 0% | 0% |
| Azerbaijan | 1 | 0% | 0% | 4 | 0% | 0% |
| Bangladesh | | | | 5 | 0% | 0% |
| Greenland | | | | 5 | 0% | 0% |
| Lebanon | | | | 5 | 0% | 0% |
| British Virgin Islands | 2 | 0% | 0% | 2 | 0% | 0% |
| Saint Helena | 2 | 0% | 0% | 2 | 0% | 0% |
| Kiribati | 1 | 0% | 0% | 3 | 0% | 0% |
| Poland | 1 | 0% | 0% | 3 | 0% | 0% |
| Togo | 1 | 0% | 0% | 3 | 0% | 0% |
| Saint Kitts and Nevis | 1 | 0% | 0% | 3 | 0% | 0% |
| United Arab Emirates | | | | 4 | 0% | 0% |
| El Salvador | 1 | 0% | 0% | 2 | 0% | 0% |
| Iceland | 1 | 0% | 0% | 2 | 0% | 0% |
| Kyrgyzstan | 1 | 0% | 0% | 2 | 0% | 0% |
| Lesotho | | | | 3 | 0% | 0% |
| North Korea | | | | 3 | 0% | 0% |
| Tajikistan | | | | 3 | 33% | 33% |
| Anguilla | 1 | 0% | 0% | 1 | 0% | 0% |
| Barbados | 1 | 0% | 0% | 1 | 0% | 0% |
| Denmark | 1 | 0% | 0% | 1 | 0% | 0% |
| Nauru | 1 | 0% | 0% | 1 | 0% | 0% |
| Tonga | 1 | 0% | 0% | 1 | 0% | 0% |
| Saint Barthelemy | 1 | 0% | 0% | 1 | 0% | 0% |
| Guinea-Bissau | 1 | 0% | 0% | 1 | 0% | 0% |
| Malta | 1 | 0% | 0% | 1 | 0% | 0% |
| Niue | 1 | 0% | 0% | 1 | 0% | 0% |
| Austria | 1 | 0% | 0% | 1 | 0% | 0% |
| Albania | | | | 2 | 0% | 0% |
| Ireland | | | | 2 | 50% | 50% |
| Uzbekistan | | | | 2 | 0% | 0% |
| Romania | | | | 2 | 0% | 0% |
| Finland | | | | 1 | 100% | 100% |
| Guam | | | | 1 | 0% | 0% |
| Gambia | | | | 1 | 0% | 0% |
| Croatia | | | | 1 | 0% | 0% |

| | | | |
|---|---|---|---|
| Jordan | 1 | 0% | 0% |
| Palestine | 1 | 0% | 0% |
| Burkina Faso | 1 | 100% | 0% |
| Bulgaria | 1 | 0% | 0% |
| Niger | 1 | 0% | 0% |

**SI Table 4. Overlap of critical natural assets with cultural diversity.**

| Country (ranked by # languages) | # Indigenous and Non-Migrant Languages | # languages intersecting critical natural assets for local NCP (LCNA) | # languages intersecting critical natural assets for global NCP (CGCNA) | % of languages on LCNA | % of languages on GCNA |
|---|---|---|---|---|---|
| Global | 6,474 | 6,220 | 5,976 | 96.1% | 92.3% |
| Papua New Guinea | 829 | 822 | 796 | 99.2% | 96.0% |
| Indonesia | 687 | 665 | 655 | 96.8% | 95.3% |
| Nigeria | 493 | 428 | 431 | 86.8% | 87.4% |
| India | 408 | 400 | 393 | 98.0% | 96.3% |
| Mexico | 277 | 271 | 234 | 97.8% | 84.5% |
| Cameroon | 267 | 256 | 252 | 95.9% | 94.4% |
| Democratic Republic of the Congo | 202 | 201 | 202 | 99.5% | 100.0% |
| China | 193 | 192 | 191 | 99.5% | 99.0% |
| Philippines | 171 | 167 | 157 | 97.7% | 91.8% |
| Brazil | 159 | 144 | 159 | 90.6% | 100.0% |
| Australia | 143 | 127 | 130 | 88.8% | 90.9% |
| Chad | 120 | 117 | 111 | 97.5% | 92.5% |
| Tanzania | 115 | 114 | 111 | 99.1% | 96.5% |
| United States | 109 | 100 | 92 | 91.7% | 84.4% |
| Malaysia | 109 | 105 | 105 | 96.3% | 96.3% |
| Myanmar | 109 | 106 | 109 | 97.2% | 100.0% |
| Vanuatu | 108 | 105 | 93 | 97.2% | 86.1% |
| Nepal | 104 | 102 | 102 | 98.1% | 98.1% |
| Viet Nam | 93 | 91 | 89 | 97.8% | 95.7% |
| Russian Federation | 91 | 88 | 86 | 96.7% | 94.5% |
| Peru | 88 | 82 | 87 | 93.2% | 98.9% |
| Ethiopia | 81 | 80 | 80 | 98.8% | 98.8% |
| Canada | 75 | 72 | 70 | 96.0% | 93.3% |
| Côte d'Ivoire | 74 | 72 | 70 | 97.3% | 94.6% |
| Colombia | 73 | 58 | 72 | 79.5% | 98.6% |
| Ghana | 71 | 67 | 65 | 94.4% | 91.5% |
| Solomon Islands | 69 | 68 | 57 | 98.6% | 82.6% |
| Sudan | 67 | 67 | 65 | 100.0% | 97.0% |
| Laos | 67 | 66 | 67 | 98.5% | 100.0% |
| Burkina Faso | 66 | 61 | 51 | 92.4% | 77.3% |
| Mali | 63 | 61 | 26 | 96.8% | 41.3% |
| Central African Republic | 62 | 62 | 62 | 100.0% | 100.0% |
| Pakistan | 58 | 52 | 46 | 89.7% | 79.3% |
| South Sudan | 57 | 57 | 57 | 100.0% | 100.0% |
| Kenya | 56 | 47 | 46 | 83.9% | 82.1% |
| Congo | 55 | 55 | 55 | 100.0% | 100.0% |
| Benin | 50 | 49 | 46 | 98.0% | 92.0% |
| Thailand | 48 | 44 | 45 | 91.7% | 93.8% |
| Iran | 43 | 35 | 30 | 81.4% | 69.8% |
| Togo | 40 | 36 | 34 | 90.0% | 85.0% |
| Angola | 40 | 40 | 40 | 100.0% | 100.0% |
| Gabon | 40 | 38 | 40 | 95.0% | 100.0% |
| Mozambique | 40 | 40 | 40 | 100.0% | 100.0% |
| Bangladesh | 36 | 36 | 33 | 100.0% | 91.7% |
| Uganda | 36 | 36 | 36 | 100.0% | 100.0% |
| Zambia | 35 | 35 | 35 | 100.0% | 100.0% |

| | | | | | |
|---|---|---|---|---|---|
| New Caledonia | 33 | 33 | 30 | 100.0% | 90.9% |
| Senegal | 31 | 31 | 25 | 100.0% | 80.6% |
| Guinea | 31 | 31 | 30 | 100.0% | 96.8% |
| Venezuela | 31 | 31 | 31 | 100.0% | 100.0% |
| Afghanistan | 30 | 28 | 28 | 93.3% | 93.3% |
| Bolivia | 30 | 29 | 30 | 96.7% | 100.0% |
| Liberia | 27 | 27 | 27 | 100.0% | 100.0% |
| Botswana | 26 | 24 | 26 | 92.3% | 100.0% |
| Guatemala | 23 | 23 | 23 | 100.0% | 100.0% |
| Italy | 21 | 20 | 19 | 95.2% | 90.5% |
| Bhutan | 21 | 21 | 21 | 100.0% | 100.0% |
| Namibia | 21 | 21 | 21 | 100.0% | 100.0% |
| Ecuador | 20 | 19 | 20 | 95.0% | 100.0% |
| Niger | 19 | 15 | 11 | 78.9% | 57.9% |
| Cambodia | 19 | 17 | 17 | 89.5% | 89.5% |
| East Timor | 19 | 18 | 18 | 94.7% | 94.7% |
| Guinea-Bissau | 18 | 18 | 17 | 100.0% | 94.4% |
| Paraguay | 18 | 18 | 18 | 100.0% | 100.0% |
| Sierra Leone | 18 | 18 | 18 | 100.0% | 100.0% |
| Micronesia | 17 | 17 | 0 | 100.0% | 0.0% |
| Georgia | 16 | 15 | 15 | 93.8% | 93.8% |
| Germany | 15 | 15 | 15 | 100.0% | 100.0% |
| Zimbabwe | 15 | 15 | 15 | 100.0% | 100.0% |
| Algeria | 13 | 7 | 5 | 53.8% | 38.5% |
| China-Taiwan | 13 | 12 | 12 | 92.3% | 92.3% |
| Argentina | 13 | 13 | 13 | 100.0% | 100.0% |
| Azerbaijan | 13 | 13 | 13 | 100.0% | 100.0% |
| France | 13 | 13 | 13 | 100.0% | 100.0% |
| Spain | 13 | 13 | 13 | 100.0% | 100.0% |
| Japan | 12 | 12 | 8 | 100.0% | 66.7% |
| Equatorial Guinea | 12 | 11 | 11 | 91.7% | 91.7% |
| Turkey | 12 | 11 | 11 | 91.7% | 91.7% |
| Malawi | 12 | 12 | 12 | 100.0% | 100.0% |
| Netherlands | 12 | 12 | 12 | 100.0% | 100.0% |
| South Africa | 12 | 12 | 12 | 100.0% | 100.0% |
| Iraq | 11 | 10 | 8 | 90.9% | 72.7% |
| Greece | 11 | 10 | 10 | 90.9% | 90.9% |
| Madagascar | 11 | 11 | 11 | 100.0% | 100.0% |
| Tajikistan | 11 | 11 | 11 | 100.0% | 100.0% |
| Belgium | 10 | 10 | 10 | 100.0% | 100.0% |
| Guyana | 10 | 9 | 10 | 90.0% | 100.0% |
| Suriname | 10 | 8 | 10 | 80.0% | 100.0% |
| Switzerland | 10 | 10 | 10 | 100.0% | 100.0% |
| Oman | 10 | 9 | 0 | 90.0% | 0.0% |
| Eritrea | 9 | 9 | 9 | 100.0% | 100.0% |
| Somalia | 9 | 9 | 9 | 100.0% | 100.0% |
| Fiji | 8 | 8 | 4 | 100.0% | 50.0% |
| Morocco | 8 | 7 | 5 | 87.5% | 62.5% |
| Finland | 8 | 8 | 8 | 100.0% | 100.0% |
| Sweden | 8 | 8 | 8 | 100.0% | 100.0% |
| Libya | 7 | 2 | 1 | 28.6% | 14.3% |
| Yemen | 7 | 7 | 3 | 100.0% | 42.9% |
| Gambia | 7 | 5 | 5 | 71.4% | 71.4% |

| | | | | | |
|---|---|---|---|---|---|
| Syria | 7 | 6 | 5 | 85.7% | 71.4% |
| Egypt | 7 | 6 | 6 | 85.7% | 85.7% |
| Austria | 7 | 7 | 7 | 100.0% | 100.0% |
| Brunei | 7 | 7 | 7 | 100.0% | 100.0% |
| French Guiana | 7 | 7 | 7 | 100.0% | 100.0% |
| Panama | 7 | 6 | 7 | 85.7% | 100.0% |
| Ukraine | 7 | 7 | 7 | 100.0% | 100.0% |
| French Polynesia | 7 | 5 | 0 | 71.4% | 0.0% |
| Israel | 5 | 4 | 2 | 80.0% | 40.0% |
| Chile | 5 | 4 | 4 | 80.0% | 80.0% |
| Mauritania | 5 | 5 | 4 | 100.0% | 80.0% |
| Portugal | 5 | 5 | 4 | 100.0% | 80.0% |
| Belize | 5 | 5 | 5 | 100.0% | 100.0% |
| Honduras | 5 | 5 | 5 | 100.0% | 100.0% |
| Mongolia | 5 | 5 | 5 | 100.0% | 100.0% |
| Montenegro | 5 | 5 | 5 | 100.0% | 100.0% |
| Nicaragua | 5 | 5 | 5 | 100.0% | 100.0% |
| North Macedonia | 5 | 5 | 5 | 100.0% | 100.0% |
| Serbia | 5 | 5 | 5 | 100.0% | 100.0% |
| Uzbekistan | 5 | 5 | 5 | 100.0% | 100.0% |
| Albania | 4 | 4 | 4 | 100.0% | 100.0% |
| Armenia | 4 | 4 | 4 | 100.0% | 100.0% |
| Costa Rica | 4 | 4 | 4 | 100.0% | 100.0% |
| Kazakhstan | 4 | 4 | 4 | 100.0% | 100.0% |
| Norway | 4 | 3 | 4 | 75.0% | 100.0% |
| Sri Lanka | 4 | 4 | 4 | 100.0% | 100.0% |
| United Kingdom | 4 | 4 | 4 | 100.0% | 100.0% |
| Cook Islands | 4 | 1 | 0 | 25.0% | 0.0% |
| Croatia | 3 | 3 | 2 | 100.0% | 66.7% |
| Jordan | 3 | 3 | 2 | 100.0% | 66.7% |
| Andorra | 3 | 3 | 3 | 100.0% | 100.0% |
| Bulgaria | 3 | 3 | 3 | 100.0% | 100.0% |
| Czechia | 3 | 3 | 3 | 100.0% | 100.0% |
| Hungary | 3 | 3 | 3 | 100.0% | 100.0% |
| Lesotho | 3 | 3 | 3 | 100.0% | 100.0% |
| Lithuania | 3 | 3 | 3 | 100.0% | 100.0% |
| Romania | 3 | 3 | 3 | 100.0% | 100.0% |
| Saudi Arabia | 3 | 3 | 3 | 100.0% | 100.0% |
| Slovakia | 3 | 3 | 3 | 100.0% | 100.0% |
| Comoros | 3 | 3 | 0 | 100.0% | 0.0% |
| Northern Mariana Islands | 3 | 2 | 0 | 66.7% | 0.0% |
| Palau | 3 | 3 | 0 | 100.0% | 0.0% |
| São Tomé and Príncipe | 3 | 3 | 0 | 100.0% | 0.0% |
| Cyprus | 2 | 2 | 1 | 100.0% | 50.0% |
| Djibouti | 2 | 2 | 1 | 100.0% | 50.0% |
| Belarus | 2 | 2 | 2 | 100.0% | 100.0% |
| China-Hong Kong | 2 | 2 | 2 | 100.0% | 100.0% |
| Denmark | 2 | 2 | 2 | 100.0% | 100.0% |
| Estonia | 2 | 2 | 2 | 100.0% | 100.0% |
| Grenada | 2 | 2 | 2 | 100.0% | 100.0% |
| Ireland | 2 | 2 | 2 | 100.0% | 100.0% |
| Jamaica | 2 | 2 | 2 | 100.0% | 100.0% |
| Kyrgyzstan | 2 | 2 | 2 | 100.0% | 100.0% |

| | | | | | |
|---|---|---|---|---|---|
| Latvia | 2 | 2 | 2 | 100.0% | 100.0% |
| Liechtenstein | 2 | 2 | 2 | 100.0% | 100.0% |
| Luxembourg | 2 | 2 | 2 | 100.0% | 100.0% |
| Moldova | 2 | 2 | 2 | 100.0% | 100.0% |
| Palestine | 2 | 2 | 2 | 100.0% | 100.0% |
| Poland | 2 | 2 | 2 | 100.0% | 100.0% |
| South Korea | 2 | 2 | 2 | 100.0% | 100.0% |
| Trinidad and Tobago | 2 | 2 | 2 | 100.0% | 100.0% |
| Turkmenistan | 2 | 2 | 2 | 100.0% | 100.0% |
| Mayotte | 2 | 2 | 0 | 100.0% | 0.0% |
| Tonga | 2 | 2 | 0 | 100.0% | 0.0% |
| Wallis and Futuna | 2 | 2 | 0 | 100.0% | 0.0% |
| Western Sahara | 2 | 2 | 0 | 100.0% | 0.0% |
| Aland Islands | 1 | 1 | 1 | 100.0% | 100.0% |
| Antigua and Barbuda | 1 | 1 | 1 | 100.0% | 100.0% |
| Bahamas | 1 | 1 | 1 | 100.0% | 100.0% |
| Barbados | 1 | 1 | 1 | 100.0% | 100.0% |
| Bosnia and Herzegovina | 1 | 1 | 1 | 100.0% | 100.0% |
| British Virgin Islands | 1 | 1 | 1 | 100.0% | 100.0% |
| Burundi | 1 | 1 | 1 | 100.0% | 100.0% |
| China-Macao | 1 | 1 | 1 | 100.0% | 100.0% |
| Curaçao | 1 | 1 | 1 | 100.0% | 100.0% |
| Dominica | 1 | 1 | 1 | 100.0% | 100.0% |
| El Salvador | 1 | 1 | 1 | 100.0% | 100.0% |
| Eswatini | 1 | 1 | 1 | 100.0% | 100.0% |
| Guadeloupe | 1 | 1 | 1 | 100.0% | 100.0% |
| Haiti | 1 | 1 | 1 | 100.0% | 100.0% |
| Iceland | 1 | 1 | 1 | 100.0% | 100.0% |
| Isle of Man | 1 | 1 | 1 | 100.0% | 100.0% |
| Jersey | 1 | 1 | 1 | 100.0% | 100.0% |
| Kuwait | 1 | 1 | 1 | 100.0% | 100.0% |
| Lebanon | 1 | 1 | 1 | 100.0% | 100.0% |
| Malta | 1 | 1 | 1 | 100.0% | 100.0% |
| Martinique | 1 | 1 | 1 | 100.0% | 100.0% |
| Montserrat | 1 | 1 | 1 | 100.0% | 100.0% |
| New Zealand | 1 | 1 | 1 | 100.0% | 100.0% |
| North Korea | 1 | 1 | 1 | 100.0% | 100.0% |
| Rwanda | 1 | 1 | 1 | 100.0% | 100.0% |
| Saint Kitts and Nevis | 1 | 1 | 1 | 100.0% | 100.0% |
| Saint Lucia | 1 | 1 | 1 | 100.0% | 100.0% |
| Saint Martin | 1 | 1 | 1 | 100.0% | 100.0% |
| Saint Vincent and the Grenadines | 1 | 1 | 1 | 100.0% | 100.0% |
| Slovenia | 1 | 1 | 1 | 100.0% | 100.0% |
| Tunisia | 1 | 1 | 1 | 100.0% | 100.0% |
| Turks and Caicos Islands | 1 | 1 | 1 | 100.0% | 100.0% |
| U.S. Virgin Islands | 1 | 1 | 1 | 100.0% | 100.0% |
| American Samoa | 1 | 1 | 0 | 100.0% | 0.0% |
| Anguilla | 1 | 1 | 0 | 100.0% | 0.0% |
| Aruba | 1 | 1 | 0 | 100.0% | 0.0% |
| Bahrain | 1 | 1 | 0 | 100.0% | 0.0% |
| Cape Verde Islands | 1 | 1 | 0 | 100.0% | 0.0% |
| Cocos (Keeling) Islands | 1 | 1 | 0 | 100.0% | 0.0% |
| Faroe Islands | 1 | 1 | 0 | 100.0% | 0.0% |

| | | | | | |
|---|---|---|---|---|---|
| Gibraltar | 1 | 1 | 0 | 100.0% | 0.0% |
| Greenland | 1 | 1 | 0 | 100.0% | 0.0% |
| Guam | 1 | 1 | 0 | 100.0% | 0.0% |
| Kiribati | 1 | 1 | 0 | 100.0% | 0.0% |
| Maldives | 1 | 1 | 0 | 100.0% | 0.0% |
| Marshall Islands | 1 | 1 | 0 | 100.0% | 0.0% |
| Mauritius | 1 | 1 | 0 | 100.0% | 0.0% |
| Monaco | 1 | 1 | 0 | 100.0% | 0.0% |
| Nauru | 1 | 1 | 0 | 100.0% | 0.0% |
| Niue | 1 | 1 | 0 | 100.0% | 0.0% |
| Norfolk Island | 1 | 1 | 0 | 100.0% | 0.0% |
| Qatar | 1 | 1 | 0 | 100.0% | 0.0% |
| RÃ©union | 1 | 1 | 0 | 100.0% | 0.0% |
| Samoa | 1 | 1 | 0 | 100.0% | 0.0% |
| San Marino | 1 | 0 | 0 | 0.0% | 0.0% |
| Seychelles | 1 | 1 | 0 | 100.0% | 0.0% |
| Singapore | 1 | 1 | 0 | 100.0% | 0.0% |
| Tokelau | 1 | 1 | 0 | 100.0% | 0.0% |
| Tuvalu | 1 | 1 | 0 | 100.0% | 0.0% |
| United Arab Emirates | 1 | 1 | 0 | 100.0% | 0.0% |
| Grand Total [1] | 7,410 | | | | |

1: Total here does not equal total languages intersected for entire dataset as some languages occur in multiple countries.

Total Languages: 7,117

Total Indigenous Languages defined by Polygons: 6,474

Language Dataset: *Ethnologue* , 23rd edition
